# Parallel evolution of UbiA superfamily proteins into aromatic *O*-prenyltransferases in plants

**DOI:** 10.1101/2020.07.07.192757

**Authors:** Ryosuke Munakata, Alexandre Olry, Tomoya Takemura, Kanade Tatsumi, Takuji Ichino, Cloé Villard, Joji Kageyama, Tetsuya Kurata, Masaru Nakayasu, Florence Jacob, Takao Koeduka, Hirobumi Yamamoto, Eiko Moriyoshi, Tetsuya Matsukawa, Jeremy Grosjean, Célia Krieger, Akifumi Sugiyama, Masaharu Mizutani, Frédéric Bourgaud, Alain Hehn, Kazufumi Yazaki

**Author notes:** **Author for correspondence** Kazufumi Yazaki, Alain Hehn.

## Abstract

Plants produce approximately 300 aromatic molecules enzymatically linked to prenyl side chains via C-O bonds. These *O*-prenylated aromatics have been found in taxonomically distant plant taxa as compounds beneficial or detrimental to human health, with *O*-prenyl moieties often playing crucial roles in their biological activities. To date, however, no plant gene encoding an aromatic *O*-prenyltransferase (*O*-PT) has been described. This study describes the isolation of an aromatic *O*-PT gene, *CpPT1*, belonging to the UbiA superfamily, from grapefruit (*Citrus* × *paradisi,* Rutaceae). This gene is responsible for the biosynthesis of *O*-prenylated coumarin derivatives that alter drug pharmacokinetics in the human body. Another coumarin *O*-PT gene of the same protein family was identified in *Angelica keiskei*, an apiaceous medicinal plant containing pharmaceutically active *O*-prenylated coumarins. Phylogenetic analysis of these *O*-PTs suggested that aromatic *O*-prenylation activity evolved independently from the same ancestral gene in these distant plant taxa. These findings shed light on understanding the evolution of plant secondary metabolites via the UbiA superfamily.

## Introduction

Plants produce many *O*-prenylated aromatic molecules possessing prenyl side chains attached to the aromatic cores *via* C-O bonds. These aromatic core structures include flavonoids, coumarins, xanthones, and aromatic alkaloids, with roughly half of them (ca. 150 structures) being classified as coumarins^1, 2^. Some *O*-prenylated aromatics have pharmaceutical activities, whereas others are deleterious to human health^1, 2^. These beneficial/detrimental activities are often due to or enhanced by *O*-prenyl moieties^3–6^.

Native coumarin *O*-prenyltransferases (*O*-PT) of Rutaceae and Apiaceae, plants that accumulate large amounts of *O*-prenylated coumarins, have been characterized biochemically, with members of the membrane-bound UbiA superfamily of proteins found to be involved in coumarin *O*-prenylation^7, 8^. To date, approximately 50 UbiA superfamily genes have been found to encode aromatic *C*-PTs, which transfer prenyl moieties to aromatic cores *via* C-C bonds. Although these genes were shown to encode enzymes involved in plant primary and secondary metabolism, no gene encoding an aromatic *O*-PT has yet been identified in plants. *O*-Prenylated aromatic compounds have been detected in several distant plant families, including Asteraceae, Boraginaceae, Fabaceae, Hypericaceae, Rutaceae and Apiaceae, but are not ubiquitous throughout the plant kingdom^1^. The lack of knowledge of aromatic *O*-PT genes has prevented a determination of the appearance of aromatic *O*-prenylation activity during plant speciation.

Among Rutaceae, *Citrus* species accumulate large amounts of *O*-prenylated coumarins, especially in their flavedo (outer pericarp)^9–12^. Citrus *O*-prenylated coumarins have shown various pharmaceutical properties^2^, including anti-cancer^4, 13^, anti-microbial^3^, and anti-inflammatory^14^ activities, although some of these derivatives have shown undesirable effects in humans. Citrus fruits and juices enhance the bioavailability of orally administrated medications, which can lead to overdoses and increased side effects^5, 15^. The ‘grapefruit-drug interactions’ have been found to alter the pharmacokinetics of more than 85 medications, including statins and calcium channel blockers^15^. The United States Food and Drug Administration has cautioned consumers not to consume grapefruits or grapefruit juice at times close to taking such drugs^16^. Citrus species are thought to alter drug pharmacokinetics by inactivating CYP3A4, the major xenobiotic-metabolizing enzyme in the intestines and liver^5, 15^.

Furanocoumarins (FCs) are tricyclic coumarins containing a furan ring. *O*-geranylated forms of FCs including bergamottin and its oxidative derivatives (Fig. 1a) are promising candidates causing grapefruit-drug interactions due to their potent inhibition of CYP3A4^5, 15, 17^. CYP3A4-catalyzed metabolism of their furan rings produces reactive chemicals that inactivate this enzyme itself^18^, while their *O*-geranyl side chains contribute to binding to the active site of CYP3A4^6^ toward metabolization of the furan ring. Bergamottin and 6’,7’-dihydroxybergamottin show 7- and 160-fold higher *in vitro* inhibitory activity, respectively, than the non-geranylated form, bergaptol^5^. Furthermore, *O*-geranyl moieties act as linkers to form FC dimers, called paradisins, which are more potent CYP3A4 inactivators than monomeric *O*-geranylated FCs^5^. These findings suggest that paradisins, along with *O*-geranylated FC monomers, may be involved in grapefruit-drug interactions.

**Fig. 1.**
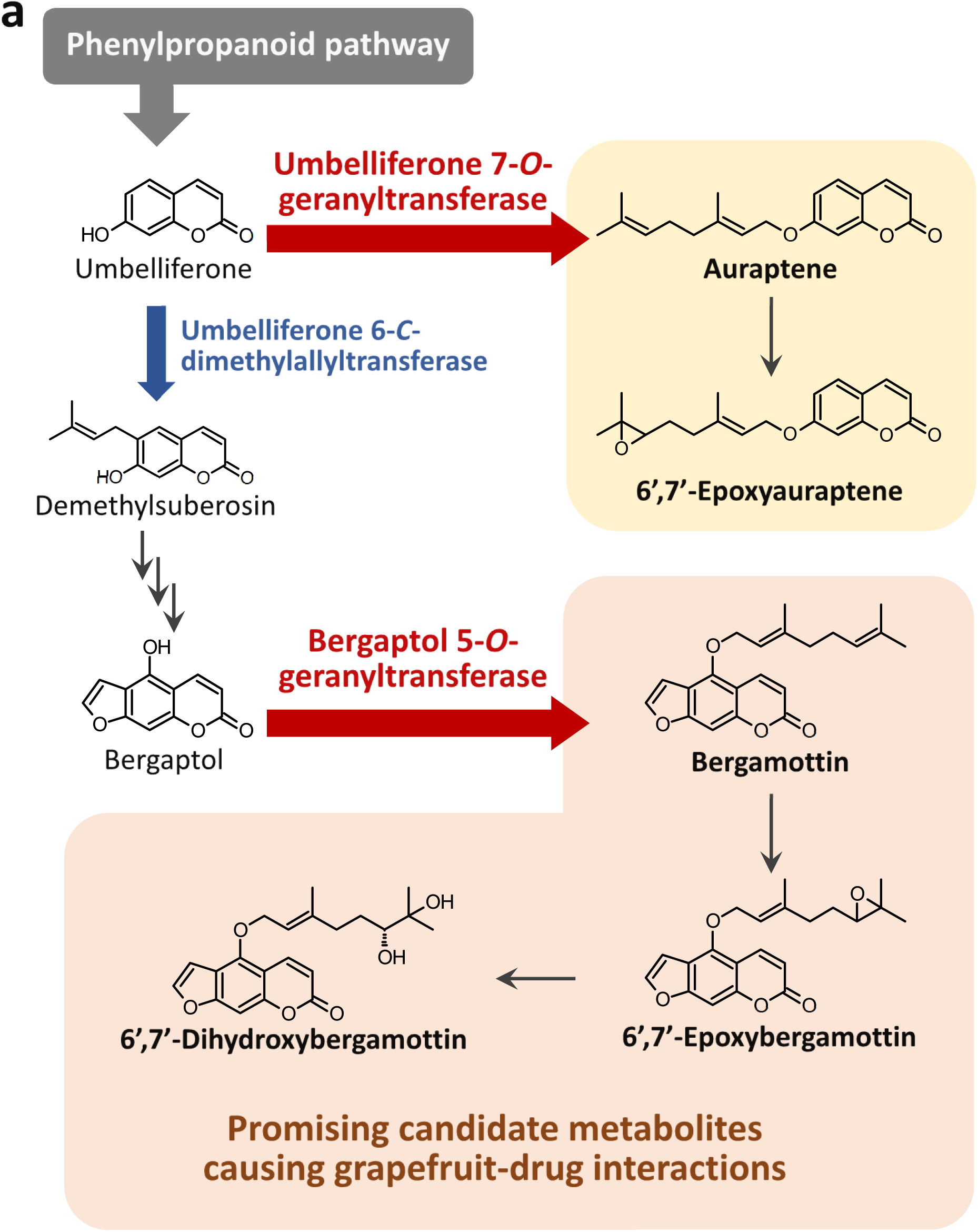
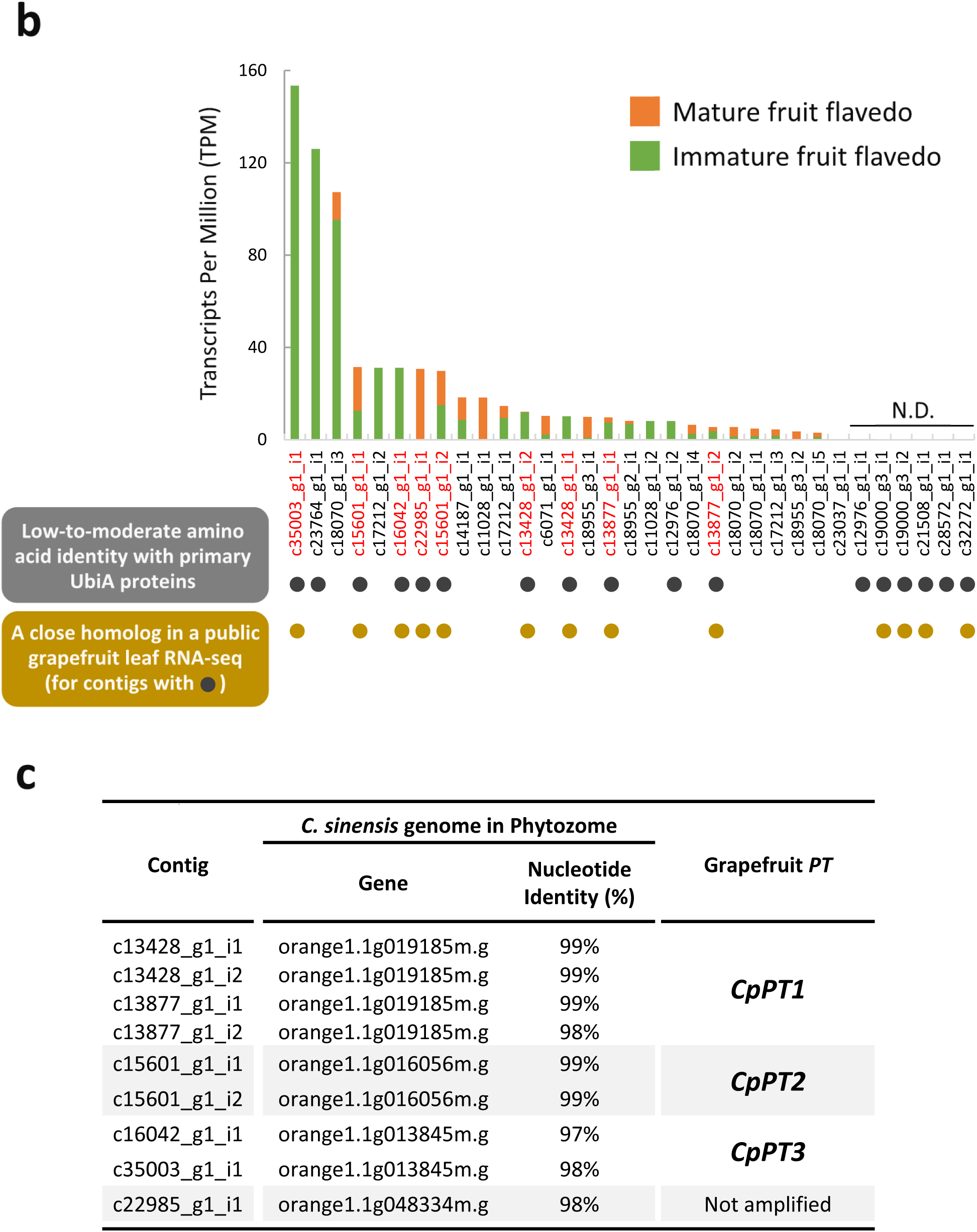
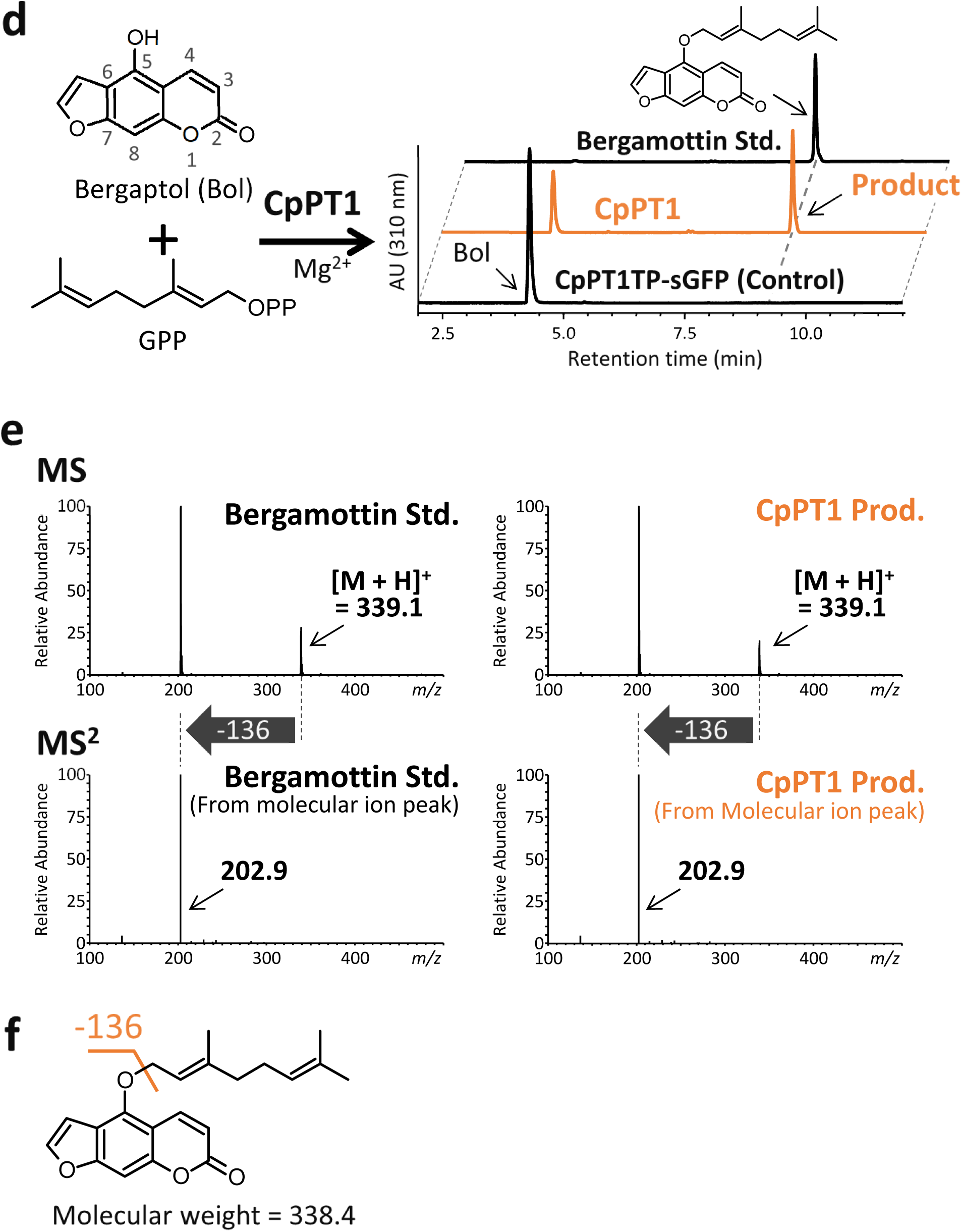
Isolation of a bergaptol 5-*O*-geranyltransferase gene from grapefruit. **a** Biosynthetic pathway of the major *O*-prenylated aromatic compounds in grapefruit. Biosynthetic steps catalyzed by *O*-prenyltransferases (PTs) and a *C*-PT are shown in red and blue, respectively. Metabolites derived from umbelliferone 7-*O*-geranyltransferase (GT) and bergaptol 5-*O*-GT are highlighted in yellow and orange, respectively. Bergamottin and its downstream metabolites are considered promising candidates responsible for grapefruit-drug interactions. **b** *In silico* search for aromatic *O*-PT candidates in a transcriptome dataset constructed from immature and mature grapefruit flavedo tissues. Contigs annotated as UbiA superfamily genes were rated based on (1) low-to-moderate amino acid identities between their encoded proteins and primary metabolism-related UbiA PTs (gray circles), (2) the presence in a public transcript dataset from grapefruit leaves of close homologs of contigs meeting the first criterion (yellow circles), and (3) transcripts per million (TPM)-based expression of grapefruit flavedo contigs to remove contigs with undetectable expression. The contigs meeting these three criteria are highlighted in red. For detailed information about screening for criteria (1) and (2), please see Supplementary Table 1 and 2, respectively. N.D., not detected. **c** Mapping of the nine candidates from **(a)** onto the sweet orange genome by blastn search in Phytozome. **d** HPLC analysis of a B5OGT reaction mixture of recombinant CpPT1. Microsomes prepared from *N. benthamiana* leaves expressing *CpPT1* were used as crude enzymes, with the negative control being microsomes prepared from *N. benthamiana* leaves expressing a chimeric protein consisting of N-terminal amino acids 1–70 of CpPT1, including the transit peptide (TP), and synthetic green fluorescence protein (CpPT1TP-sGFP). UV chromatograms of the full assay and the negative control assay at 310 nm are shown at a comparable scale. **e** MS^2^ analysis of the enzymatic product in the positive ion mode. A loss of 136 mass units was predicted to correspond to fragmentation resulting from the loss of the entire *O*-geranyl moiety attached to the aromatic ring. **f** Proposed MS^2^ fragmentation of bergamottin.

Starting with transcriptome analysis of flavedo tissues, this study describes the isolation of a gene encoding a coumarin *O*-PT involved in bergamottin biosynthesis in grapefruit and the functional characterization of its gene product. The contribution of *O*-PT orthologs to coumarin biosynthesis was assessed in various *Citrus* species. In addition, an aromatic *O*-PT was isolated from *Angelica keiskei*, an apiaceous medicinal plant producing *O*-prenylated coumarins^19^. The evolutionary development of aromatic *O*-prenylation activity in plants was assessed by phylogenetic analysis of *O*-PTs from taxonomically distant families Rutaceae and Apiaceae.

## Results

### Construction of a transcriptome dataset from grapefruit flavedo tissues

Because the native enzymes catalyzing coumarin *O*-prenylation in lemon flavedo were shown to possess characteristics common to PTs in the UbiA superfamily^8^, the genomes of *C. sinensis* (sweet orange, the male parent of grapefruit) and *C. clementina* (clementine) in the public Phytozome database were searched to identify genes in this family. The search term “UbiA” identified 26 and 27 loci in the sweet orange and clementine genomes, respectively. In contrast, search of the *Arabidopsis thaliana* genome identified only six loci, which formed a minimal gene set only for the six primary metabolic pathways relevant to the UbiA superfamily^20^. These *in silico* searches identified members of the UbiA superfamily potentially involved in the specialized metabolism of *Citrus* genus, as exemplified by the synthesis of 8-*C*-geranylumbelliferone by a lemon UbiA *C-*PT, ClPT1^21^. To better identify aromatic *O*-PT candidates, we performed transcriptome analysis of grapefruit, which is rich in *O*-prenylated coumarins and is representative of grapefruit-drug interactions.

Grapefruit primarily accumulates two types of *O*-prenylated phenolics, auraptene- and bergamottin-related compounds, which are likely synthesized by distinct *O*-geranylation pathways, catalyzed by umbelliferone 7-*O*-geranyltransferase (U7OGT) and bergaptol 5-*O*-GT (B5OGT), respectively (Fig. 1a)^9, 10^. Quantification of these major *O*-prenylated coumarins in different grapefruit organs revealed that they are most abundant in flavedo tissues of immature and mature fruits (Supplementary Figs. 1 and 2), from which we constructed a transcriptome dataset.

### Isolation of candidate genes encoding *O*-PTs from grapefruit

To comprehensively identify UbiA PTs involved in plant specialized metabolism in the grapefruit flavedo transcriptome^10, 30^, *in silico* screening was performed using seven query sequences (Supplementary Table 1), *i.e.*, ClPT1^21^ and six sweet orange proteins probably orthologous to *Arabidopsis thaliana* UbiA PTs functionally involved in primary metabolic pathways^20^. Candidates for coumarin *O*-PTs were selected based on three criteria: (1) low-to-moderate amino acid identity to UbiA proteins in plant primary metabolism (Fig. 1b and Supplementary Table 1); (2) presence in another public transcriptome dataset prepared from grapefruit leaves that accumulate *O*-geranylated coumarins (Fig. 1b and Supplementary Table 2); and (3) transcripts per million (TPM)-based expression levels of grapefruit flavedo contigs to remove those with zero TPM (Fig. 1b). Nine contigs, mapped to four genes in the sweet orange genome, were selected (Fig. 1c).

Using RT-PCR in reference to corresponding sweet orange sequences, we isolated the full coding sequences (CDSs) of three candidate genes from grapefruit flavedo, naming them *C.* × *paradisi PT 1–3* (*CpPT1–3*), respectively (Fig. 1c). We failed to amplify a CDS for the other candidate gene. However, the transcript corresponding to c22985_g1_i1 seems to be nonfunctional, due to the lack of a coding region containing the second aspartate-rich motif that is essential for prenylation reactions in UbiA proteins^24, 25^. *In silico* analysis predicted that the polypeptides CpPT1 and CpPT2 each include two aspartate-rich motifs, multiple transmembrane regions, and transit peptides (TPs), all of which are characteristics of plant UbiA PT proteins^20–23^ (Supplementary Fig. 3). Although CpPT3 was not predicted to have a TP, its score was just below the threshold for the detection of a TP.

### Functional screening of CpPTs

These individual PTs were transiently expressed in *Nicotiana benthamiana* leaves by agroinfiltration. Microsomes prepared from these leaves were subjected to B5OGT and U7OGT assays in the presence of MgCl_2_ as a cofactor. Neither CpPT2 nor CpPT3 was able to synthesize any *O*-geranylated products in B5OGT assays, in which bergaptol was used as the prenyl acceptor substrate (Supplementary Fig. 4a and b). CpPT2 was also unable to synthesize any product in U7OGT assays, in which umbelliferone was the aromatic substrate, whereas CpPT3 catalyzed the production of two products, 8-*C*-geranylumbelliferone and a by-product, but not auraptene (Supplementary Fig. 4). Because CpPT3 and ClPT1 had the same enzymatic properties and were highly (95%) homologous (Supplementary Fig. 3a)^21^, we concluded that these two enzymes are orthologous to each other. CpPT2 and CpPT3 were also incubated in the presence of various substrate pairs, but no clear *O*-prenylation activity was detected (Supplementary Fig. 4a and b).

Although CpPT1-expressing microsomes did not yield any products in U7OGT assays, HPLC analysis showed that these microsomes generated a product in B5OGT assays (Fig. 1d). This product had the identical retention time and MS and MS^2^ spectra as bergamottin, a finding confirmed by direct comparison with a standard specimen (Fig. 1e). In MS^2^ analysis using the positive ion mode, the major peak after fragmentation of the molecular ion of bergamottin (*m/z* = 339) was at *m/z* = 203, with the difference of 136 mass units corresponding to the molecular weight of a geranyl chain (Fig. 1f). This total loss of a prenyl moiety is possibly unique to *O*-prenylated aromatics, as one carbon at the benzyl position is left after the fragmentation of *C*-prenyl moieties^26^, resulting in a loss of 124 mass units for *C*-geranyl moieties^21, 26^. These biochemical findings suggested that CpPT1 is a strong B5OGT candidate.

### Enzymatic properties of CpPT1

The specificity of CpPT1 for coumarin molecules as prenyl acceptors was analyzed in the presence of the prenyl donor geranyl diphosphate (GPP; Table 1 and Supplementary Fig. 5a). CpPT1 was able to transfer prenyl moieties to coumarin molecules with hydroxy groups at the C5 (No. 3 and 7) and C8 (No. 13 and 16) positions as aromatic substrates. These enzymatic products were identified as *O*-geranylated forms by direct comparison with available standards (Supplementary Fig. 5b–e) and predicted by their MS^2^ fragmentation patterns if standards were unavailable (Supplementary Fig. 5f– i). Coumarin and FC derivatives without a hydroxy group at C5 or C8, as well as molecules in other phenolic classes (No. 19–25), were not recognized as substrates.

**Table 1.**
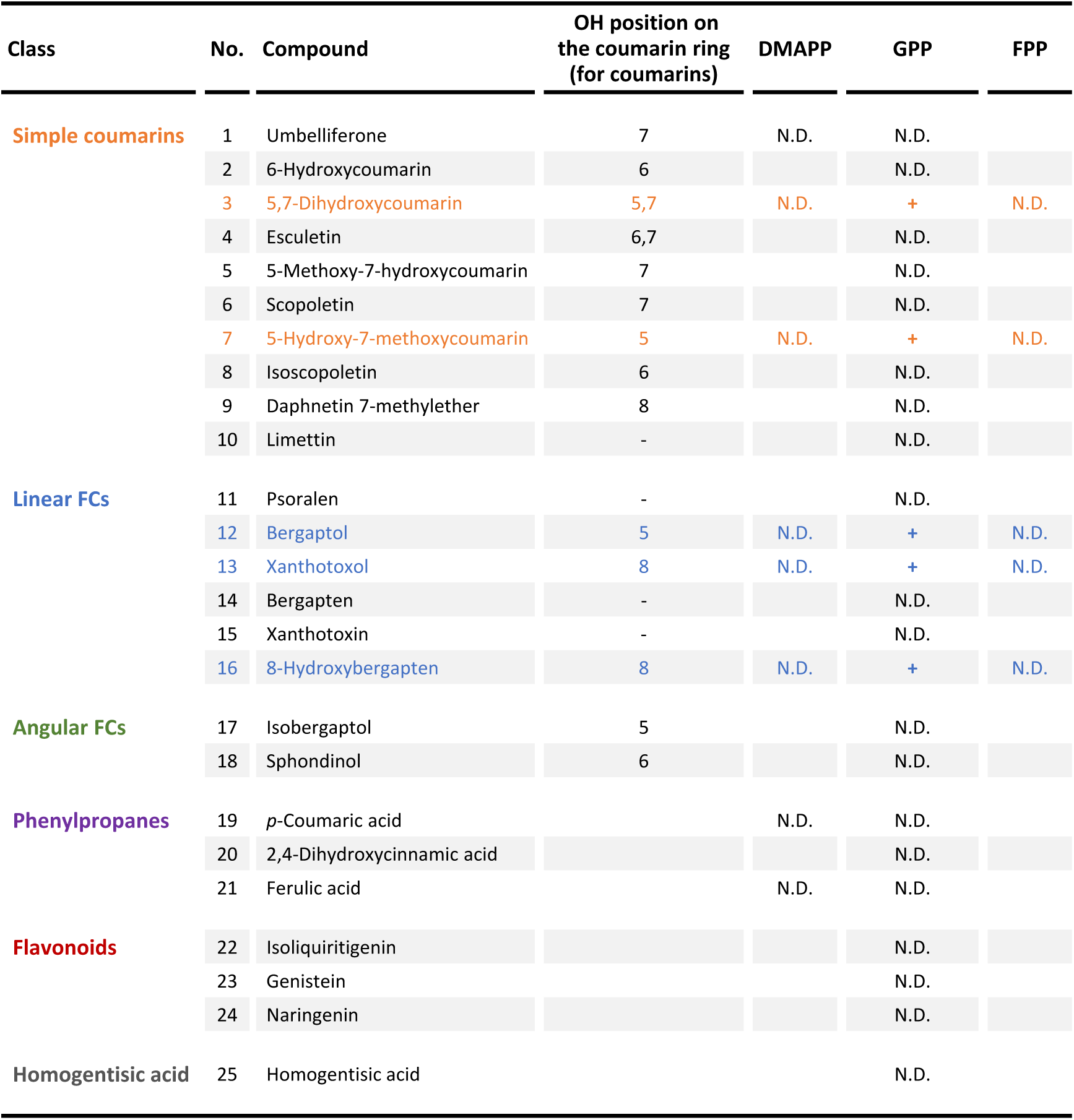
Substrate specificity of CpPT1. Simple coumarins (No. 1–10), linear FCs (11–16), angular FCs (17 and 18), phenylpropanes (19– 21), flavonoids (22–24) and homogentisic acid (25) were tested as possible prenyl acceptor substrates of CpPT1. Dimethylallyl diphosphate (DMAPP), geranyl diphosphate (GPP), and farnesyl diphosphate (FPP) were tested as possible prenyl donor substrates. Independent triplicate reactions gave the same result. The results of the aromatic substrates accepted by CpPT1 are highlighted in color. The substrate pairs resulting in enzymatic products are marked with pluses. N.D., not detected. The chemical structures of the aromatic substrates are shown in Supplementary Fig. 5a (all molecules) and Table 2 (molecules accepted by CpPT1).

The prenyl donor specificity of CpPT1 was investigated in parallel using dimethylallyl diphosphate (DMAPP) and farnesyl diphosphate (FPP), employing coumarin derivatives accepted in GT assays, but no reaction products were observed for any combination (Table 1). Umbelliferone, *p*-coumaric acid, and ferulic acid were also tested in dimethylallyltransferase (DT) assays because their dimethylallylated forms have been found in Rutaceae species with *C-*dimethylallylated umbelliferone molecules being precursors of FCs (Fig. 1a)^27, 28^. However, no reaction products were observed (Table 1). Taken together, these biochemical analyses demonstrated that CpPT1 functions as a coumarin *5/8-O*-GT.

Kinetic analysis of the *O*-geranylation activity of CpPT1 in the presence of the five coumarin substrates demonstrated that bergaptol was the optimal prenyl acceptor (Table 2a). Kinetic analysis for GPP measured in the presence of bergaptol or its structural isomer, xanthtotoxol, resulted in similar apparent *K*_m_ values, irrespective of prenyl acceptor substrates (Table 2b). These results indicated that the recombinant CpPT1 mainly functions as B5OGT, with an optimal pH in the neutral-to-weak-alkaline region (Supplementary Fig. 6a). Analysis of its divalent cation preference showed that CpPT1 recognized Mg^2+^ as its best cofactor (Supplementary Fig. 6b). The B5OGT enzymatic activities of recombinant CpPT1 and the native microsomes prepared from lemon flavedo were similar^8^.

**Table 2.**
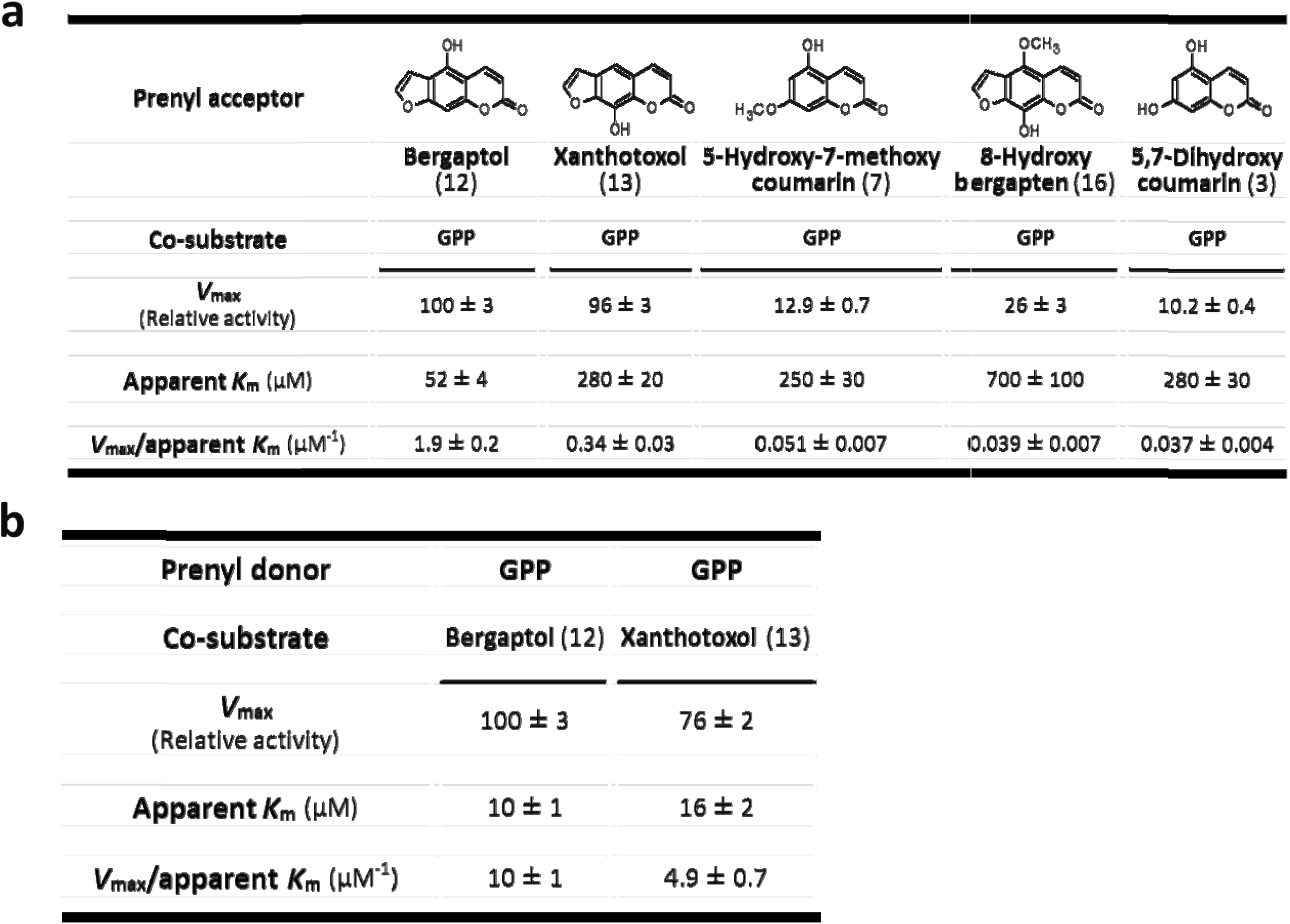
Kinetics of geranyltransferase activities of CpPT1. Kinetic analysis of *O*-geranylation activity of CpPT1 for coumarin molecules (**a**) and geranyl diphosphate (GPP) (**b**). *V*_max_ values are shown relative to B5OGT activity. The *O*-geranylated products of 5-hydroxy-7-methoxycoumarin (5H7M) and bergaptol were quantified by comparison with standard specimens of 5-geranyloxy-7-methoxycoumarin and bergamottin, respectively. The enzymatic products of 5,7-dihydroxycoumarin, xanthotoxol, and 8-hydroxybergapten were quantified as equivalents to 5,7-dihydroxycoumarin, imperatorin (*O-* dimethylallylated xanthotoxol), and phellopterin (*O-*dimethylallylated 8-hydroxybergapten), respectively, due to unavailability or instability of their *O*-geranylated compounds. All results are expressed as means ± standard errors of three independent experiments. The numerical codes for prenyl acceptor molecules are linked to those in Table 1.

### *In planta* gene expression profile and subcellular localization of CpPT1

To assess the involvement of *CpPT1* in the biosynthesis of *O*-prenylated coumarins in grapefruit, the levels of expression of *CpPT1* were determined in different organs. Assessment of both immature and mature fruits showed that *CpPT1* was highly expressed in flavedo but weakly expressed in albedo (Fig. 2a). This gene is also expressed in buds and leaves at similar-to-lower levels than in flavedo tissues (Fig. 2a). This expression profile fits with the accumulation patterns of bergamottin and its downstream derivatives (Supplementary Fig. 2).

**Fig. 2.**
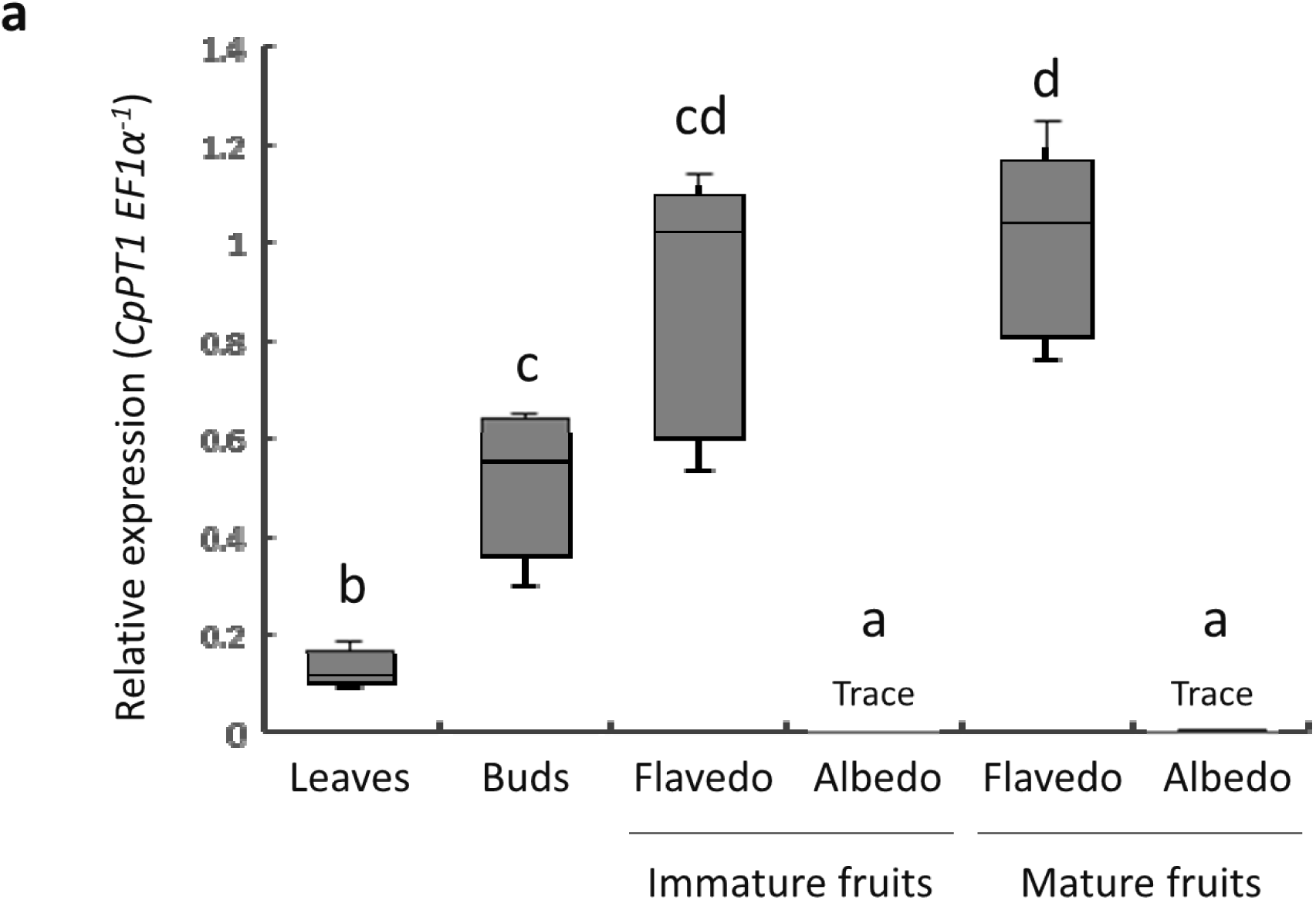
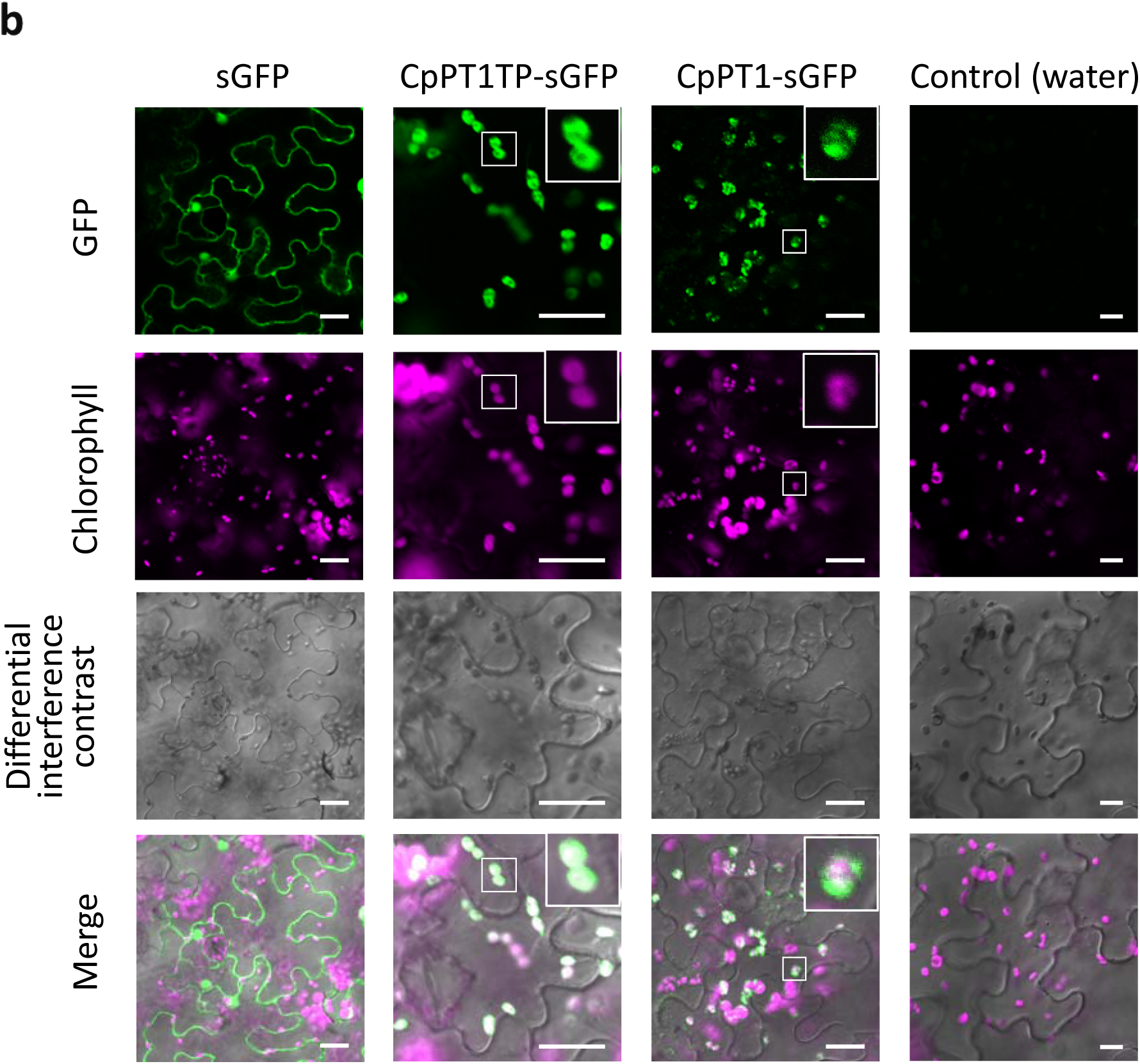
Organ-specific gene expression and subcellular localization of CpPT1. **a** Organ-specific expression of *CpPT1*. Ratios of the relative expression of *CpPT1* to *CpEF1α* in grapefruit leaves, buds, and the flavedo and albedo of immature and mature fruits, normalized to the average *CpPT1/CpEF1α* ratio in mature flavedo (n = 5 biological replicates). Relative levels of expression are shown as box plots (center line, median; box limits, first and third quartiles; whiskers, minimum and maximum). Significant differences between groups are indicated by letters (*p* < 0.05 by Games-Howell tests). **b** Subcellular localization of CpPT1TP-sGFP and CpPT1-sGFP. Free sGFP, CpPT1TP-sGFP, and CpPT1-sGFP were transiently expressed in *N. benthamiana* leaves by agroinfiltration, with the negative control consisting of leaves infiltrated by water. For merging, the brightness and contrast of the fluorescent images were adjusted in an unbiased manner, with magenta being a pseudo-color for chlorophyll autofluorescence signal. Enlarged images are inserted for CpPT1TP-sGFP and CpPT1-sGFP. Scale bars indicate 20 µm.

To assess the subcellular localization of CpPT1 *in planta*, synthetic GFP (sGFP) was fused to the C-terminus of the first 70 amino acids containing the predicted TP of CpPT1 (CpPT1TP-sGFP) or to the C-terminus of the full-length polypeptide (CpPT1-sGFP) (Supplementary Fig. 3b). Confocal microscopy of epidermal cells of *N. benthamiana* leaves expressing these GFP-fusion proteins indicated that both chimeric proteins localize to chloroplasts (Fig. 2b). These results strongly suggest that CpPT1 functions in plastids in grapefruit, consistent with the plastid localization of the MEP pathway that provides GPP in plant cells.

### FC chemotypes related to the gene structures of *CpPT1* orthologs in *Citrus*

*Citrus* domestication involved crossing of the four ancestral species, citron (*C. medica*), pure (or ancestral) mandarin (*C. reticulata*), papeda (*C. micrantha*), and pummelo (*C. grandis*), among themselves and/or with their descendants, generating most of the currently cultivated varieties, such as sweet orange, lemon, and grapefruit^29–31^. The concentrations and composition of coumarins in the flavedo of these species vary, with papeda, pummelo and citron varieties producing high quantities, and mandarin varieties producing low quantities, of coumarins (Fig. 3a)^10^. Interestingly, the major *O*-geranylated FC in citrus, bergamottin, and its related metabolites are undetectable in citron varieties, despite their high contents of FCs (Fig. 3a and b)^10^.

**Fig. 3.**
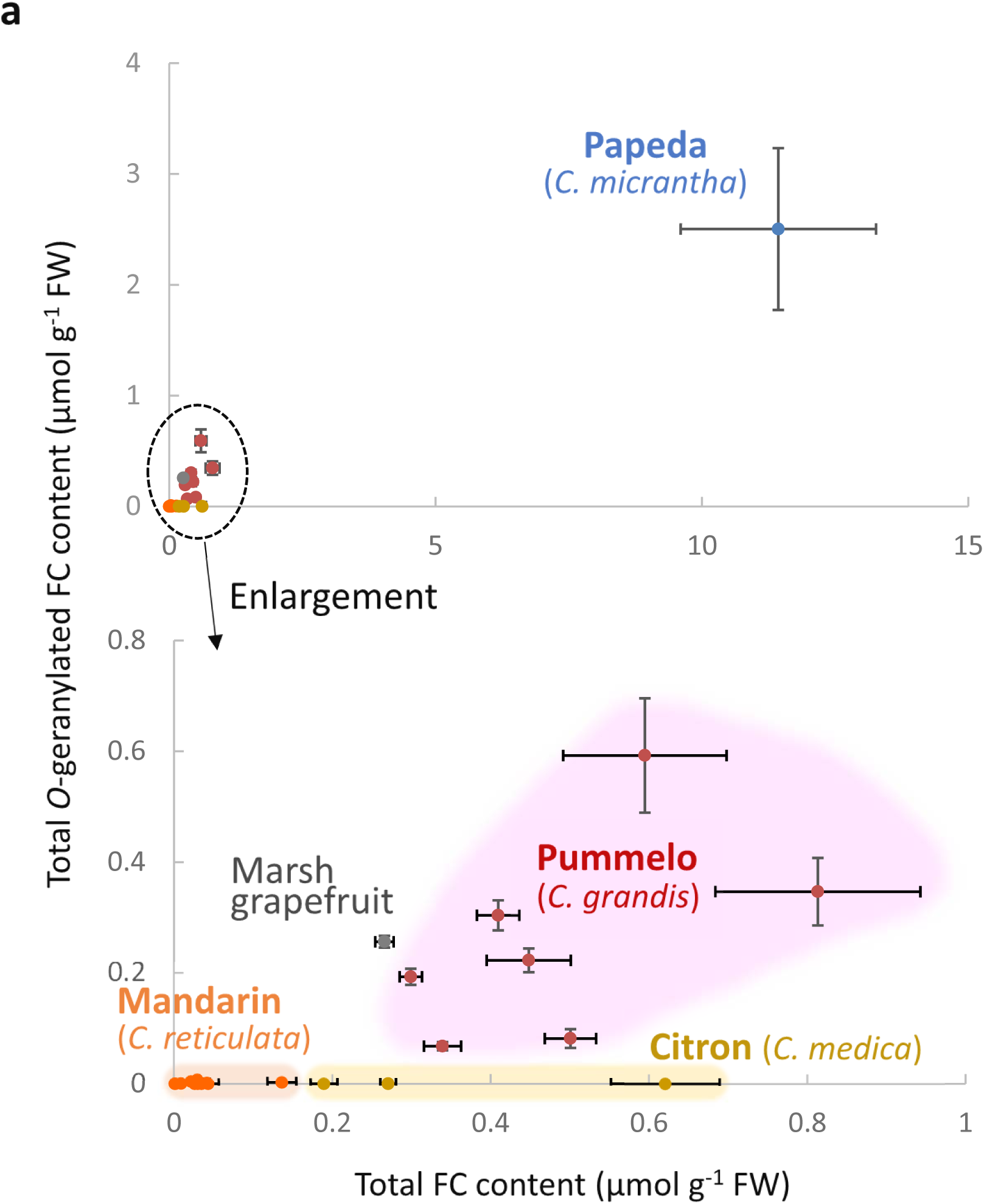
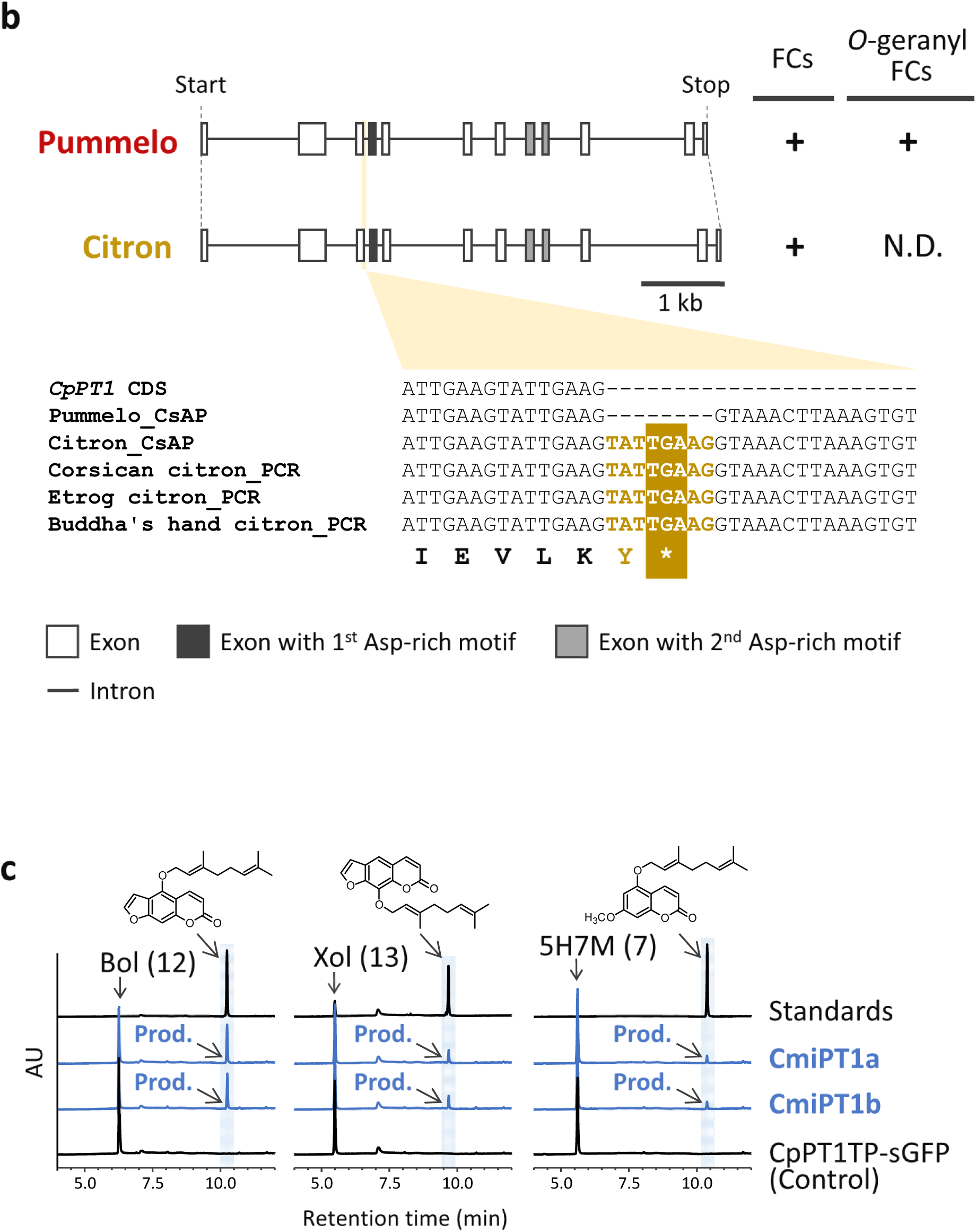
Conservation of *CpPT1* orthologs in *Citrus* genus. **a** Total FC and *O*-geranylated FC contents in flavedo of papeda (*C. micrantha*, blue circle); in different varieties of pummelo (*C. grandis*, red circles), citron (*C. medica*, yellow circles), and mandarin (*C. reticulata*, orange circles); and in marsh grapefruit (grey circle). Quantitative data, expressed as means ± standard errors of four samples each of Reinking and Tahiti pummelo and five samples each of all other varieties, have been reported previously^10^.Trace amounts of metabolites were set at zero for figure construction. Varieties of pummelo tested included Chandler, Deep Red, Kao Pan, Pink, Reinking, Seedless, and Tahiti pummelo; varieties of cintron tested included Buddha’s hand, Corsican, and Etrog citron; and varieties of mandarin tested included Beauty, Cleopatra, Dancy, Fuzhu, Nan Feng Mi Chu, Owari Satsuma, San Hu Hong Chu, Shekwasha, Sunki, Wase Satsuma, and Willowleaf mandarins. All mandarin varieties tested were domesticated, possessing pummelo-derived genomic segments. The FC molecules assayed included 6’,7’-dihydroxybergamottin, 8-geranyloxypsoralen, bergamottin, bergapten, bergaptol, byakangelicin, byakangelicol, cnidicin, cnidilin, epoxybergamottin, heraclenin, heraclenol, imperatorin, isoimperatorin, isopimpinellin, oxypeucedanin, oxypeucedanin hydrate, phellopterin, psoralen, xanthotoxin, and xanthotoxol, with 6’,7’-dihydroxybergamottin, 8-geranyloxypsoralen, bergamottin, and epoxybergamottin being *O*-geranylated FC derivatives. **b** Gene structures of the *CpPT1* orthologs of pummelo (*C. grandis*) and citron (*C. medica*) deposited in the *Citrus sinensis* annotation project (CsAP) genomic database, along with the FC profiles of their flavedo as determined in **(a)**^10^. The citron-specific insertion at the end of the third exon was confirmed by PCR in three citron varieties, Corsican, Etrog, and Buddha’s hand citron, in which none of the four major *O*-geranylated FC derivatives was detectable^10^. The coding sequence (CDS) of *CpPT1* and the related pummelo genomic sequence are shown to indicate the exon-intron structure proximate to the insertion. **c** Isolation of functional *CpPT1* orthologs from papeda. UV chromatograms at 310, 300, and 330 nm of GT assay mixtures of *C. micrantha* PT1a/b (CmiPT1a/b) with the aromatic substrates bergaptol (Bol, No. 12), xanthotoxol (Xol, No. 13), and 5-hydroxy-7-methoxycoumarin (5H7M, No. 7), respectively, with CpPT1TP-sGFP used as a negative control. All chromatograms are shown at a comparable scale except for that of standard. The numerical codes of the prenyl acceptors are identical to those in Supplementary Fig. 5a and Table 2.

The relationship between *CpPT1* orthologs and the coumarin profile was investigated in members of the genus *Citrus*. A blastn search using the *CpPT1* CDS detected a single close homolog each in the genomes of pummelo, citron, and pure mandarin (Supplementary Fig. 7a). Because genomic information on papeda was unavailable, we isolated two full-length CDSs highly homologous to *CpPT1* from papeda by RT-PCR and named them *CmiPT1a/b* (Supplementary Fig. 8a).

The pummelo genome has a putative *CpPT1* orthologous gene, in accordance with the parent-child kinship between pummelo and grapefruit (Fig. 3b and Supplementary Fig. 7a)^29^. Biochemical characterization demonstrated that, like *CpPT1*, papeda *CmiPT1a/b* encode functional *O*-GTs for both bergaptol and xanthotoxol (Fig. 3c and Supplementary Fig. 8b–d). In contrast, the citron *CpPT1* ortholog has an insertion of an eight-bp repeat containing an in-frame stop codon at the 3’ end of its third exon (Fig. 3b and Supplementary Fig. 7a). This insertion was confirmed by PCR sequencing of the genomes of the three citron varieties previously shown to be devoid of *O*-geranylated FCs (Fig. 3b)^10^. In contrast, these citron varieties contain two other *O*-geranylated coumarins, 5G7M and auraptene^10^, suggesting that these citron varieties possess GPP pools available for coumarin prenylation. Together with previous findings^10^, these results suggest that the eight-bp insertion causes loss of function of the citron *CpPT1* ortholog, resulting in the absence of *O*-geranylated FCs from this species. The *CpPT1* ortholog in pure mandarin was found to contain a deletion and an insertion (Supplementary Fig. 7), consistent with undetectable or very low accumulation of *O*-geranylated FCs in domesticated mandarin varieties^10^.

### Isolation of a coumarin *O*-PT from Apiaceae

*O*-prenylated coumarins have also been detected in vegetables and medicinal plants in the family Apiaceae^2, 19^. As this family is taxonomically distant from Rutaceae in angiosperms^32^, we sought to identify an *O*-PT gene involved in the synthesis of *O*-prenylated coumarins in *Angelica keiskei* to obtain evolutionary insight into the emergence of aromatic *O*-prenylation activity in plants. *A. keiskei* is a medicinal plant endemic to Japan and is locally consumed as a vegetable^19^. We selected a variety of *A. keiskei* accumulating *O*-dimethylallylated bergaptol (isoimperatorin) and its oxidative derivatives, as well as *C*-prenylated chalcones^19^. Biochemical characterization of the *O-* dimethylallyltransferase (DT) activities for FCs using crude enzymes prepared from leaves of *A. keiskei* showed that the native bergaptol 5-*O-*DT (B5ODT) activity leading to the synthesis of isoimperatorin required divalent cations as a cofactor and was associated with the cell membrane (Supplementary Fig. 9a–d). Native *A. keiskei* microsomes also possessed xanthotoxol 8-*O*-DT activity (X8ODT), resulting in the production of imperatorin, although this product was not detected in the *A. keiskei* plants used in this study (Supplementary Fig. 9e and f). These findings suggest that the UbiA protein superfamily is involved in the *O*-prenylation of FCs in Apiaceae as well as in Rutaceae.

Using degenerate primers designed based on conserved amino acid regions among UbiA PTs, a full-length CDS was isolated by RT-PCR and subsequent rapid amplification of cDNA ends (RACE)from the leaves of *A. keiskei* (Supplementary Fig. 3a). This gene, named *AkPT1*, was found to encode a protein with the three conserved polypeptide features of UbiA PTs, similar to CpPT1 (Supplementary Fig. 3 and 10). *In vitro* enzymatic characterization using the *N. benthamiana* transient expression system demonstrated that AkPT1 has B5ODT and X8ODT activities (Fig. 4, Supplementary Fig. 11a, and Supplementary Fig. 12a and b). AkPT1 did not prenylate umbelliferone and isoliquiritigenin, the prenyl acceptor involved in the biosynthesis of *C*-prenylated chalcones in *A. keiskei* (Fig. 4c). For bergaptol and xanthotoxol, this enzyme accepted GPP less efficiently than DMAPP as a prenyl donor (Fig. 4c and Supplementary Fig. 12c–f), suggested that, in *A. keiskei*, AkPT1 acts primarily as a coumarin 5/8-*O*-DT. The enzymatic properties associated with the B5ODT activity of AkPT1 were also determined (Supplementary Fig. 11b–d).

**Fig. 4.**
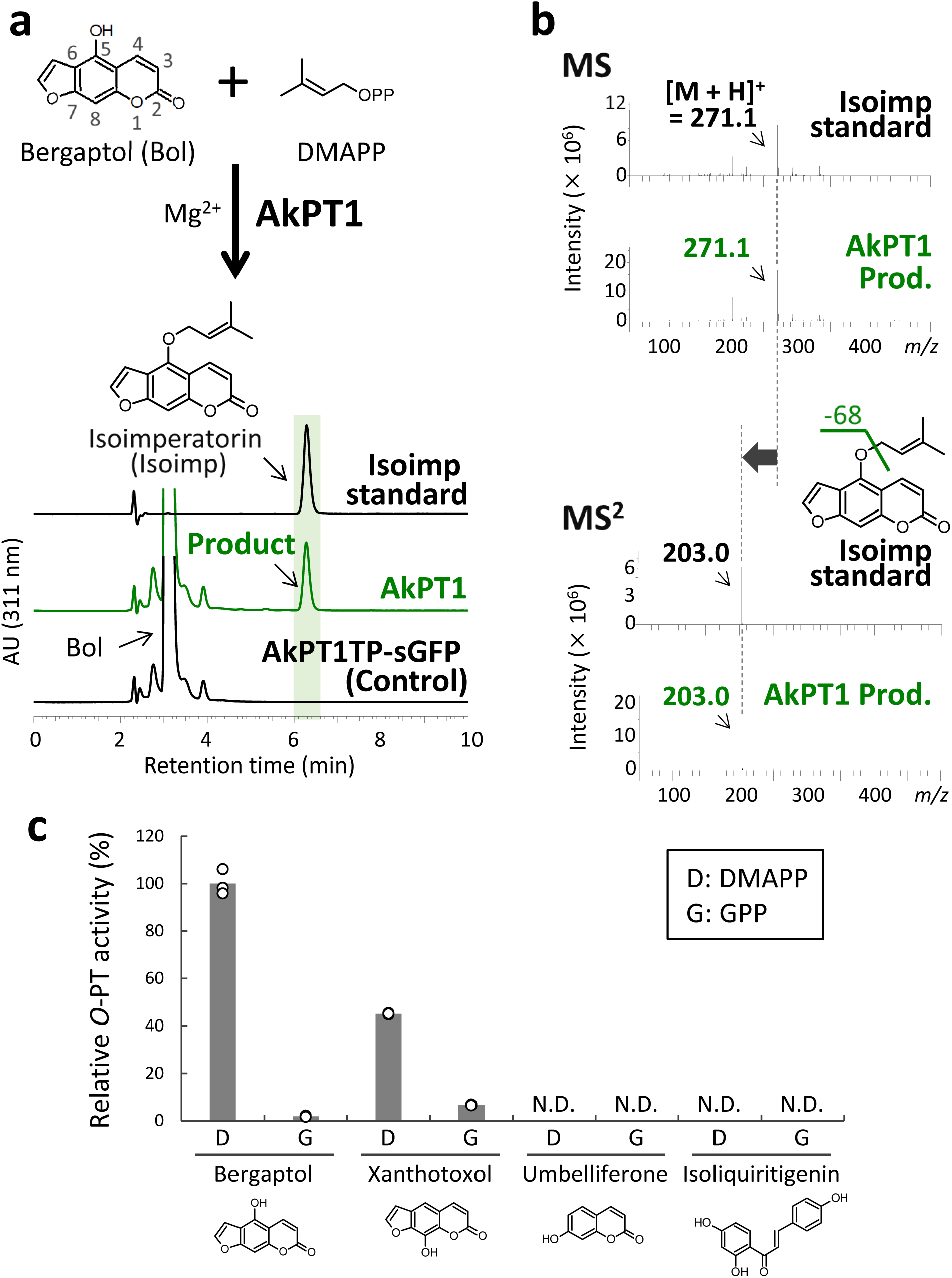
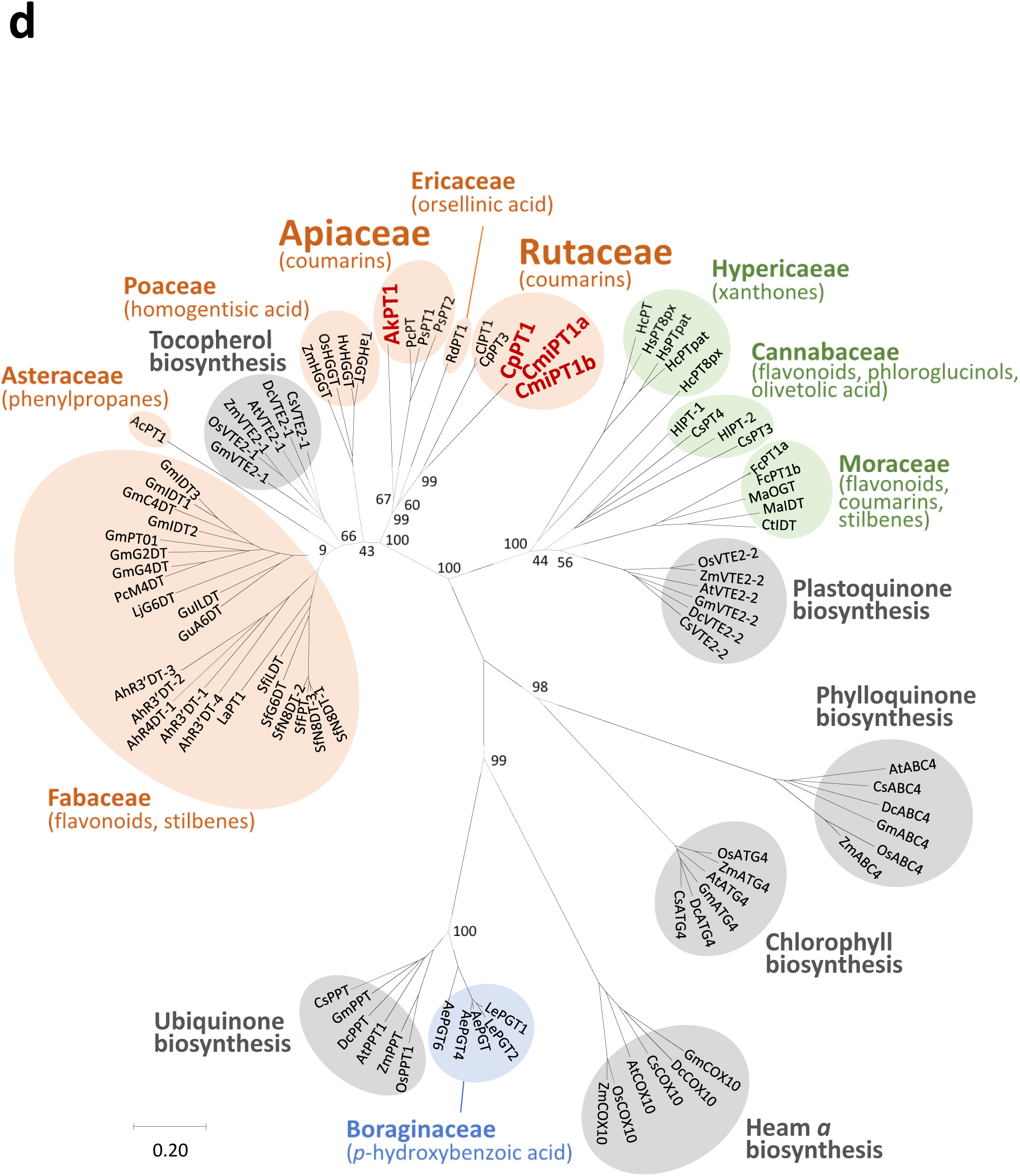
Phylogenetic relationship of aromatic *O*-PTs in the UbiA superfamily from Rutaceae and Apiaceae. **a** UV chromatograms at 311 nm of B5ODT reaction mixtures with *N. benthamiana* leaf microsomes containing recombinant AkPT1 and AkPT1TP-sGFP (negative control). **b** MS^2^ spectrum of the reaction product in the positive ion mode. The loss of 68 mass units probably corresponds to fragmentation caused by the loss of the *O*-dimethylallyl moiety attached to the FC structure. **c** Substrate specificity of AkPT1. Bars represent AkPT1 relative to the average B5ODT activity in triplicate samples. N.D., not detected. **d** Neighbor-joining phylogenetic tree of UbiA proteins. The tree was constructed with 1,000 bootstrap tests based on a ClustalW multiple alignment of UbiA proteins, including CpPT1, CmiPT1a/b, and AkPT1. Bootstrap values are shown for nodes separating clades and for nodes between *C*-PTs and *O*-PTs in Rutaceae and Apiaceae. The bar represents an amino acid substitution rate per site of 0.20. The clades of primary metabolism-related proteins are marked with grey circles. The clades of secondary metabolism-related PTs are highlighted with differently colored circles depending on their possible ancestors, with VTE2-1-, VTE2-2-, and PPT-related clades indicted in orange, green, and blue, respectively, together with their aromatic substrates at the family scale. Apiaceae and Rutaceae *O*-PTs are shown in red. Detailed information about input sequences is provided in Supplementary Table 3.

### Phylogenetic relationship of aromatic *O*-PTs from distant angiosperm families

CpPT1, CmiPT1a/b, and AkPT1 were subjected to phylogenetic analysis. Both neighbor-joining and multiple likelihood-based phylogenetic trees showed that, of the six primary metabolism clades, *O*-PTs from both plant families were located closest to the VTE2-1 clade responsible for tocopherol biosynthesis (Fig. 4d and Supplementary Fig. 13). These findings suggest that molecular evolution of *VTE2-1* is responsible for the emergence of *O*-PT genes in both Rutaceae and Apiaceae. Of the PTs possibly derived from VTE2-1s, however, CpPT1 and CmiPT1a/b were included in one clade together with citrus *C*-PTs, *i.e.*, ClPT1 and CpPT3^21^, whereas AkPT1 was included in another clade, together with the other Apiaceous PTs catalyzing umbelliferone *C*-dimethylallylation, which is involved in the formation of FC core structures, such as PcPT1, PsPT1, and PsPT2 (Fig. 4d and Supplementary Fig. 13)^33, 34^. A search for *CpPT1* orthologs in public transcriptomes of Apiaceous species that produce *O*-dimethylallylated FCs detected no candidate *CpPT1* orthologs (Supplementary Tables 4 and 5)^35^. Similarly, no *AkPT1* orthologs were detected in our grapefruit flavedo transcriptome (Supplementary Table 6). These *in silico* analyses suggest that the two *O*-PT genes of Rutaceae and Apiaceae each evolved independently from *VTE2-1* in a parallel manner. Although two aromatic *O*-PT genes belonging to the UbiA superfamily have been reported in bacteria^36, 37^, CpPT1, CmiPT1, and AkPT1 show substantially higher homologies with other plant than with bacterial UbiA *O*-PTs (Supplementary Table 7), indicating that these plant *O*-PTs emerged in a plant taxon-specific manner.

## Discussion

Aromatic prenylation diversifies the chemical structures of plant metabolites, as these enzymes vary widely in substrate specificity for both prenyl donors and acceptors and regio-specificity^38^. In addition, the structures of these metabolites are further altered by the chemical modifications of transferred prenyl moieties, through, for example, hydroxylation, cyclization and dimerization^38^. These diversities in prenylation and subsequent chemical modification resulted in the diversification of their biological activities^39^. To date, several UbiA PTs catalyzing aromatic *C*-prenylations have been reported to be involved in plant primary and secondary metabolism^20^, whereas the present study showed the divergent evolution of plant UbiA proteins into aromatic *O*-PTs. Similar to aromatic *C*-prenylation by other UbiA PTs, CpPT1 catalyzes both substrate- and regio-specific reactions. UbiA *C*-PTs transfer prenyl moieties to carbons at the *ortho*-positions of phenolic hydroxy moieties, with VTE2-1, the predicted ancestor, showing this regio-specificity^40^. CpPT1 was found to generate only *O*-geranylated products from two simple coumarin derivatives possessing the C5-hydroxy moiety (No. 3 and No.7), despite the availability of the C6 position on these molecules for *C*-prenylation. Thus, CpPT1 able to specifically catalyze O-prenylation may have derived from a neo-functionalized form of C-PT.

The enzymatic properties of native B5OGT were similar to those of the recombinant CpPT1 and the native *O*-PTs detected in citrus flavedo^8^. The expression pattern of *CpPT1* among grapefruit organs matches the accumulation pattern of its reaction products, including bergamottin and its derivatives. The predicted or biochemically verified functions of CpPT1 orthologs were associated with the accumulation of major citrus *O-*geranylated FCs in the flavedo of various ancestral *Citrus* species, strongly suggesting that CpPT1 and its orthologues play pivotal roles in the *O*-geranylation of FCs in *Citrus* species. FCs also accumulate in the pulp or flesh of citrus fruits, although these concentrations are generally lower than in flavedo^10^. However, *O*-geranylated FCs are undetectable in the pulp of citron varieties, which possess *CpPT1* orthologs with an eight base pair insertion resulting in an in-frame stop codon (Supplementary Fig. 14)^10^. Thus, this gene may be a promising target that can weaken grapefruit-drug interactions during the breeding of citrus fruits. Accumulation of other prenylated coumarins, such as auraptene and *O*-dimethylallylated FCs, in citron varieties containing an insertion in the *CpPT1* gene, suggests the presence of coumarin *PT*(*s*) distinct from *CpPT1* or *ClPT1* orthologs in citrus genomes.

Although *CpPT1* is well conserved among members of the genus *Citrus*, the phylogenetic analysis of CpPT1, CmiPT1a/b and AkPT1 suggested that Rutaceae and Apiaceae independently acquired aromatic *O*-prenylation activity in a parallel evolutionary manner, providing another example of repeated molecular evolution of plant UbiA proteins. These proteins were found to evolve independently toward the biosynthesis of specialized PTs by accepting similar or the same aromatic substrates such as flavonoids and coumarins in convergent evolutionary manners^23, 41^. Therefore, repeated molecular evolutionary pathways of members of the UbiA superfamily regarding different aspects of aromatic prenylation ability could underlie the biosynthesis of no less than 1,000 prenylated aromatics in plants^38^.

Because *O*-prenylation enhances the relevant biological activities of FCs, both Rutaceae and Apiaceae likely acquired coumarin *O*-prenylation ability for chemical defenses^5, 42^. Interestingly, the coumarin accumulation patterns of these plant families are similar. Rutaceae and Apiaceae store large quantities of *O*-prenylated coumarins in oil cavities and oil ducts, respectively, with both of them being extracellular compartments filled with hydrophobic metabolites, such as essential oil terpenes^11, 43, 44^. *O*-Prenylation largely increases the hydrophobicity of aromatic molecules due to the masking of hydroxyl residues by hydrophobic prenyl chains. This reaction may enhance the accumulation of coumarins in hydrophobic extracellular compartments, although the mechanisms by which hydrophobic metabolites are exported to such compartments are undetermined. In addition, FCs are generally toxic, being responsible for photo-induced genotoxicity and P450 inactivation^18, 42, 45^, suggesting that sequestering these molecules from vital organelles by storing them in extracellular compartments reduces the risk of self-toxicity. This strategy would be complementary to glycosylation associated self-resistance mechanism that increases the hydrophilicity of specialized metabolites, allowing their sequestration in intracellular vacuoles^46^.

In summary, the present study provides experimental evidence for the functional diversification of plant UbiA proteins to aromatic *O*-PTs, an evolutionary process that likely occurred independently in Rutaceae and Apiaceae. Identification of *CpPT1* may enable the efficient creation of citrus varieties showing reduced grapefruit-drug interactions. Correlation between *CpPT1* gene expression patterns and/or genotypes with coumarin profiles may help determine coumarin metabolism in this agronomically important genus, in which few genes have been identified to date^47^. Knock out of genes responsible for the formation of FC backbones may reduce grapefruit-drug interactions as well as citrus phototoxicity caused by FCs photosensitization^48^, which limits the application of citrus essential oils as cosmetic ingredients^49, 50^. The diverse pharmaceutical activities of *O*-prenylated coumarins, often due to *O*-prenyl moieties^2–4, 13, 28^, suggest that the coumarin *O*-PT genes identified in this study could serve to produce valuable varieties of coumarins.

## Materials and methods

### Plant materials and reagents

Grapefruits (*Citrus* × *paradisi* cv. Marsh) grown at Yuasa farm of Kindai University were collected and different organs (e.g. young leaves, mature leaves, buds, and the albedo and flavedo of immature and mature fruits) prepared as described (Supplementary Fig. 1). Other citrus samples were grown at and collected the Agronomic Research Station INRA/CIRAD of San Giuliano in Corsica (France). *Angelica keiskei* plants (the Oshima variety) for pilot experiments were maintained in the soil field of the Yamashina Botanical Research Institute, Nippon Shinyaku Co., Ltd. (Japan), and the same variety of *A. keiskei* for main experiments were commercially purchased in Japan. Plant tissues were immediately frozen in liquid nitrogen and stored at –80°C if necessary. Phenolic compounds and prenyl diphosphates for characterization of CpPT1 were purchased from Sigma Aldrich (St. Louis, MO, USA), Herboreal Ltd (Dalkeith, UK), Extrasynthase (Lyon, France), Tokyo Chemical Industry Co., Ltd (Tokyo, Japan), and Indofine Chemical Company (Hillsborough, NJ, USA). DMAPP and GPP for the other experiments were synthesized as described^51^, and kindly provided by Dr. T. Kuzuyama (The University of Tokyo) and Dr. T. Kawasaki (Kyoto University), respectively. Auraptene standards were kindly provided by Dr. A. Murakami (University of Hyogo) and Dr. Y. Uto (Tokushima University), and 8-geranylumbelliferone was generously provided by Dr. Y. Uto.

### Construction of transcriptome datasets from immature and mature flavedo samples

Immature and mature grapefruit flavedo tissue samples were each ground to fine powder with mortars and pestles. Total RNA was were extracted from each using RNeasy Plant Mini kits (Qiagen, Hilden, Germany) and cDNA libraries were prepared using NEBNext® Ultra™ RNA Library Prep Kits for Illumina® (New England BioLabs, Ipswich, MA, USA) according to the manufacturers’ protocols. The two resulting cDNA libraries each consisted of approximately 400–600 bp of grapefruit cDNA sequences which were tagged with different index sequences for analysis of comparative expression. The cDNA libraries were purified using MinElute Gel Extraction Kits (Qiagen) and quantified with KAPA Library Quantification kits (Roche, Basel, Switzerland). The libraries were diluted to 4 nM, mixed and sequenced by MiSeq (illumina, San Diego, CA, USA). The resulting pair-end reads (2 × 301 bp) were filtered with Trimmomatic to remove low-quality bases and adaptor sequences. The remained reads were *de novo* assembled to 64,959 contigs using Trinity. TPM and fragments per kilobase of transcript per million mapped reads (FPKM) of contigs in the two flavedo samples were calculated with RNA-seq by expected maximization (RSEM) based on a reference sequence set created with Bowtie2. Contigs were annotated with Blast2GO (https://www.blast2go.com/).

### Quantification of coumarins in grapefruit organs

Coumarins were extracted from grapefruit leaves, buds, immature fruit flavedo, immature fruit albedo, mature fruit flavedo, and mature fruit albedo by suspending 10 mg dry weight of each powdered sample in 400 μl of 80% (v/v) methanol, vortexing, and centrifuging at 10,000 × *g* for 5 min. This extraction procedure was performed three times and the three resulting supernatant fractions were pooled. Each suspension was filtered through a 0.45-μm Minisart RC4 filter (Sartorius, Göttingen, Germany) and analyzed by liquid chromatography/mass spectrometry (LC/MS) using an Acquity ultra-high-performance liquid chromatography (UPLC) H-Class/Xevo TQD system (Waters, Milford, MA, USA). Separation conditions were: sample injection, 2 μl; an Acquity UPLC BEH C18 column (1.7 µm, 2.1 × 50 mm; Waters) with a UPLC BEH C18 VanGuard pre-column (1.7 µm, 2.1 × 5 mm) at 40 °C; mobile phases, solvent A (water containing 0.1% (v/v) formic acid) and solvent B (acetonitrile) using elution programs of 10%–90% B from 0–15 min (linear gradient), 90% B from 15–16 min, 100% B from 16–20 min, and 10% B from 20–25 min; and flow rate, 0.2 ml min^-1^. MS conditions were: positive electrospray ionization mode; source temperature, 150 °C; desolvation gas temperature, 400 °C, nebulizer N_2_ gas flow rate, 50 l h^-1^, desolvation N_2_ gas flow rate, 800 l h^-1^, capillary voltage, 3.15 kV; and cone voltage, 35 V. Coumarins were detected by selected ion recording mode with *m/z* = 299.2 for auraptene, *m/z* = 339.3 for bergamottin, *m/z* = 373.3 for 6’,7’-dihydroxybergamottin, *m/z* = 315.2 for epoxyauraptene, and *m/z* = 355.3 for epoxybergamottin. Data were analyzed using MassLynx v. 4.1 software (Waters). The coumarin contents were calculated from peak areas using calibration curves constructed using the authentic compounds.

### Isolation of citrus PT genes and construction of their plant expression plasmids

Immature grapefruit flavedo was ground to fine powder with a mortar and a pestle. Total RNA was extracted using RNeasy Plant Mini kits (Qiagen), followed by reverse transcription with SuperScript™ III Reverse Transcriptase (Thermo Fisher Scientific, Waltham, MA, USA). The full CDS of *CpPT1* in the cDNA was PCR amplified with KOD-plus neo (Toyobo, Osaka, Japan) and the primer pair CpPT1_Fw and CpPT1_Rv (Supplementary Table 8). The PCR product was subjected to adenine overhanging and inserted into pGEM T-easy vector (Promega, Madison, WI, USA) for sequencing. *CpPT1* CDS was PCR amplified using KOD-plus neo and the primer pair CpPT1_TOPO_Fw1 and CpPT1_TOPO_Rv (Supplementary Table 8). The amplicon was inserted into the vector pENTR™/D-TOPO™ (Thermo Fisher Scientific) by directional TOPO reaction and finally into the vector pGWB502^52^ by LR recombination to yield a construct containing *P35S-CpPT1-TNos*.

The full CDSs of *CpPT2* and *CpPT3* were amplified from grapefruit flavedo by RT-PCR using TaKaRa Ex Taq polymerase (Takara, Kusatsu, Japan) and the primer pairs for *CpPT2* (CpPT2_Fw and CpPT2_Rv) and *CpPT3* (CpPT3_Fw and CpPT3_Rv), and inserted into the vector pMD-19 by TA cloning for sequencing. The full CDS of *CpPT2* in the pMD-19 vector was introduced into the vector pRI201 by double digestion with *Nde*I and *Sal*I followed by ligation. The full CDS of *CpPT3* was PCR amplified using pMD19-*CpPT3* as a template, KOD-plus (Toyobo), and the primer pair CpPT3_BamHI_Fw and CpPT3_XhoI_Rv (Supplementary Table 8). The amplicon was subcloned into pGEM T-easy, digested with *Bam*HI and *Xho*I, and ligated into the vector pENTR2B (Thermo Fisher Scientific) that had been similarly digested. The *CpPT3* CDS was subsequently subcloned into the vector pGWB502 by LR recombination^52^. pRI201-*CpPT3* was constructed in a manner similar to that for pRI201-*CpPT2*.

*C. micrantha* leaves were ground to fine powder with a mortar and a pestle, and total RNA was extracted using the protocol for difficult samples in E.Z.N.A.^®^ Plant RNA Kits (Omega Biotek, Norcross, GA, USA). The samples were reverse transcribed and specific sequences were amplified using the SuperScript™ One-Step RT-PCR System with Platinum™ *Taq* DNA (Thermo Fisher Scientific) and the primer pair CmiPT1_fw and CmiPT1_Rv (Supplementary Table 8). The amplicon was subcloned into the vector pCR^™^8/GW/TOPO^®^ (Thermo Fisher Scientific) by TA cloning for sequencing, and inserted into the vector pENTR2B by in-fusion reaction using *Bam*HI and *Xho*I sites, and introduced into the vector pGWB502 by LR recombination^52^.

### Isolation of *AkPT1* and construction of plant expression vectors for *in vitro* characterization

Leaves of *A. keiskei* plants (the Ohshima variety) maintained in the Yamashina Botanical Research Institute were crushed to fine powder. Total RNA was extracted with RNeasy Plant Mini kits, genomic DNA was removed with DNA-*free*^TM^ (Thermo Fisher Scientific), and first-strand cDNA was synthesized with SuperScript III Reverse Transcriptase. The cDNA pool was used as a template for PCR amplification with the degenerate primer pair AkPT_DGP1_Fw and AkPT_DGP1_Rv (Supplementary Table 8). The resulting PCR products were utilized as templates for the second PCR amplification using a second pair of degenerate primers AkPT_DGP1_Fw and AkPT1_DGP2_Rv (Supplementary Table 8). Detailed conditions for these amplifications have been described^53^. The products of the second PCR reaction were inserted into pGEM T-easy for sequencing. Based on isolated partial sequences, the full CDS of a PT gene named *AkPT1a* was obtained by 5’- and 3’-RACE using the internal gene-specific primers AkPT1_5’RACE_Rv and AkPT1_3’RACE_Fw (Supplementary Table 8), respectively, and the SMARTer RACE cDNA Amplification Kit (Takara) according to the manufacturers’ guidelines.

Because the CDS of *AkPT1a* possibly had one PCR-error-derived mutation, this gene was again isolated. A cDNA pool was also prepared from leaves of *A. keiskei* plants (the Ohshima variety), which were commercially obtained, essentially as described above. PCR using this cDNA sample as a template, KOD-plus neo (Toyobo), and the primer pair for *AkPT1a* (AkPT1_ Fw and AkPT1_Rv) (Supplementary Table 8) amplified the full CDS homologous to *AkPT1a*, named *AkPT1b*, which was subsequently renamed *AkPT1* and used for all experiments in this study. The CDS of *AkPT1b* was PCR amplified using KOD-plus neo and the primer pairs AkPT1_TOPO_Fw and AkPT1_TOPO_Rv (Supplementary Table 8), and inserted into the vector pGWB502^52^ via the pENTR™/D-TOPO™ vector.

### Enzymatic characterization of CpPT1 and AkPT1

Plasmids expressing PTs were individually introduced into *Agrobacterium tumefaciens* LBA4404 strain and these transformants were co-infiltrated into *N. benthamiana* leaves, along with the *A. tumefacien* C58C1 strain harboring the plasmid pBIN61-P19^54^. Microsomes were prepared from these leaves as described^33^, suspended into 100 mM Tris-HCl, pH 8.0, buffer, and stored at −80°C. Microsome preparations were diluted into reaction buffer, 50 mM Tris, 20 mM MES-HCl (pH 7.6), depending on the potencies of PT activities. Standard B5OGT reaction mixtures (100 µl) containing microsomes, 400 µM bergaptol, 400 µM GPP, and 10 mM MgCl_2_, pH 7.6, were incubated at 28 °C for 20 h unless otherwise indicated. In substrate screening, the concentrations of prenyl acceptor and donor substrates were both set at 200 µM. In kinetic analysis, the concentrations of substrates ranged from 1–200 µM for bergaptol, 5–500 µM for xanthotoxol, 10–750 µM for 5,7-dihydroxycoumarin, 20–750 µM for 5-hydroxy-7-methoxycoumarin, 20–1000 for µM 8-hydroxybegapten, and 2–500 µM for GPP.

For enzymatic characterization of AkPT1, a standard mixture (200 µl) containing 250 µM prenyl acceptor, 250 µM prenyl donor, 10 mM MgCl_2_, and AkPT1 microsomes suspended in 100 mM Tris-HCl containing 1 mM DTT (pH 8.0) was incubated for 60 min at 30 °C. In kinetic analysis, the concentrations of substrates ranged from 2–63 µM for bergaptol and 2–16 µM for DMAPP.

### Extraction of reaction products

CpPT1 enzymatic reactions were stopped by the addition of 10 µl of 1 M HCl. The substrates and product were extracted with 500 µl of ethyl acetate. The mixtures were vortexed for 15 min and centrifuged at 5,200 × *g* for 5 min, and 450 µl of each upper phase was collected. In substrate screening, this procedure was repeated once using an additional 500 µl of ethyl acetate, and the resulting ethyl acetate fraction (450 µl) was combined with the first fraction. The combined extract was vacuum evaporated to dryness, and dissolved in 100 µl of methanol by vortexing for 15 min. After centrifugation for 30 min at 24,100 × *g*, the supernatant was subjected to LC/MS analysis. Chemicals in AkPT1 reaction mixtures were obtained by one-cycle ethyl acetate extraction using essentially the same protocol.

### LC/MS analysis of extracts of CpPT1 reaction mixtures

Enzyme products were detected and quantified with a NEXERA UHPLC system (Shimadzu, Kyoto, Japan) equipped with a photodiode array (PDA, SPDM20A, Shimadzu). The chromatographic column was a C18 reverse phase column (LC Kinetex XB-C18 100 Å, 1.8 μm, 150 × 2.1 mm, Phenomenex). The products were separated using a gradient of solvent A (water with 0.1 (v/v) formic acid) and solvent B (acetonitrile with 0.1 % (v/v) formic acid), consisting of 20% solvent B at 0.01 min; 20% B at 0.74 min; 90% B at 8.00 min; 100% B at 10.00 min; 100% B at 15.00 min; 20% B at 15.01 min; and STOP at 17.51 min. The flow rate was set at 0.3 ml min^-1^. Enzymatic products were screened in a range of 250–370 nm. MS was performed on a Shimadzu MS2020 (Shimadzu) in both positive and negative modes.

LC/MS^2^ analyses were performed on a Dionex Ultimate 3000 UHPLC Chain equipped with a Thermo LTQ-ORBITRAP detector (Thermo Fischer Scientific) and Phenomenex Kinetex XB-C18 (150 × 2.1mm, 2.6 µm, Phenomenex, Le Pecq, France). Compounds were separated with a gradient program of solvent A (water with 0.1% (v/v) formic acid) and solvent B (acetonitrile with 0.1% (v/v) formic acid), consisting of 10% B from 0 to 1 min, a gradient of 10% to 70% B until 15 min, 100% B at 21 min and maintained for 4 min, and a return to initial conditions over 1 min, at a flow rate of 0.2 ml min^-1^at 40 °C. HESI Probe was used at 300 °C. MS signals were scanned between *m/z* = 100 and 600 in positive mode and MS^2^ data were obtained for the ten most intense MS signals.

### LC/MS analyses of extracts of AkPT1 reaction mixtures

Extracts of AkPT1 reaction mixtures were chromatographically separated on a LiChrosphereRP-18 column (4.0 mm × 250 mm, Merck) at a flow rate of 1.0 ml min^-1^ and at 40 °C under the control of a D-2000 Elite HPLC system (Hitachi, Tokyo, Japan). An isocratic separation program of 20% (v/v) solvent A (water with 0.3% (v/v) acetic acid) and 80% (v/v) solvent B (methanol with 0.3% (v/v) acetic acid) was used except for screening of xanthotoxol *O*-DT activity. Extracts of xanthotoxol *O*-DT assay were analyzed with a linear gradient program, consisting of 20% to 90% (v/v) of solvent B (methanol with 0.3% (v/v) acetic acid) in solvent A (water with 0.3% (v/v) acetic acid) over 45 min. Enzymatic products were scanned at a range of 200–370 nm with a L2445 Diode Array Detector (Hitachi).

Reaction products of AkPT1 were identified using LC-IT-TOF-MS (Shimadzu), a TSK gel ODS-80Ts column (2 mm × 250 mm, Tosoh) and a linear gradient program composed of 20% to 80% (v/v) solvent B (acetonitrile with 0.1% (v/v) formic acid) in solvent A (water with 0.1% (v/v) formic acid) at a flow rate of 0.2 ml min^-1^ and at 40 °C. Precursor ions for MS^2^ analysis were selected in a range of *m/z* = 50–500.

### Quantitative RT-PCR

Total RNA was prepared with RNeasy Plant Mini Kits (Qiagen), and contaminating DNA was eliminated by treatment with gDNA remover (Toyobo), according to the manufacturers’ instructionss. The RNA was reverse-transcribed to cDNA using ReverTra Ace qPCR RT Master Mix (Toyobo) according to the manufacturer’s instructions. Quantitative PCR was performed on a CFX96 Touch Deep Well system (Bio Rad, Hercules, CA, USA), using Thunderbird SYBR qPCR Mix (Toyobo) according to the manufacturers’ instructions. Each PCR mixture consisted of cDNA template, 7.5 pmol of each *CpPT1* (CpPT1_qPCR_Fw and CpPT1_qPCR_Rv) or *CpEF1α* (CpEF1α_Fw and CpEF1α_Rv) primer (Supplementary Table 8), 0.5 μl of fluorescent probe and 12.5 μl of Thunderbird SYBR qPCR Mix in a total volume of 25 μl. The amplification protocol for *CpEF1α* consisted of an initial denaturation at 95°C for 2 min, followed by 40 cycles of denaturation at 95°C for 15 s, annealing at 55°C for 30 s, and extension at 72°C for 30 s, whereas the amplification protocol for *CpPT1* consisted of an initial denaturation at 95°C for 2 min, followed by 40 cycles of denaturation at 95°C for 15 s and annealing and extension at 60°C for 30 s.

### Construction of plasmids for expression of GFP-fusion proteins and microscopic observation

For subcellular localization analysis, *CpPT1TP* encoding the first 70 amino acids of CpPT1; *CpPT1* (-stop) encoding the full CDS without the stop codon of CpPT1; *CpPT3TP* encoding the first 55 amino acids of CpPT3; and *AkPT1TP* encoding the first 60 amino acids of AkPT1 were PCR amplified using their respective primer pairs, CpPT1_TOPO_Fw2 and CpPT1_TP210_Rv, CpPT1_TOPO_Fw2 and CpPT1_woStop_Rv, CpPT3_TOPO_Fw and CpPT3_TP165_Rv, and AkPT1_TOPO_Fw and AkPT1_TP180_Rv (Supplementary Table 8), and KOD-plus enzyme kits (Toyobo). The resulting PCR products were introduced into the pGWB505 vector by directional TOPO cloning using the pENTR™/D-TOPO™ vector and subsequent LR recombination, yielding constructs containing *P35S-CpPT1(-stop)-sGFP-Tnos*, *P35S-CpPT1TP-sGFP-Tnos, P35S-CpPT3TP-sGFP-Tnos,* and *P35S-AkPT1TP-sGFP-Tnos*. CpPT1TP-sGFP, CpPT3TP-sGFP, and AkPT1TP-sGFP were also used as negative controls for *in vitro* characterization of CpPT1, CpPT3, and AkPT1, respectively. The GFP-fusion proteins of CpPT1 and AkPT1 were transiently expressed in *N. benthamiana* leaves by agroinfiltration and in onion epidermal cells by particle bombardment, respectively, and microscopic analysis were performed when using of pHKN29 containing *P35S*-*sGFP*-*Tnos* and pWxTP-DsRed as controls for free sGFP and plastid localizations^55, 56^, respectively, as described^22^.

### Isolation of partial *CpPT1* genomic sequence in citron varieties

Flavedo slices of Corsican citron, Etrog citron, and Buddha’s Hand citron were ground with mortars and pestles to fine powder and their genomic DNA was extracted using E.Z.N.A. ^®^ SP Plant DNA Kit (Omega Biotek). Partial genomic sequences of *CpPT1* were PCR amplified in these preparations using SapphireAmp Fast PCR Master Mix (Takara) and the primer pair citron_Fw and citron_Rv (Supplementary Table 8). The PCR products were cloned into pCR^™^8/GW/TOPO for sequencing.

### Biochemical characterization of native microsomes from *A. keiskei* leaves

The microsomal fractions from ca. 10 g of *A. keiskei* leaves were prepared essentially as described^8^. The *in vitro* characteristics of coumarin PT activities in the microsomal fractions from *A. keiskei* leaves were assessed similar to assessments of microsomes prepared from *N. benthamiana* leaves expressing *AkPT1*.

### *In silico* analysis

The nucleotide sequences in the NCBI (https://www.ncbi.nlm.nih.gov/), Phytozome (https://phytozome.jgi.doe.gov/pz/portal.html#), *Citrus sinensis* Annotation Project (http://citrus.hzau.edu.cn/orange/download/index.php), and OneKP (https://www.onekp.com/) databases were searched for PT sequences^57, 58^. The transit peptides and multiple transmembrane regions of PTs were predicted by ChloroP (http://www.cbs.dtu.dk/services/ChloroP/) and TMHMM Server v. 2.0 (http://www.cbs.dtu.dk/services/TMHMM/), respectively. Local blast searches of the grapefruit flavedo transcriptome and the calculation of amino acid identities among PTs were performed with Bioedit (http://www.mbio.ncsu.edu/BioEdit/bioedit.html). PT sequences were multiply aligned by ClustalW, and neighbor-joining and multiple likelihood phylogenetic trees were constructed using MEGA-X (http://www.megasoftware.net/). During *in silico* searches by blast, the terminals of hit regions of a fished gene or protein were manually adjusted to avoid mapping of a base of query to multiple sites, and the homology between a fished sequence and a query was recalculated relative to all the adjusted hit regions.

### Statistics and reproducibility

*V*_max_ and apparent *K*_m_ were calculated by a non-linear least-squares method using Sigmaplot 12.3. Organ specificities of the expression of *CpPT1* and of the accumulation of metabolites accumulation were statistically analyzed by the Games-Howell test using R software version 3.4.1^59^. The organ specificities of coumarin contents and *CpPT1* expression were analyzed in five buds, five leaves, flavedo and albedo from five immature fruits, and flavedo and albedo from five mature fruits. *In vitro* enzymatic assays were performed in three independent experiments. Chromatograms of *in vitro* enzymatic assays are representative of three independent experiments.

### Data availability

The accession numbers of *CpPT1 – 3*, *CmiPT1a/b*, and *AkPT1* are LC557129 *–* LC557131, LC557132/LC557133, and LC557134, respectively. The raw RNA-seq reads are available as DRA010472.

## Acknowledgments

We thank Dr. Yann Froelicher (CIRAD, UMR AGAPSan Giuliano, France), Dr. Patrick Ollitrault (Centro de Protección Vegetal y Biotecnología, Valencia, Spain), and Dr. Nobumasa Nito (Kindai University) for providing citrus samples. We also thank Dr. David Baulcombe (Cambridge University, UK) for the pBIN61-P19 plasmid, Dr. Tsuyoshi Nakagawa (Shimane University, Japan) for pGWB vectors, Dr. Toshiaki Mitsui (Niigata University) for the pWxTP-DsRed plasmid, and Dr. Hiroshi Kouchi (International Christian University) for the pHKN29 plasmid. We are grateful to Dr. Akira Murakami (University of Hyogo) and Dr. Yoshihiro Uto (Tokushima University) for prenylated coumarin standards, and Dr. Tomohisa Kuzuyama (The University of Tokyo) and Dr. Takashi Kawasaki (Kyoto University) for GPP. We also thank Ms. Keiko Kanai, Mr. Patrick Riveron and Mr. Clément Charles for technical assistance. LC-IT-TOF/MS analyses of the enzymatic characteristics of AkPT1 were performed in collaboration with the Development and Assessment of Sustainable Humanosphere (DASH) system of the Research Institute for Sustainable Humanosphere (RISH), Kyoto University (Japan). Plants were grown on the PEPor platform (Université de Lorraine, France). Transcriptome analysis of grapefruit flavedo tissues was conducted with the technical support of Dr. Tomoaki Sakamoto (Kyoto Sangyo University) in the Plant Global Education Project of Nara Institute of Science and Technology. This project was supported by a Grant–in–Aid for Scientific Research for Plant Graduate Student from the Nara Institute of Science and Technology supported by MEXT. This work was also financially supported by the SAKURA program of JSPS Research Fellowship for Young Scientists (to R.M.), by JSPS Overseas Research Fellowships (to R.M.), by Grants-in-Aid for Scientific Research (No. 26712013 to A.S. and No. 16H03282 to K.Y.), by the New Energy and Industrial Technology Development Organization (NEDO) Project (No. 16100890 to K.Y.), by the Région Grand Est and the French Science Ministry (to C.V.), by the “Bioprolor2” project (Région Grand-Est) (to A.H.), and by the “Impact Biomolecules” project of the “Lorraine Université d’Excellence” (Investissements d’avenir–ANR) (to A.H.). Additional support was provided by RISH, Kyoto University (Mission 5) (to K.Y.).

## Author contributions

R.M., A.S., A.H., F.B., M.M., and K.Y. conceived the research. T.M. maintained the grapefruit trees. R.M. and T. Kurata performed transcriptomic analyses of grapefruit flavedo. R.M. performed *in silico* screening of grapefruit flavedo transcriptome and *in silico* analyses of PT polypeptides. R.M., T.T., and M.M. isolated *CpPTs*. R.M., T.T., J.K., C.K., and M.M. constructed plasmids for characterization of *CpPTs*. R.M., T.T., and A.O. characterized the CpPTs biochemically. R.M., K.T. and T.I. performed microscopic analysis of GFP fusion proteins. E.M. and A.S. performed qRT-PCR of *CpPT1*. M.N. and A.S. quantified coumarin derivatives. F.J., T. Koeduka, and R.M. isolated *AkPT1.* R.M. characterized the *AkPT1* and *A. keiskei* microsomes biochemically. R.M. and C.V. performed phylogenetic analyses. J.G. maintained LC-MS apparatuses and optimized their conditions for this research. H.Y. prepared DMAPP and standard specimens for identification of enzymatic reaction products. R.M., A.H., and K.Y. wrote the manuscript with the contribution of the other authors.

## Supplementary information legends

**Supplementary Fig. 1 Grapefruit samples used in this study**

Grapefruit samples harvested at the Yuasa farm of Kindai University. The flavedo (outer green or yellow peel) and albedo (inner white peel) were separately collected from each fruit.

**Supplementary Fig. 2 Coumarin contents of different grapefruit organs**

Coumarin contents in five biological replicates of grapefruit leaves (L), buds (B), immature fruit flavedo (IF), immature fruit albedo (IA), mature fruit flavedo (MF), and mature fruit albedo (MA). **a–f** Box plots showing quantitation of the major *O*-prenylated coumarins in grapefruit, including **a** auraptene, **b** epoxyauraptene, **c** bergamottin, **d** epoxybergamottin, **e** 6’,7’-dihydroxybergamottin, and **f** total *O*-prenylated coumarins (center line, median; box limits, first and third quartiles; whiskers, minimum and maximum). The chemical structure of these major coumarins are shown in Fig. 1a. Significant differences between groups are indicated by letters (*p* < 0.05 by Games-Howell tests).

**Supplementary Fig. 3 *In silico* analysis of CpPT1–3 and AkPT1 polypeptides**

**a** ClustalW multiple alignment of CpPT1–3, AkPT1 and related PTs. The first and second aspartate-rich motifs are highlighted in red and orange, respectively. CpPT1–3 share 42–49% amino acid identities each other, and 50%, 44%, and 95% identifies, respectively, with ClPT1, CpPT1 and AkPT1 share 36% amino acid identity. Arrows indicate the positions used for the design of degenerate primers, AkPT_DGP1_Fw, Fw; AkPT_DGP1_Rv1, Rv1; AkPT_DGP1_Rv2, Rv2. **b** Transit peptides (TPs) of CpPT1–3 and AkPT1 predicted by ChloroP. **c** Transmembrane (TM) regions of CpPT1-3 and AkPT1 predicted by TMHMM. The possibility of each amino acid residue to be a part of a TM domain is plotted (maximum, 1). The positions of the two aspartate-rich motifs and predicted TPs are also indicated.

**Supplementary Fig. 4 Biochemical screening of CpPT2 and CpPT3**

**a** Chemical structures of tested prenyl acceptor substrates. **b** Substrate specificities of CpPT2 and CpPT3 using DMAPP and GPP as prenyl donor substrates (n =1 or 2). For CpPT2, a mixture of 5-hydroxy-7-methoxycoumarin and xanthotoxol was tested simultaneously. Trace amounts of unidentified products were found in the GT assays with 5-hydroxy-7-methoxycoumarin, bergaptol, and *p*-coumaric acid. On reverse-phase HPLC, the products of 5-hydroxy-7-methoxycoumarin and bergaptol eluted earlier than their *O*-geranylated forms. 8GU, 8-geranylumbelliferone; 6GU, 6-geranylumbelliferone; N.D., not detected. **c and d** Umbelliferone *C*-GT reactions catalyzed by CpPT3. UV chromatograms at 327 nm of umbelliferone GT reaction mixture of CpPT3 and that of ClPT1, a previously described umbelliferone *C*-GT from lemon (**d**). CpPT3TP-sGFP was used as a negative control. UV chromatograms are shown at a comparable scale except for that of standards.

**Supplementary Fig. 5 Enzymatic reactions catalyzed by CpPT1**

**a** Chemical structures of aromatic compounds tested in substrate specificity analysis of CpPT1 (Tables 1). The numerical indicators of the compounds accepted by recombinant CpPT1 are shown in colors dependent on their metabolite groups. **b–i** *O*-GT activities of CpPT1 for coumarins other than bergaptol. **b, d, f, and h** UV chromatograms of GT reaction mixtures of CpPT1 with 5-hydroxy-7-methoxycoumarin (7) (**b**), xanthotoxol (13) (**d**), 5,7-dihydroxycoumarin (3) (**f**), and 8-hydroxybergapten (16) (**h**). CpPT1-sGFP was the negative control, and its UV chromatogram is shown at a scale comparable to that of CpPT1 for each substrate set. 5G7M, 5-geranyloxy-7-methoxycoumarin. **c, e, g, and i** MS^2^ spectra of the reaction products in (**b)**, **(d)**, **(f)**, and **(h)**, respectively, in the positive ion mode. The chemical structures of the reaction products of 5,7-dihydroxycoumarin (3) and 8-hydroxybergapten (16) were predicted to be the 5- and 8-*O*-geranylated forms, respectively, considering the fragmentation patterns of the products in MS^2^ analysis and the acceptance of hydroxy groups at the C5 or C8 position by CpPT1 for other coumarins shown in Tables 1 and 2.

**Supplementary Fig. 6 Properties of the B5OGT activity of CpPT1**

**a** pH preference. Microsomes prepared from *N. benthamiana* leaves expressing *CpPT1* were subjected to B5OGT assays in 50 mM Tris, 20 mM MES-HCl, pH 6.2, 6.8, 7.1, or 7.6, or in 50 mM Tris, 20 mM MES-NaOH, pH 8.1, 8.5 or 8.9. Results are reported as the means ± standard errors of three independent experiments and shown relative to the mean B5ODT activity at pH 7.6.

**b** Divalent cation preference. B5OGT assays of CpPT1-expressing microsomes in buffer containing MgCl_2_, NiCl_2_, CoCl_2_, MnCl_2_, or CaCl_2_. Results are reported as the mean of three independent experiments and shown relative to the mean B5ODT activity in buffer containing MgCl_2_. The negative control consisted of buffer containing EDTA in place of divalent cation. N.D., not detected.

**Supplementary Fig. 7 Structures of *CpPT1* gene orthologs in *Citrus***

**a** ClustalW multiple alignments of *CpPT1* orthologs from pummelo, citron, and pure mandarin and of the CDS of *CpPT1* showing the positions of the exons. Pink and yellow boxes represent deletions and insertions, respectively, predicted to lead to a loss of gene function. Compared with *CpPT1*, the predicted CDSs of pummelo, citron, and pure mandarin orthologs were 100%, 99%, and 99% identical, respectively, except for the regions of insertion and deletion. All genomic sequences were obtained from the public *Citrus sinensis* annotation project genomic database. **b** Gene structure of pure mandarin *CpPT1* based on **(a)**.

**Supplementary Fig. 8 Isolation of *CpPT1* orthologs from papeda**

**a** ClustalW multiple alignments of *C. micrantha* PT1a/b (CmiPT1a/b) and CpPT1 polypeptides. Both proteins showed ca. 98% amino acid identity with CpPT1. **b–d** MS spectra of enzymatic reaction products of CmiPT1a/b in GT assays using bergaptol (**b**), xanthotoxol (**c**), and 5-hydroxy-7-methoxycoumarin (5H7M, **d**) as prenyl acceptor substrates. The enzymatic reaction products were identified by comparisons with standards for the *O*-geranylated forms of the coumarin substrates.

**Supplementary Fig. 9 Furanocoumarin *O*-dimethylallyltransferase activities of *Angelica keiskei* microsomes**

**a** UV chromatograms of the bergaptol 5-*O*-dimethylallyltransferase (B5ODT) assay using native microsomes prepared from leaves of *Angelica keiskei* as crude enzymes. EDTA was used instead of MgCl_2_ as a negative control, with the chromatograms shown at a comparable scale. Bol, begaptol. **b** MS^2^ spectra of the B5ODT reaction product from the fragmentation of its molecular ion peak (*m/z* = 271.1). A loss of 68 mass units was predicted to correspond to fragmentation resulting from the loss of the entire *O*-dimethylallyl moiety attached to the FC structure. **c** Divalent cation requirement of the B5ODT activity in *A. keiskei* leaf microsomes. EDTA was used instead of MgCl_2_ as a negative control. The results are reported as the mean of three independent experiments. N.D., not detected. **d** Membrane localization of the B5ODT activity of *A. keiskei* leaves. A cell-free extract (CFE) was centrifuged at 100,000 × *g* for 30 min to yield a pellet (Ppt.) and supernatant (Sup.), with all three used as crude enzymes. Results shown are the mean of three independent experiments. N.D., not detected. **e** UV chromatograms of the xanthotoxol 8-*O*-DT (X8ODT) assay with *A. keiskei* leaf microsomes. The negative control consisted of incubation without DMAPP, with the UV chromatogramss shown at a comparable scale. Xol, xanthotoxol. **f** MS^2^ spectra of the X8ODT reaction product from the fragmentation of its molecular ion peak (*m/z* = 271.1). The explanation for the predicted loss of 68 mass units is described in **b**, above.

**Supplementary Fig. 10 Subcellular localization of AkPT1**

Microscopic observation of onion epidermal cells expressing **a** free sGFP and **b and c** AkPT1TP-sGFP, a chimeric protein consisting of amino acids 1–60 of AkPT1 and sGFP. These proteins, together with the plastid marker WxTP-DsRed, were introduced into onion epidermal cells by particle bombardment. Arrowheads in **b** indicate regions enlarged in **c**. **d** Negative control, consisting of cells expressing WxTP-DsRed alone. For merging, the brightness and contrast of fluorescent images were adjusted in an unbiased manner, with magenta used as a pseudo-color for the DsRed signal. Scale bars indicate 100 µm in **a, b,** and **d** and 5 µm in **c**.

**Supplementary Fig. 11 Characterization of the B5ODT activity of AkPT1**

**a** Negative control B5ODT assays performed in the absence of bergaptol (-Bergaptol) or DMAPP (-DMAPP), with EDTA instead of MgCl_2_ (EDTA), in the absence of microsomes (-Enzyme), or in the presence of heat-denatured microsomes (Heat-denatured) or microsomes containing AkPT1TP-sGFP instead of AkPT1 (AkPT1TP-sGFP). Each bar represents the mean of three independent experiments. N.D., not detected. **b** Kinetic analysis. Apparent *K*_m_ values for bergaptol and DMAPP were calculated by nonlinear least squares method and reported as the means ± standard errors of three independent experiments. **c** pH dependency of B5ODT activity, measured in buffer containing 100 mM PIPES-KOH (pH 6.0 to 7.5) or 100 mM Tris-HCl (pH 7.5 to 9.0). Results are shown as relative to mean B5ODT activity at pH 9.0. Bars represent the mean of three independent experiments. **d** Divalent cation dependence of B5ODT activity. Reactions were performed in buffer containing MgCl_2_, MnCl_2_, CaCl_2_, CoCl_2_, or ZnCl_2_ or EDTA as a negative control. Results are shown as relative to mean B5ODT activity in the presence of Mg^2+^. Bars represent the mean of three independent experiments. N.D., not detected.

**Supplementary Fig. 12 Enzymatic reactions catalyzed by AkPT1**

(**a, b**) X8ODT, (**c, d**) B5OGT, and (**e, f**), X8OGT activities of AkPT1. Incubation with microsomes containing AkPT1TP-sGFP was used as a negative control. **(a, c, e)** UV chromatograms of reaction mixtures. Asterisks indicate molecules derived from impurities. Bol, bergaptol; Xol, xanthotoxol. (**b, d, f)** MS^2^ spectra from molecular ion peaks of enzymatic reaction products. Losses of 68 (**e**) and 136 (**d and f**) mass units were predicted to be derived from fragmentation due to the loss of *O*-dimethylallyl and *O*-geranyl moieties, respectively, attached to FC structures.

**Supplementary Fig. 13 Multiple likelihood-based phylogenetic tree of UbiA proteins**

The tree was constructed with 1,000 bootstrap tests based on a ClustalW multiple alignment of UbiA proteins, including CpPT1, CmiPT1a/b, and AkPT1. Bootstrap values are shown for nodes separating clades and for nodes between *C*-PTs and *O*-PTs in Rutaceae and Apiaceae. The bar represents an amino acid substitution rate per site of 0.50. The clades of primary metabolism-related proteins are marked in grey, whereas the clades of secondary metabolism-related PTs associated with VTE2-1, VTE2-2, and PPT are highlighted in orange, green, and blue, respectively, along with their aromatic substrates in each family, with *O*-PTs in Apiaceae and Rutaceae shown in red. Detailed information about input sequences is provided in Supplementary Table 3.

**Supplementary Fig. 14 FC profiles of pulps of ancestral *Citrus* species**

Total FC contents and the contents of *O*-geranylated FCs in pulps of papeda (*C. micrantha*, blue circle) and different varieties of pummelo (*C. grandis*, red circles), citron (*C. medica*, yellow circles), and mandarin (*C. reticulata*, orange circles) were plotted together with those of marsh grapefruit (grey circle). Quantitative data, expressed as means ± standard errors of three samples of Tahiti pumemlo and five samples each of all other varieties, have been reported^10^. Trace amounts of metabolites were set at zero for figure construction. Varieties of pummelo tested included Chandler, Deep Red, Kao Pan, Pink, Reinking, Seedless, and Tahiti pummelo; varities of citron tested included Corsican and Etrog citron; and varities of mandarin tested included Beauty, Cleopatra, Dancy, Fuzhu, Nan Feng Mi Chu, Owari Satsuma, San Hu Hong Chu, Shekwasha, Sunki, Wase Satsuma, and Willowleaf mandarin. Only the FC profiles of domesticated mandarins probably possessing pummelo-derived genomic segments were available. The FC molecules tested included 6’,7’-dihydroxybergamottin, 8-geranyloxypsoralen, bergamottin, bergapten, bergaptol, byakangelicin, byakangelicol, cnidicin, cnidilin, epoxybergamottin, heraclenin, heraclenol, imperatorin, isoimperatorin, isopimpinellin, oxypeucedanin, oxypeucedanin hydrate, phellopterin, psoralen, xanthotoxin, and xanthotoxol. Of these, 6’,7’-dihydroxybergamottin, 8-geranyloxypsoralen, bergamottin, and epoxybergamottin are *O*-geranylated FC derivatives, none of which was detectable in Corsican or Etrog citron. Buddha’s hand citron was not tested due to its pulp-less phenotype.

**Supplementary Table 1 Contigs classified into the UbiA superfamily**

Contigs in a grapefruit flavedo transcriptome dataset were classified into the UbiA superfamily by tblastn search with seven queries, *i.e.*, umbelliferone 8-*C*-geranyltransferase of lemon and ClPT1 and six proteins of sweet orange, probably orthologous to *Arabidopsis thaliana* UbiA PTs. These six proteins, VTE2-1, VTE2-2, PPT, ABC4, ATG4, and COX10, were functionally identified as being involved in the biosynthesis of tocopherol, plastoquinone, ubiquinone, phylloquinone, chlorophyll, and haem a, respectively. The homology with the seven queries in tblastn analysis and the functions predicted by homology for the leading query are shown for each UbiA family contig. UbiA family contigs showing low to moderate amino acid identities with any primary metabolism-related PT queries were designated as having unknown function and selected as candidate aromatic *O*-PTs. Contigs showing high homology with ClPT1 were also selected as candidates. NH, not hit. NA, not applicable. Further information on the queries (ClPT1, CsABC4, CsATG4, CsCOX10, CsPPT, CsVTE2-1, and CsVTE2-2) are shown in Fig. 4d, Supplementary Fig. 13 and Supplementary Table 3.

**Supplementary Table 2 *In silico* screening of a grapefruit leaf transcriptome**

A publicly available transcriptome prepared from grapefruit leaves that accumulate *O*-prenylated coumarins (transcriptome sample ID: UHJR in OneKP database) was screened with homologs for contigs considered candidates for aromatic *O*-PTs in Supplementary Table 1. Nucleotide identities were based on blastn searches using the candidate flavedo-derived contigs as queries (Supplementary Table 1). The flavedo-derived contigs with homologs showing over 95% nucleotide identity in the blastn search were retained as finer candidates.

**Supplementary Table 3 PT sequences used for *in silico* analyses**

UbiA PTs involved in (**a**) primary metabolism and (**b, c**) secondary metabolism in plants. (**b**) VTE2-1-related PTs and (**c**) VTE2-2- and PPT-related PTs.

**Supplementary Table 4 tBlastn search of an *Angelica archangelica* transcriptome using CpPT1 as a query**

Hits in an *A. archangelica* transcriptome dataset (sample ID: TQKZ in OneKP database) are listed according to their scores. CsVTE2-1 and AkPT1 were used as controls based on their taxonomic conservation. The threshold for an ortholog candidate of a query was set at 60% amino acid identity, based on previously reported amino acid identities between secondary metabolism-related UbiA PTs and their possible ancestors in a plant species. Hits meeting this criterion are highlighted in red. Three ortholog candidates are shown for AkPT1. Further information about CsVTE2-1 is available in Supplementary Table 3a.

**Supplementary Table 5 tBlastn search of a *Heracleum lanatum* transcriptome using CpPT1 as a query**

Hits in a *H. lanatum* transcriptome dataset (sample ID: CWYJ in OneKP database) are listed according to their scores. CsVTE2-1 and AkPT1 were used as controls based on their taxonomic conservation. The threshold for an ortholog candidate of a query was set at 60% amino acid identity, based on previously reported amino acid identities between secondary metabolism-related UbiA PTs and their possible ancestors in a plant species. Hits meeting this criterion are highlighted in red. Three ortholog candidates are shown for AkPT1. Further information about CsVTE2-1 is available in Supplementary Table 3a.

**Supplementary Table 6 tBlastn search of a grapefruit flavedo transcriptome using AkPT1 as a query**

Hits are listed according to their scores. DcVTE2-1 and CpPT1 were used as controls based on their taxonomic conservation. The threshold for an ortholog candidate of a query was set at 60% amino acid identity, based on previously reported amino acid identities between secondary metabolism-related UbiA PTs and their possible ancestors in a plant species. Hits meeting this criterion are highlighted in red. Contigs corresponding to CpPT1 are shown. Further information about DcVTE2-1 is available in Supplementary Table 3a.

**Supplementary table 7 Amino acid identities between bacterial and plant UbiA *O*-PTs for aromatics**

Amino acid identity of plant UbiA *O*-PTs and two bacterial UbiA *O*-PTs, CnqPT1 and AgqD, as determined by ClustalW multiple alignment. The N-terminal regions of CpPT1 (117 a.a.), CmiPT1a/b (116 a.a.), AkPT1 (108 a.a.), and CnqPT1 (39 a.a.) were truncated to adjust their polypeptide sequences for alignment with AgqD.

**Supplementary Table 8 List of PCR primers used in this study**

B=G/T/C, D=G/A/T, M=A/C, Y=C/T, R=A/G, K=G/T, H=A/C/T, V=C/A/G, and N=A/C/G/T for AkPT1_DGP primers.

## Notes

### Competing Interest Statement

The authors have declared no competing interest.

