## Supplemental Figures and Tables for "Parallel evolution of UbiA superfamily proteins into aromatic *O*-prenyltransferases in plants"

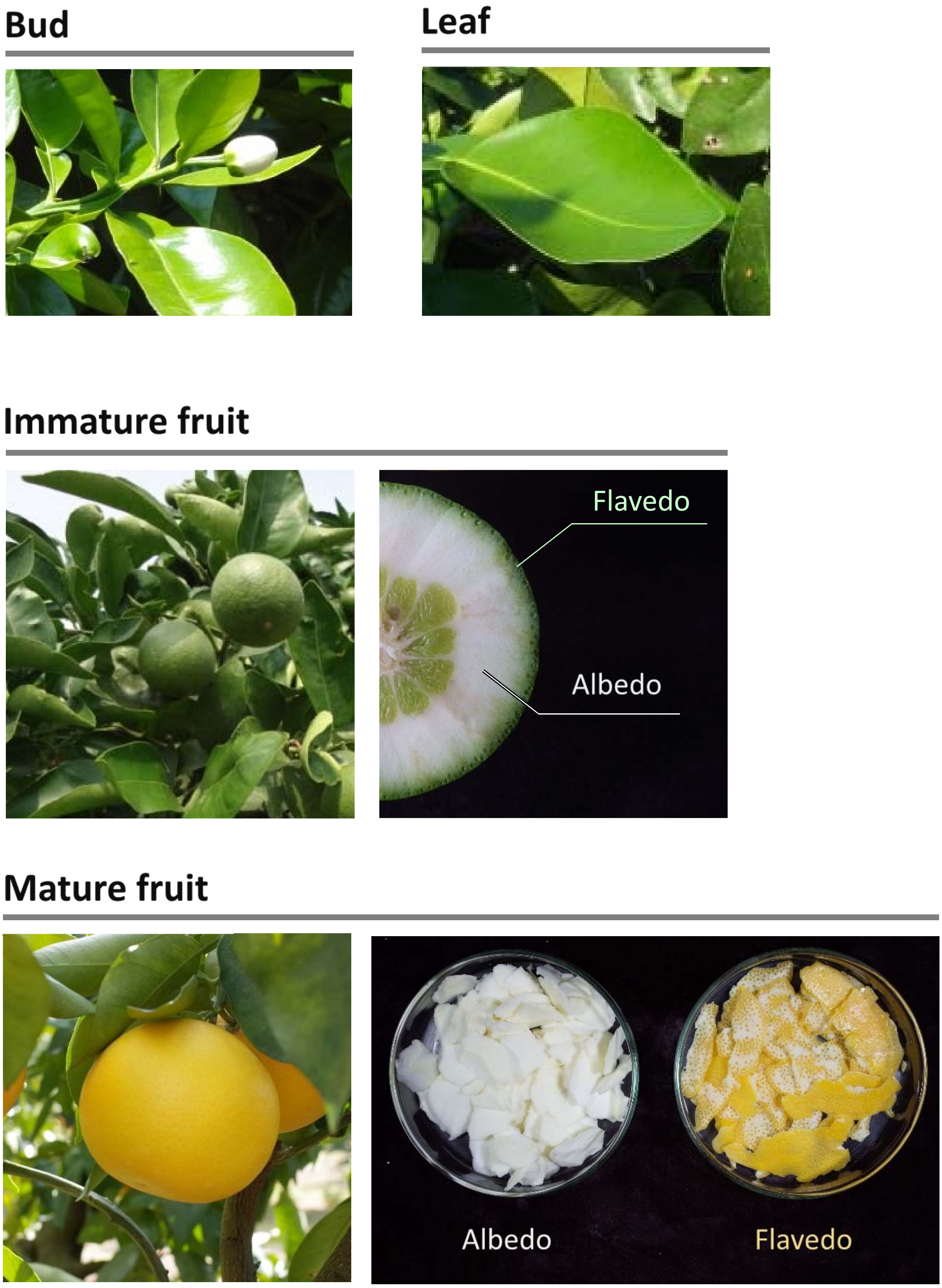


**Supplementary Fig. 1 Grapefruit samples used in this study**

Munakata *et al.*,

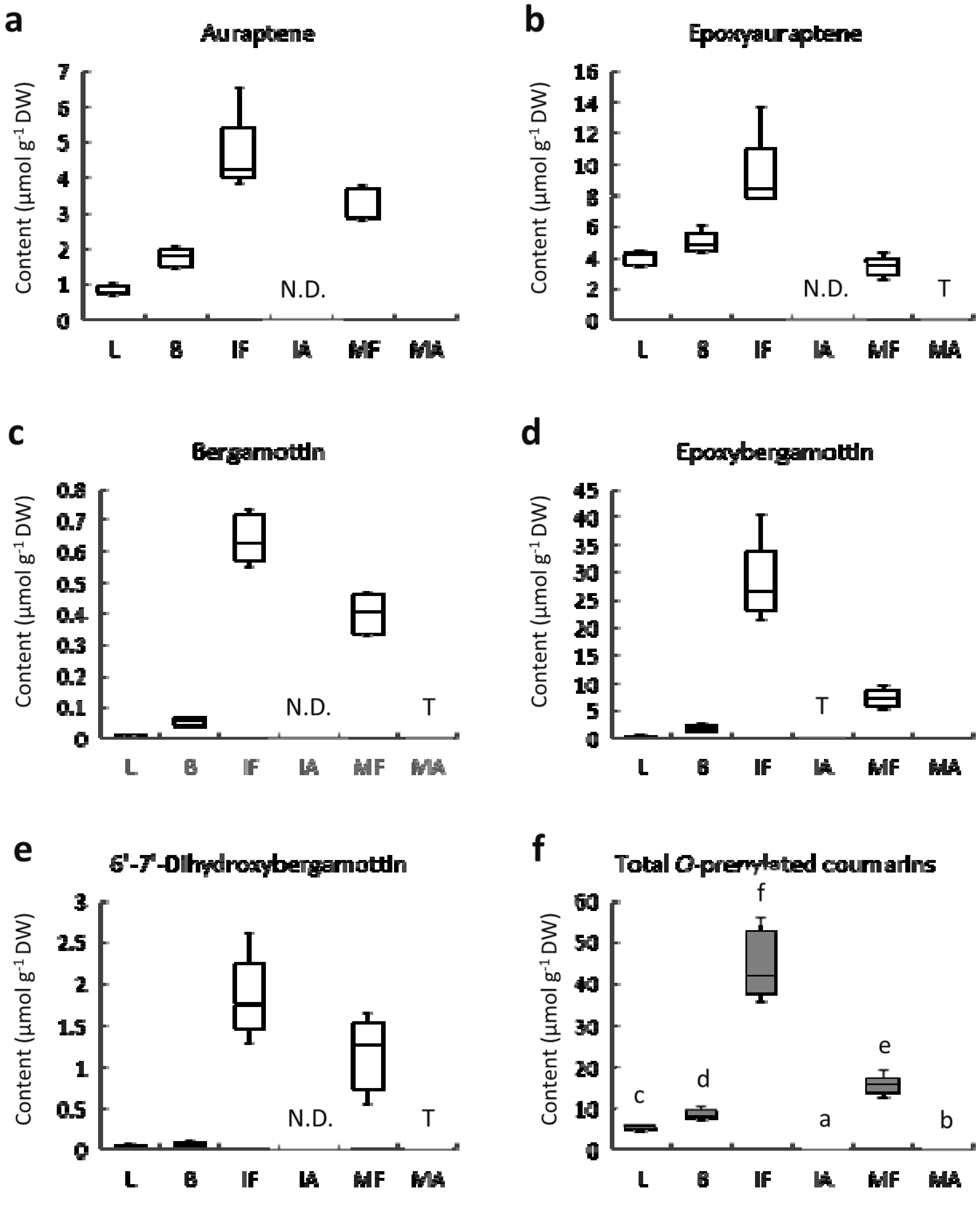


L : Leaves | IF : Immature fruit flavedo | MF : Mature fruit flavedo
B : Buds | IA : Immature fruit albedo | MA : Mature fruit albedo

**Supplementary Fig. 2 Coumarin contents of different grapefruit organs**

Munakata *et al.,*

a

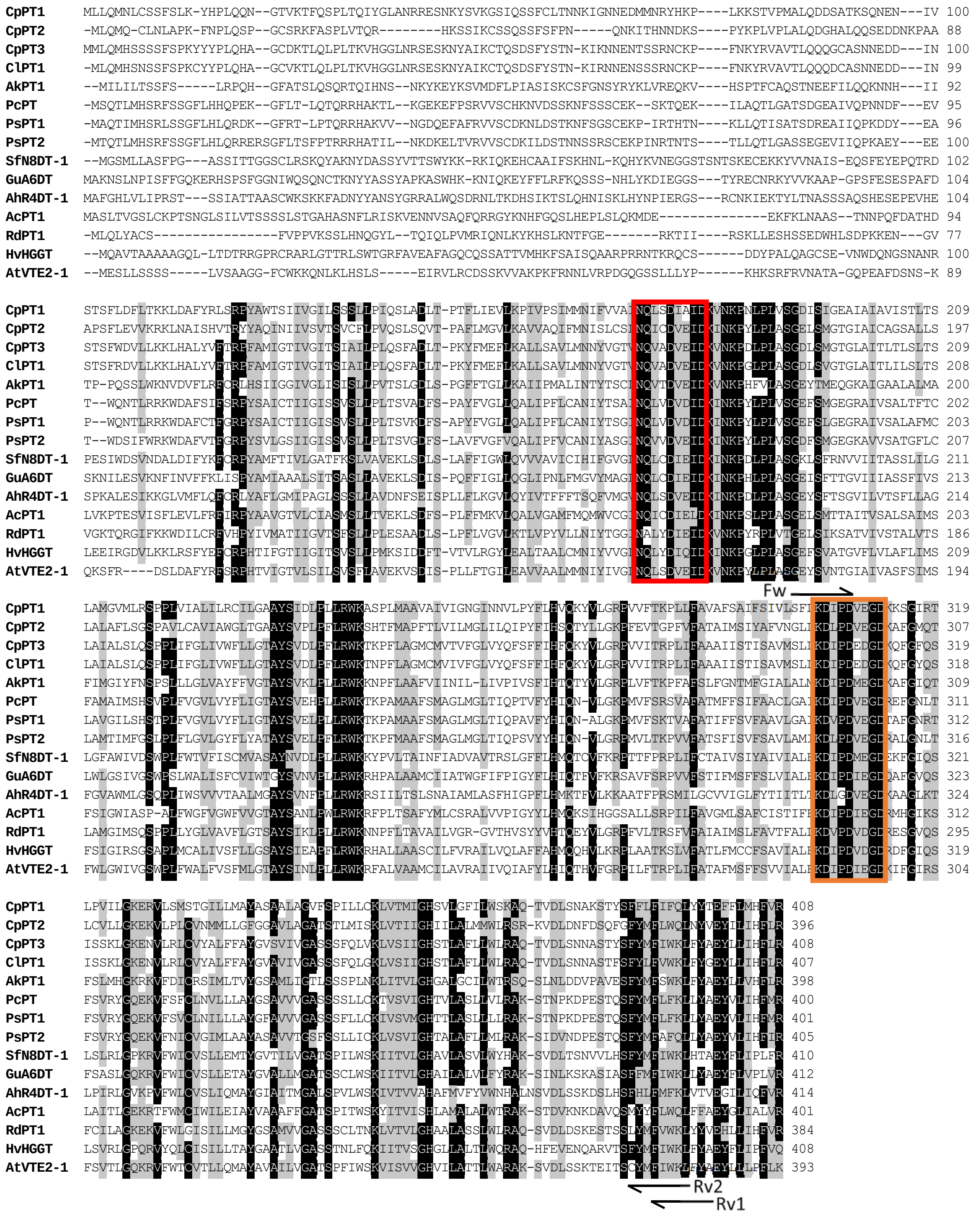


**Supplementary Fig. 3** ***In silico*** **analysis of CpPT1–3 and AkPT1 polypeptides**

Munakata *et al.,*

**b**

| Name | Score | TP | TP length |
|---|---|---|---|
| **CpPT1** | 0.519 | + | 49 |
| **CpPT2** | 0.561 | + | 39 |
| **CpPT3** | 0.465 | - | 50 |
| **AkPT1** | 0.501 | + | 46 |

**c**

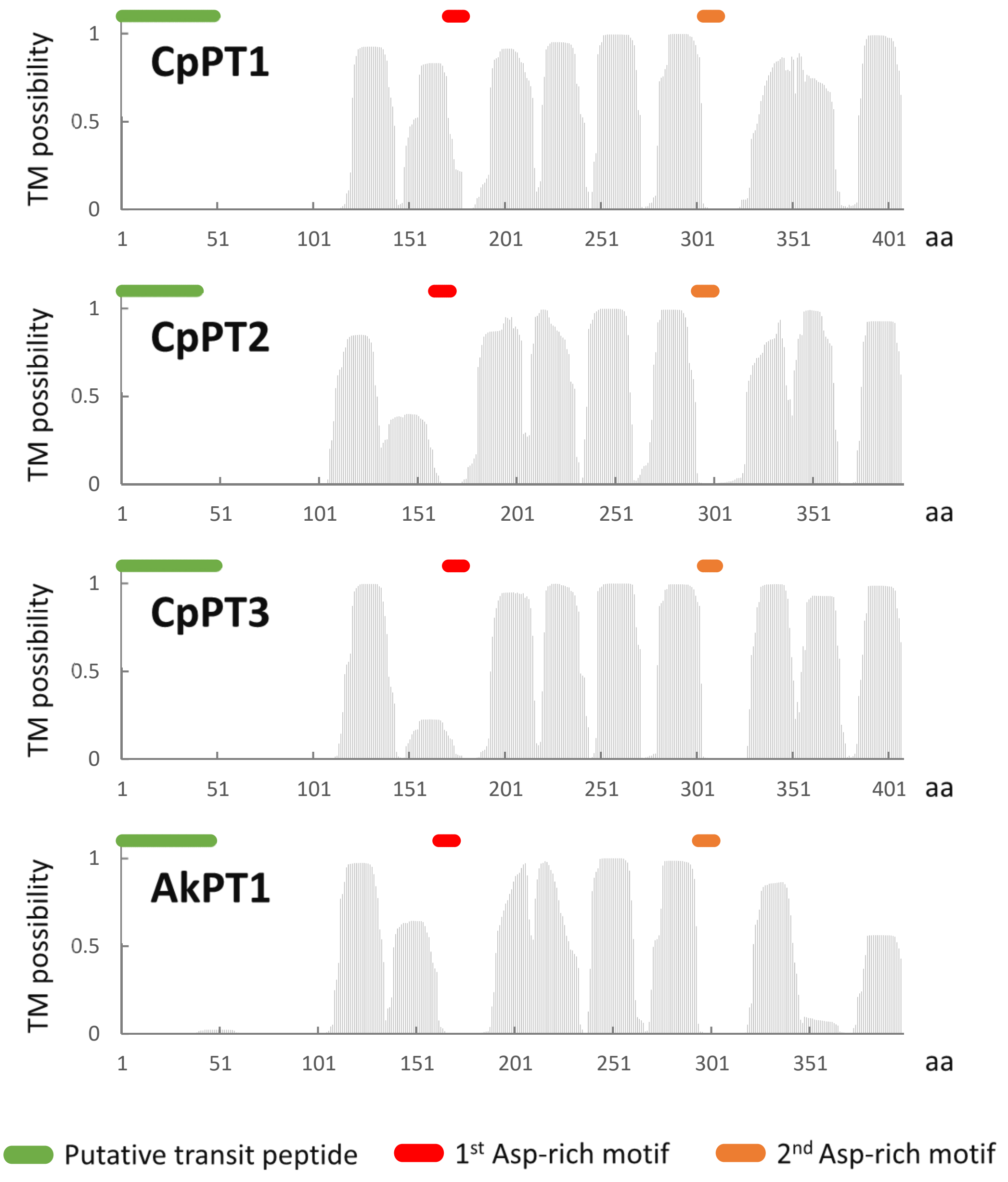


**Supplementary Fig. 3** ***In silico*** **analysis of CpPT1–3 and AkPT1 polypeptides -*continued***

Munakata *et al.,*

**a**

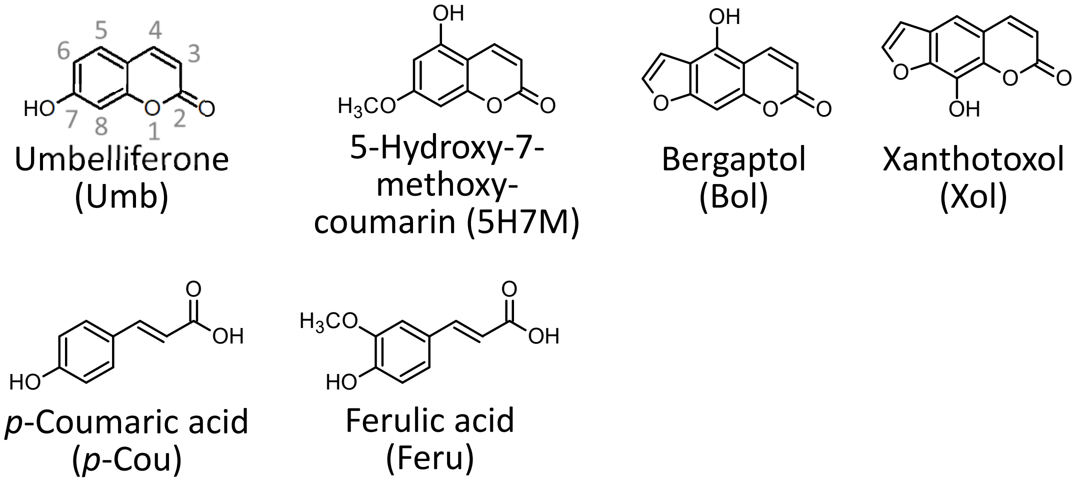


**b**

| PT | Substrate pair | | | | | | |
|---|---|---|---|---|---|---|---|
| | **Prenyl donor** | **Prenyl acceptor** | | | | | |
| | | **Umb** | **5H7M** | **Bol** | **Xol** | ***p*-Cou** | **Feru** |
| **CpPT2** | **DMAPP** | N.D. | N.D. | N.D. | N.D. | N.D. | N.D. |
| | **GPP** | N.D. | N.D. | N.D. | N.D. | N.D. | N.D. |
| **CpPT3** | **DMAPP** | N.D. | N.D. | N.D. | N.D. | N.D. | N.D. |
| | **GPP** | 8GU, 6GU? (trace) | Trace | Trace | N.D. | Trace | N.D. |

**Supplementary Fig. 4 Biochemical screening of CpPT2 and CpPT3**

Munakata *et al.,*

c

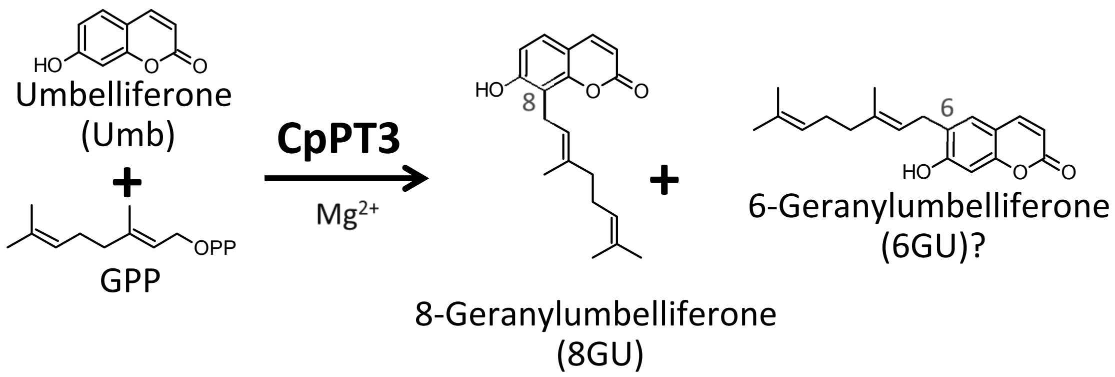


d

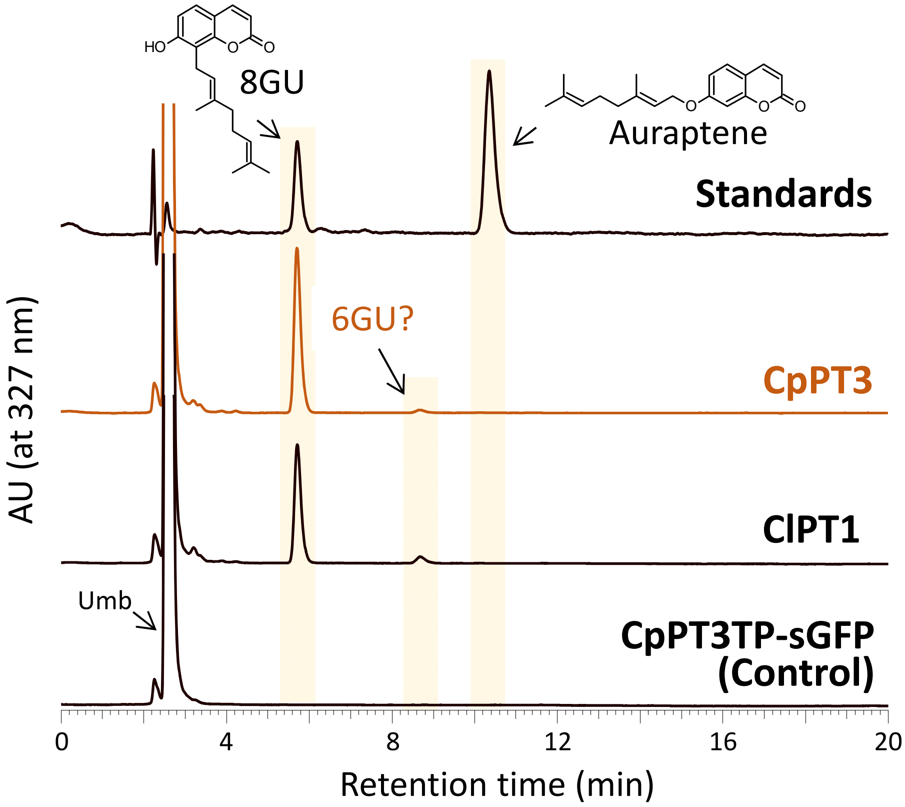


**Supplementary Fig. 4 Biochemical screening of CpPT2 and CpPT3 -*continued***

Munakata *et al.,*

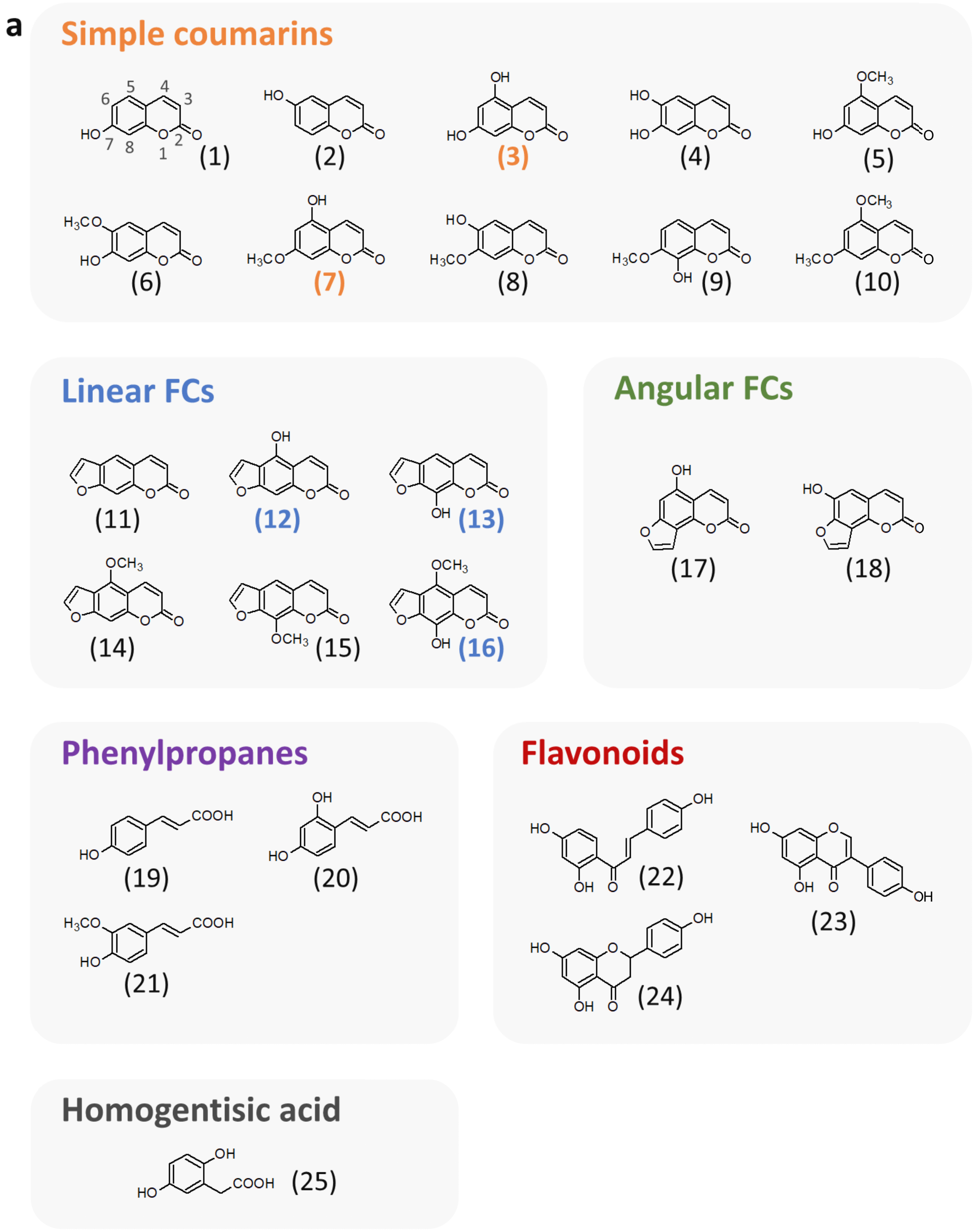


**Supplementary Fig. 5 Enzymatic reactions catalyzed by CpPT1**

Munakata *et al.,*

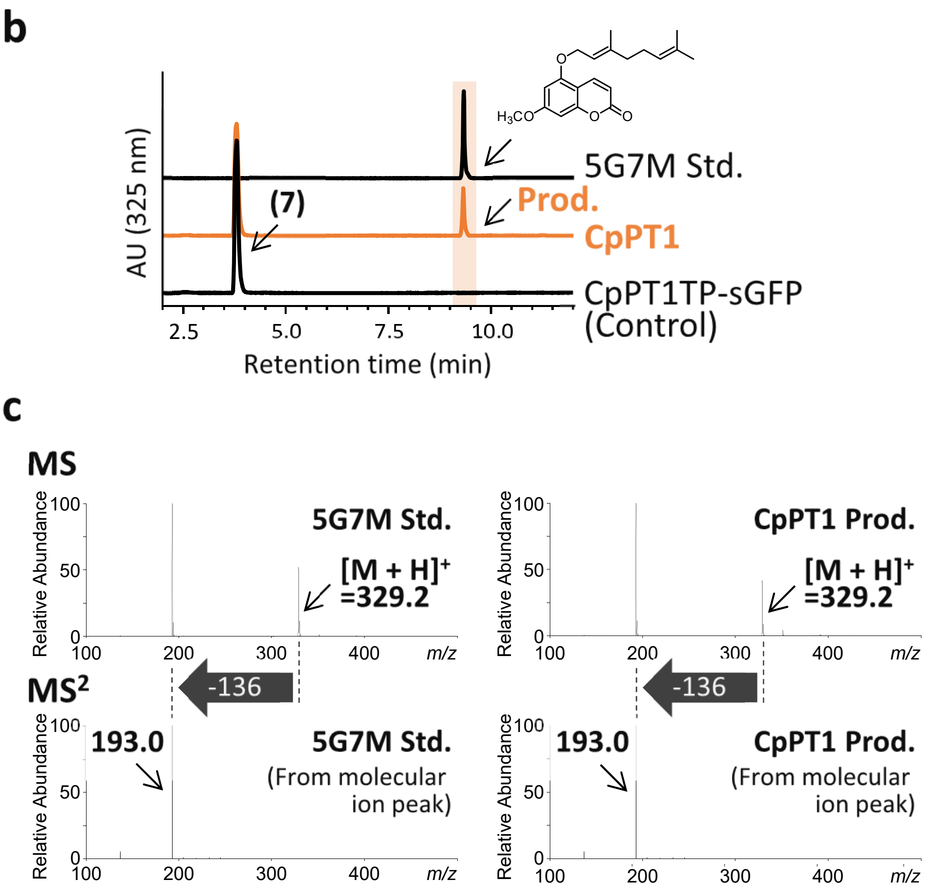


**Supplementary Fig. 5 Enzymatic reactions catalyzed by CpPT1** ***-continued***

Munakata *et al.,*

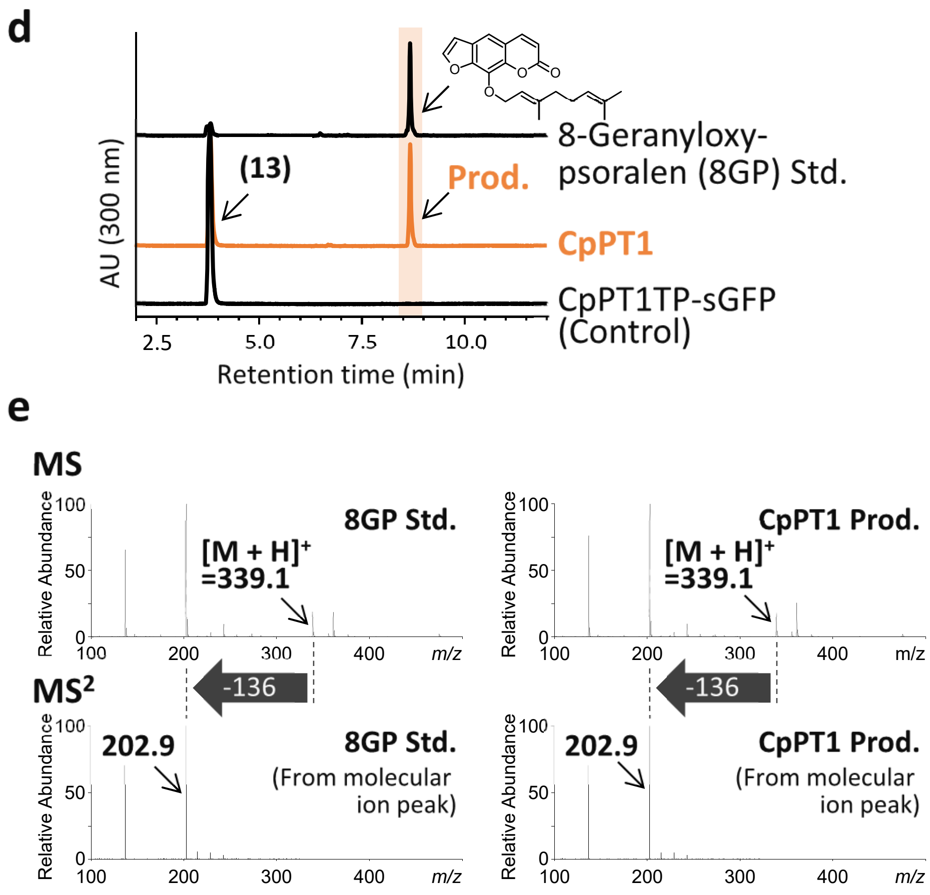


**Supplementary Fig. 5 Enzymatic reactions catalyzed by CpPT1 *-continued***

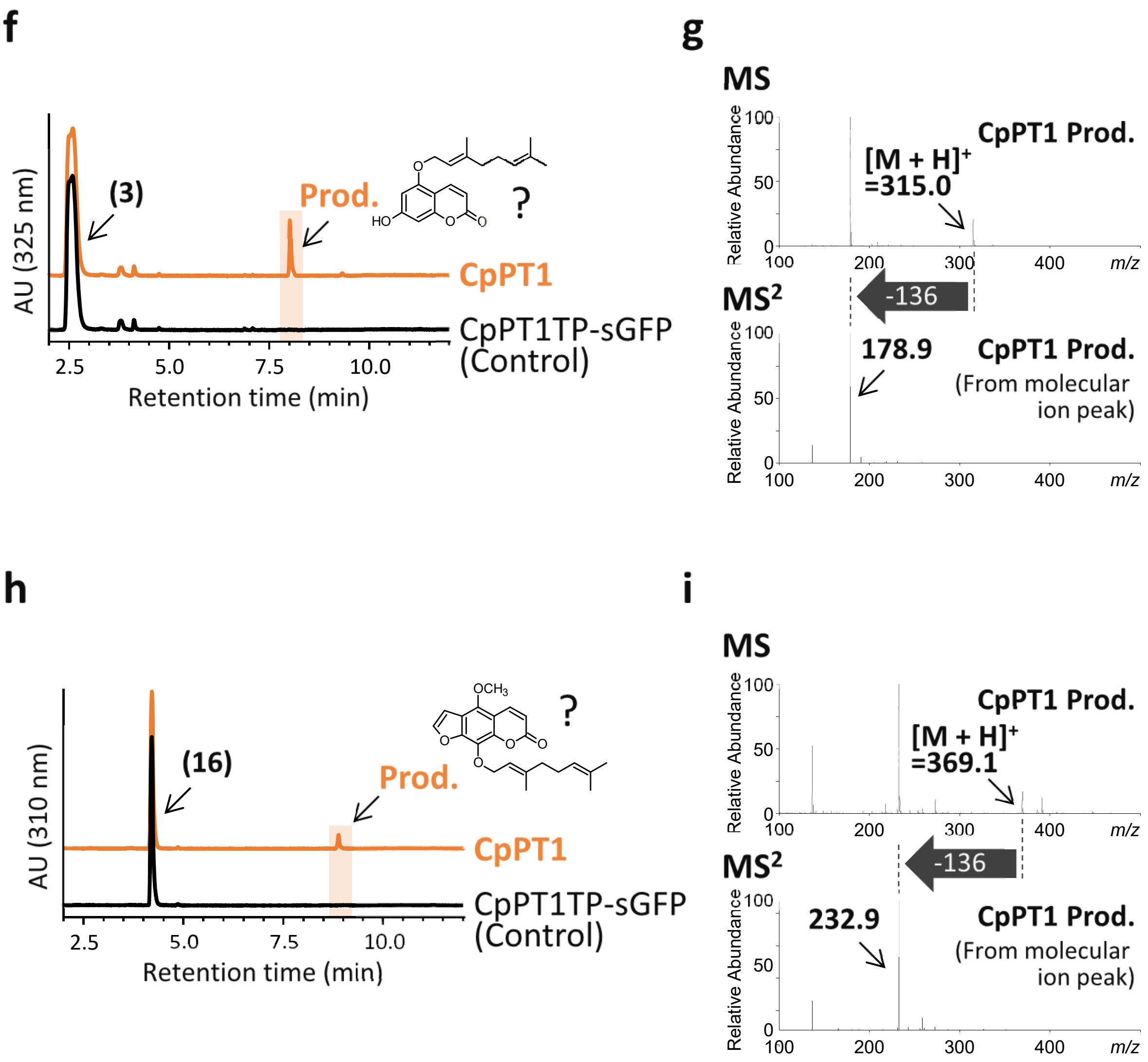


**Supplementary Fig. 5 Enzymatic reactions catalyzed by CpPT1 *-continued***

**a**

**b**

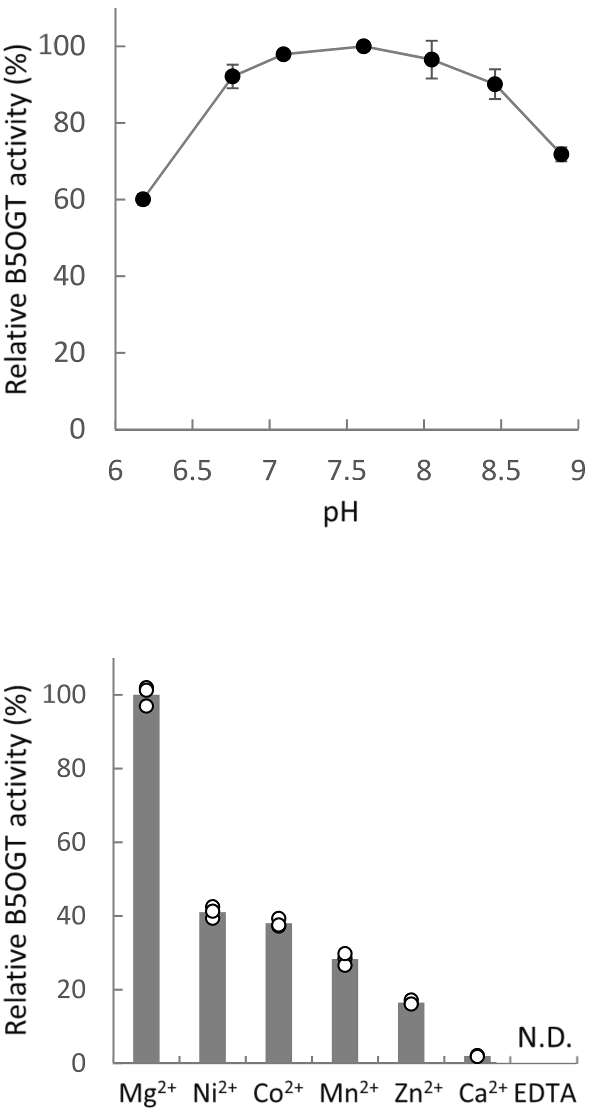


**Supplementary Fig. 6 Properties of the B5OGT activity of CpPT1**

Munakata *et al.,*

**a**

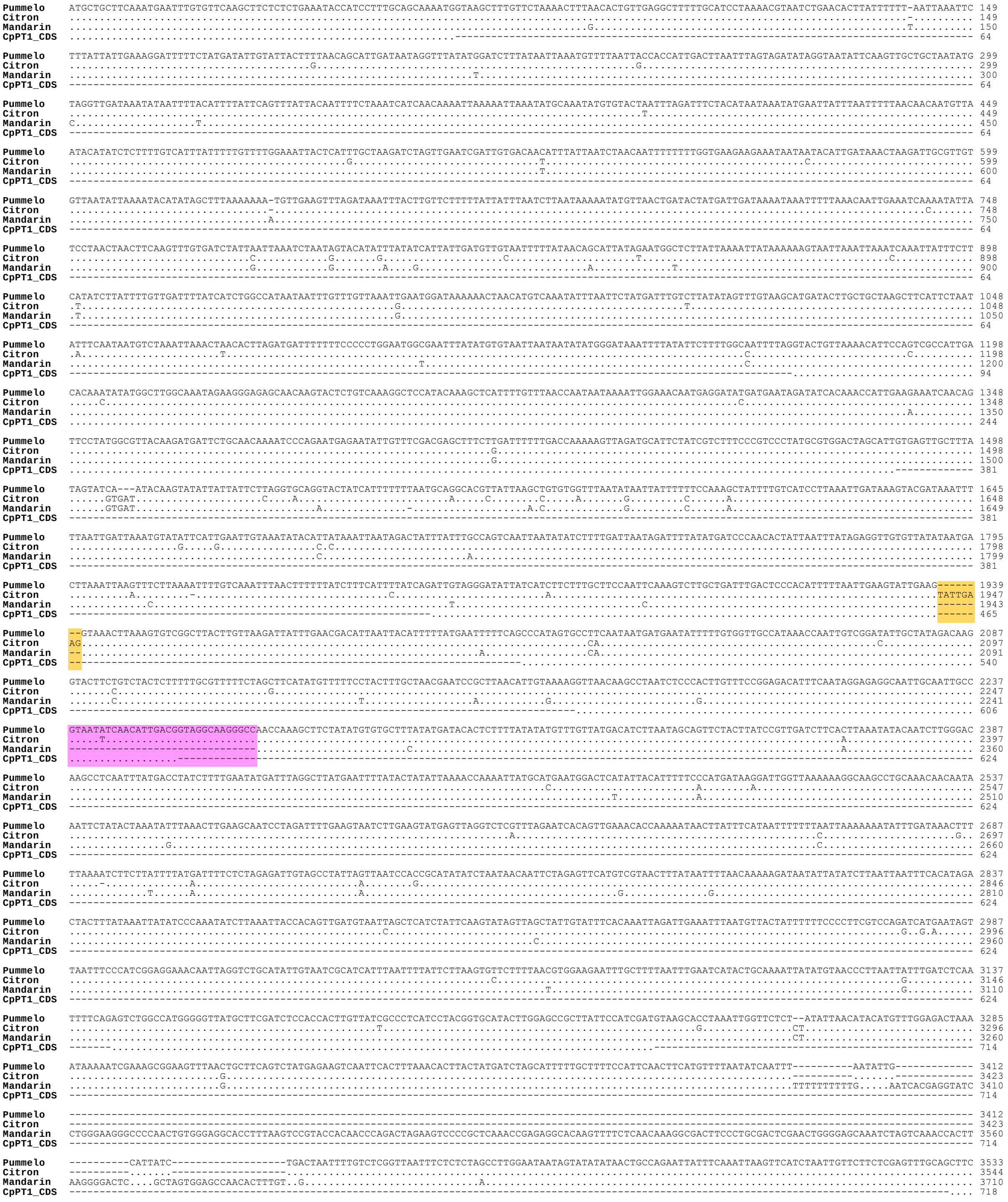


#### Supplementary Fig. 7 Structures of *CpPT1* gene orthologs in *Citrus*

Munakata *et al.,*

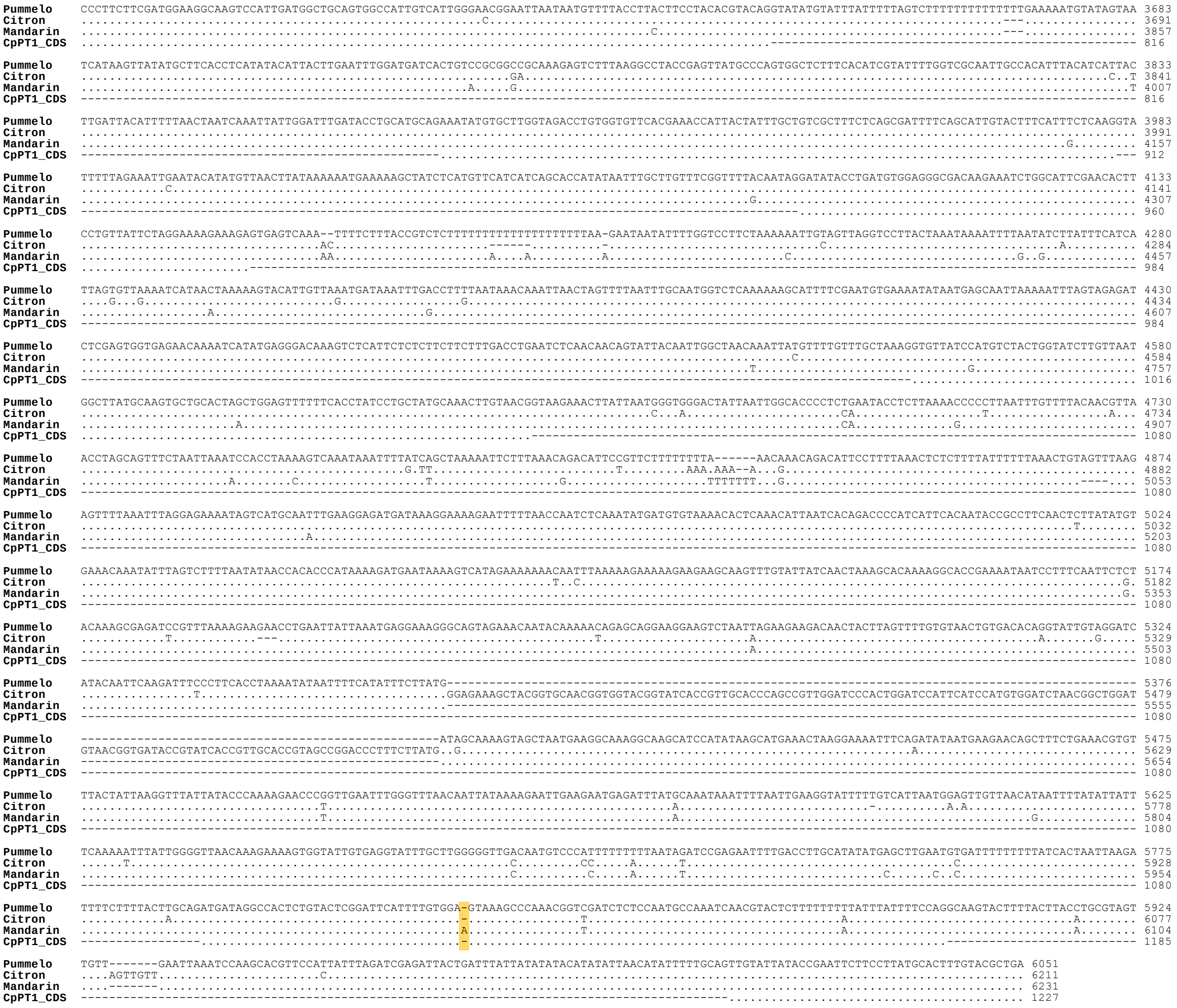

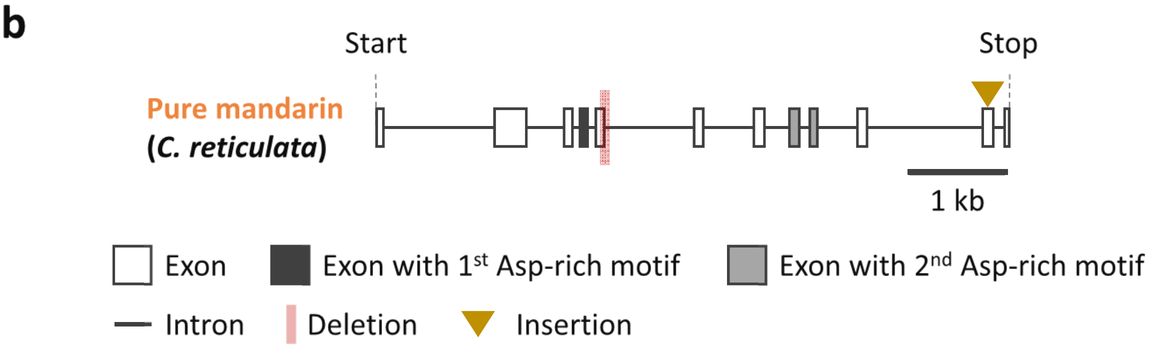


**Supplementary Fig. 7 Structures of *CpPT1* gene orthologs in *Citrus -continued***

Munakata *et al.*,

a

```
CmiPT1a  MLLQMNLCSSFSLKYHPLQQNGTVKTFHSPLTQIYGLANRRESNKYSVKGSTQSSFCLTNNKIGNNEDMMNRYHKPLK-STVPRALQDDSATKSQNENIV 99
CmiPT1b  ...........................Q..................................................-..................... 99
CpPT1    ...........................Q.......................I..........................K...M................ 100

CmiPT1a  WTNFLDFLTKKLDAFYRLSRPYAWTSIIVGILSSSLLPIQSLADLTPTFLIEVLKPIVPTIMMNIFVVAINQLSDIAIDKVNKPNLPLVSGDISIGEAIA 199
CmiPT1b  .................................................................................................... 199
CpPT1    S.S........................................................S........................................ 200

CmiPT1a  IAVISTLTSLAMGVMLRSPPLVIALILRCILGAAYSIDLPLLRWKASPLMAAVAIVIGNGINNVLPYFLHVQKYVLGRPVVFTKPLLFAVAFSAIFSIVL 299
CmiPT1b  .................................................................................................... 299
CpPT1    .................................................................................................... 300

CmiPT1a  SFLKDIPDVEGDKKSGIRTLPVILGKERVLSMSTGILLMAYASAALAGVFSPILLCKLVTMIGHSVLGFILWSKAQTVDLSNAKSTYSFFIFIFQLYYTE 399
CmiPT1b  .................................................................................................... 399
CpPT1    ..........................................................................................L......... 400

CmiPT1a  FFLMHFVR 407
CmiPT1b  ........ 407
CpPT1    ........ 408
```

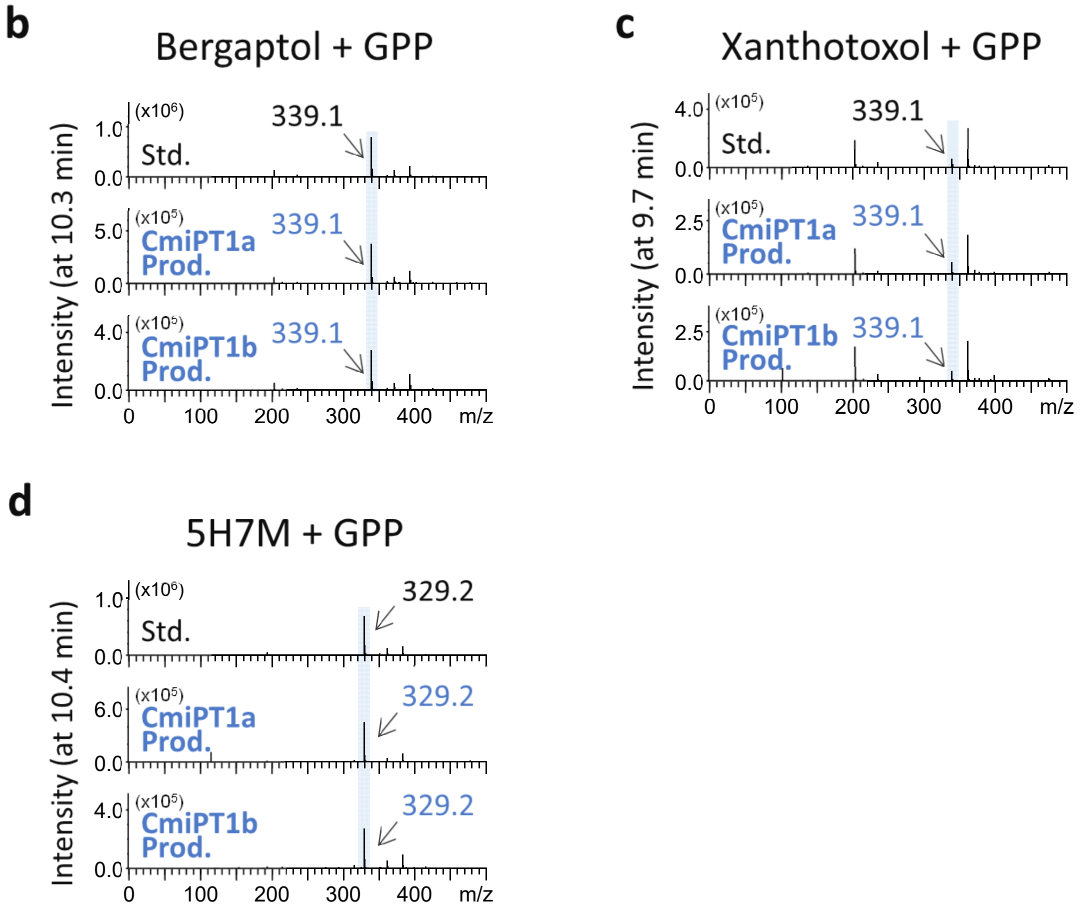


**Supplementary Fig. 8 Isolation of *CpPT1* orthologs from papeda**

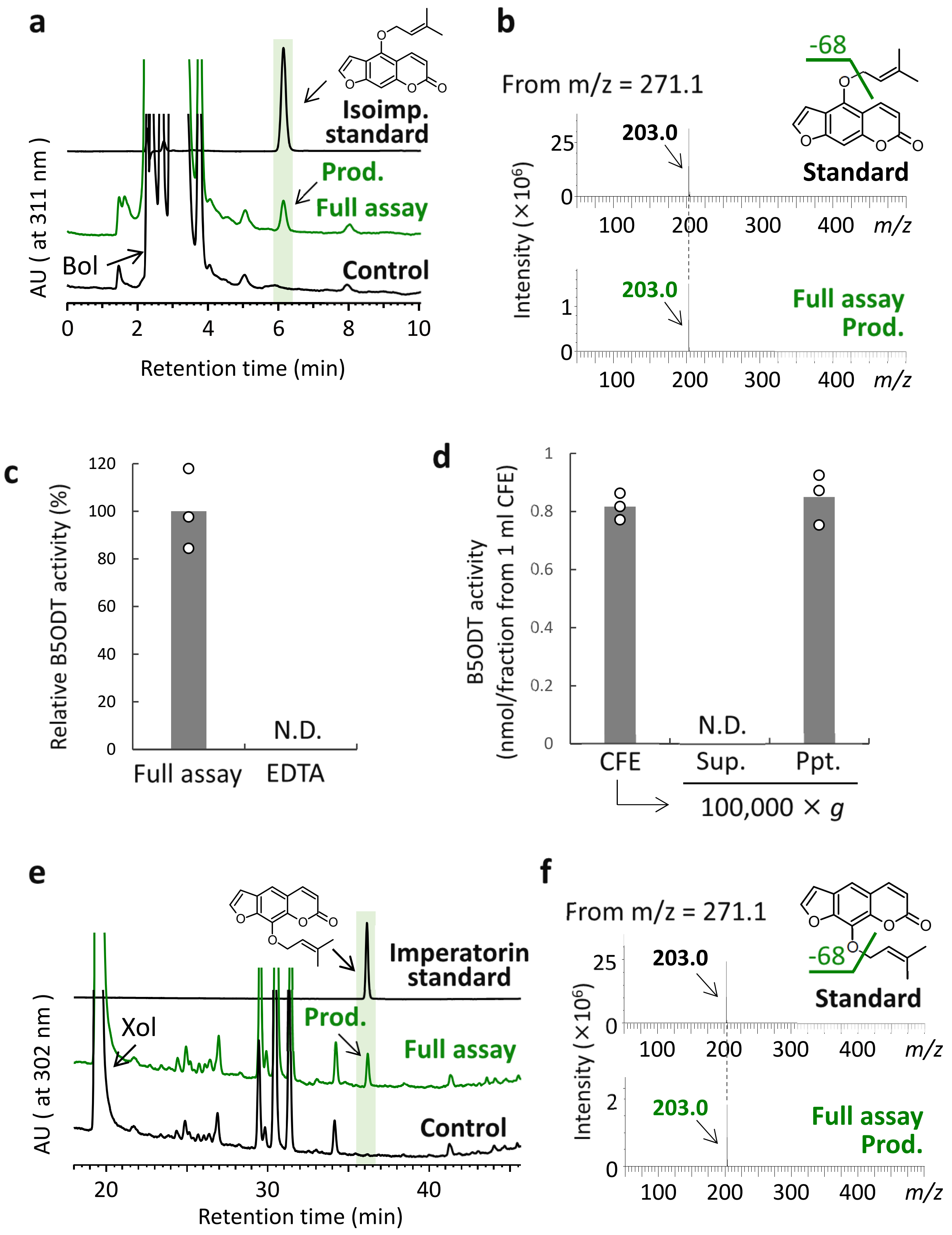


**Supplementary Fig. 9 Furanocoumarin *O*-dimethylallyltransferase activities of *Angelica keiskei* microsomes**

Munakata *et al.*,

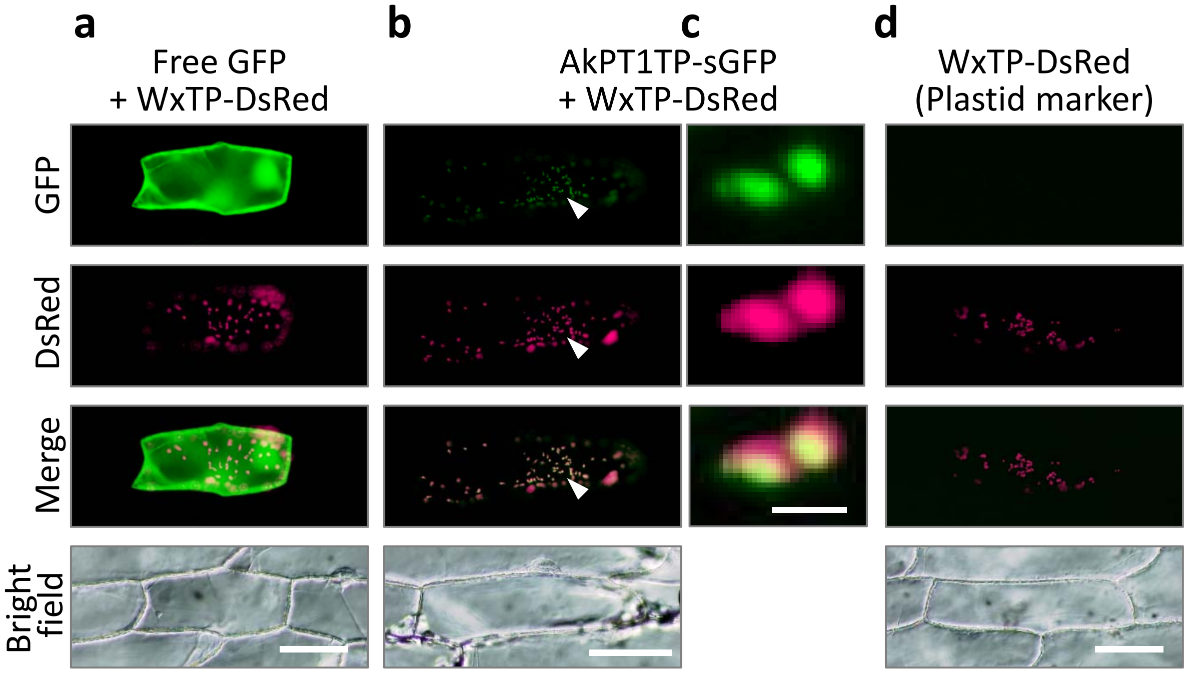


**Supplementary Fig. 10 Subcellular localization of AkPT1**

Munakata *et al.*,

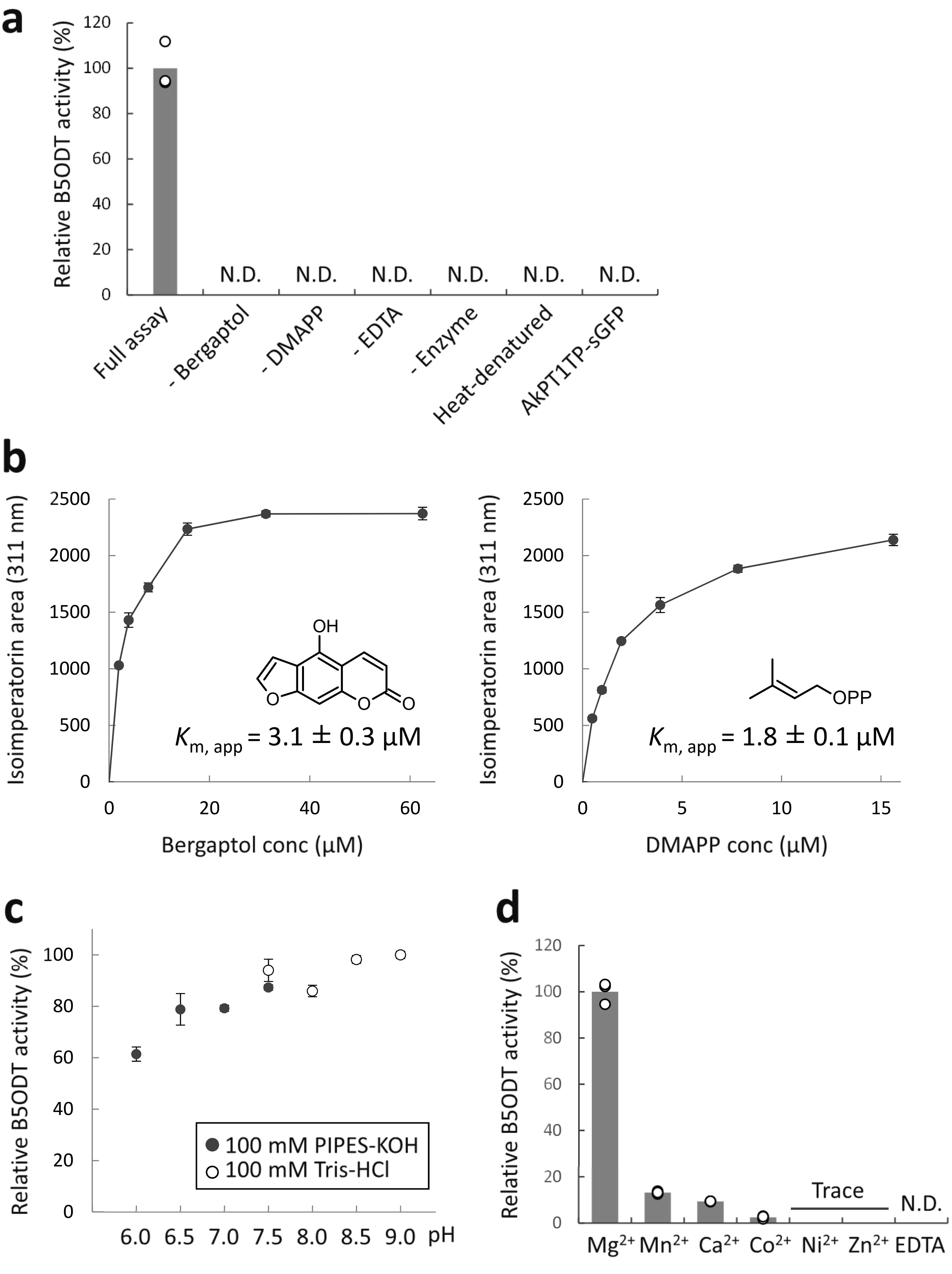


**Supplementary Fig. 11 Characterization of the B5ODT activity of AkPT1**

Munakata *et al.,*

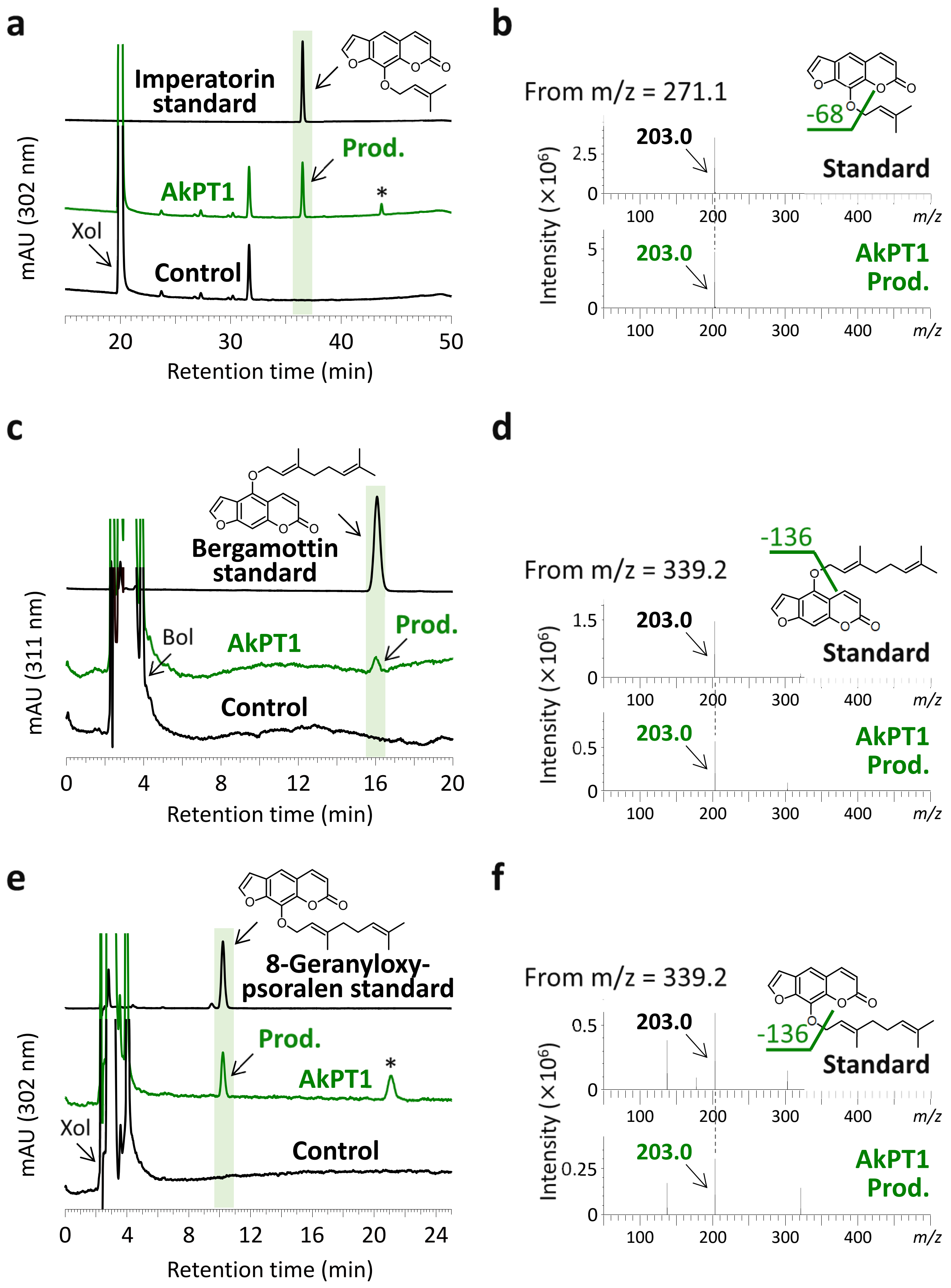


**Supplementary Fig. 12 Enzymatic reactions catalyzed by AkPT1**

Munakata *et al.,*

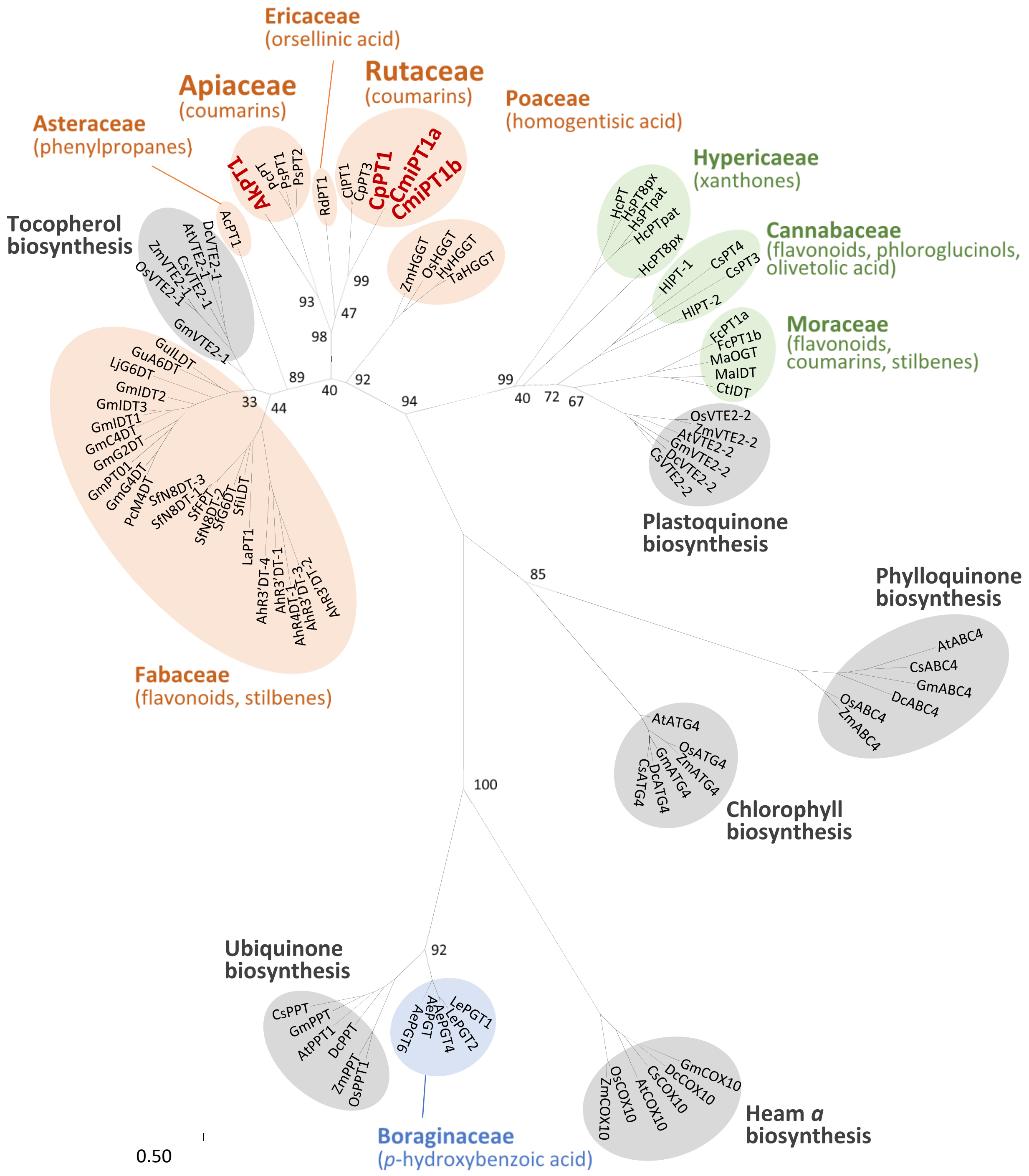


**Supplementary Fig. 13 Multiple likelihood-based phylogenetic tree of UbiA proteins**

Munakata *et al.*,

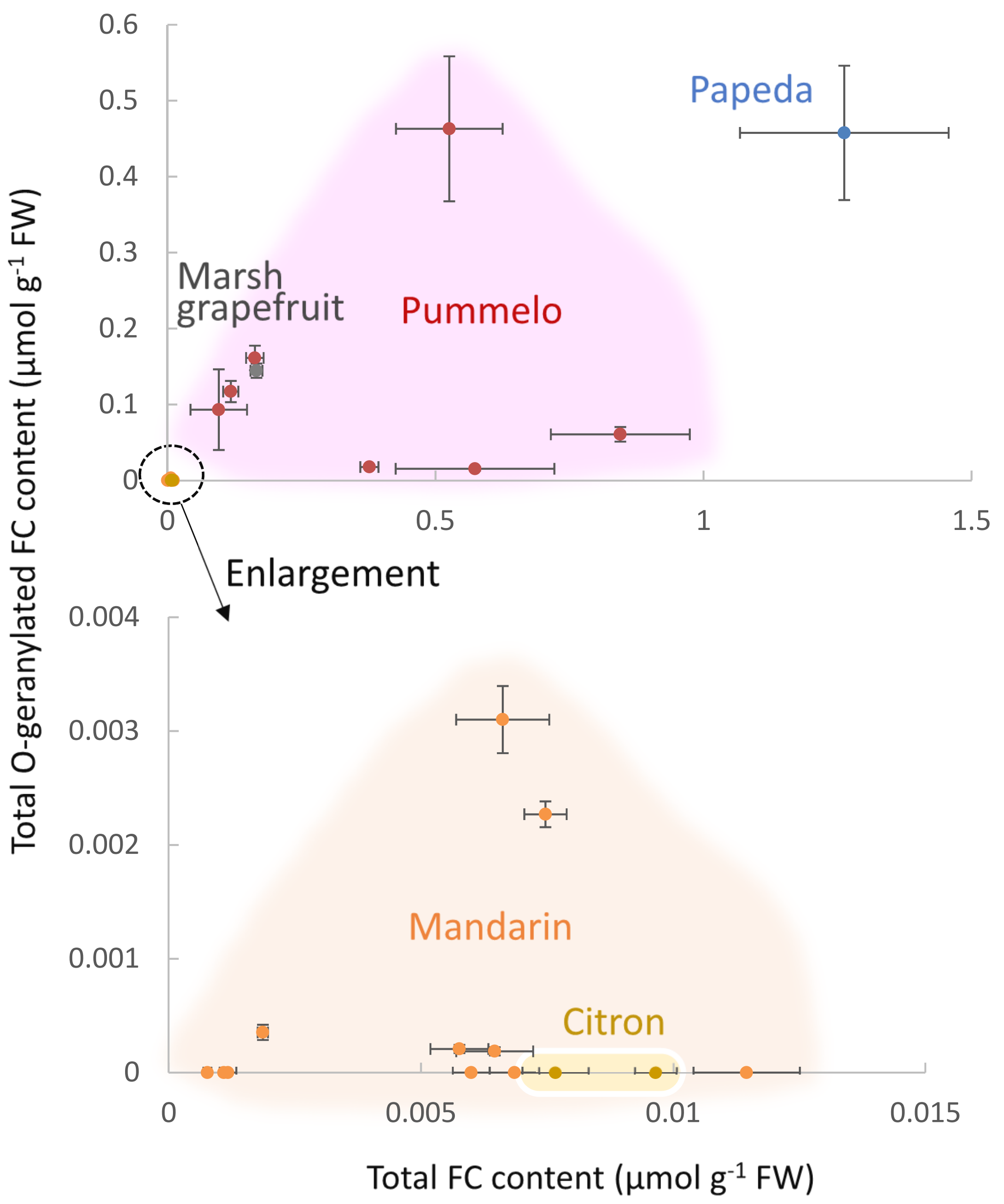


**Supplementary Fig. 14 FC profiles of pulps of ancestral *Citrus* species**

Munakata *et al.*,

### Supplementary Table 1 Contigs classified into the UbiA superfamily

| Contig | Query | | | | | | | Tophit (of all) | Amino acid identity (%) | Tophit (wo CIPT) | Amino acid identity (%) | Predicted function |
|---|---|---|---|---|---|---|---|---|---|---|---|---|
| | VTE2-1 | VTE2-2 | PPT | ABC4 | ATG4 | COX10 | CIPT | | | | | |
| c6071_g1_i1 | NH | NH | 97 | NH | 28 | NH | NH | PPT | 97 | PPT | 97 | PPT |
| c11028_g1_i1 | NH | NH | NH | 97 | NH | NH | NH | ABC4 | 97 | ABC4 | 97 | ABC4 |
| c11028_g1_i2 | NH | NH | NH | 98 | NH | NH | NH | ABC4 | 98 | ABC4 | 98 | ABC4 |
| c12976_g1_i1 | 35 | NH | NH | NH | NH | NH | 40 | CIPT | 40 | VTE2-1 | 35 | Unknown |
| c12976_g1_i2 | NH | NH | NH | NH | NH | NH | 41 | CIPT | 41 | NH | NA | Unknown |
| c13428_g1_i1 | 32 | 26 | NH | NH | NH | NH | 42 | CIPT | 42 | VTE2-1 | 32 | Unknown |
| c13428_g1_i2 | 40 | 29 | NH | NH | NH | NH | 49 | CIPT | 49 | VTE2-1 | 40 | Unknown |
| c13877_g1_i1 | 45 | 28 | NH | NH | 26 | NH | 51 | CIPT | 51 | VTE2-1 | 45 | Unknown |
| c13877_g1_i2 | 45 | 28 | NH | NH | 26 | NH | 51 | CIPT | 51 | VTE2-1 | 45 | Unknown |
| c14187_g1_i1 | 35 | 99 | NH | 27 | 27 | NH | 29 | VTE2-2 | 99 | VTE2-2 | 99 | VTE2-2 |
| c15601_g1_i1 | 39 | 32 | NH | NH | 26 | NH | 41 | CIPT | 41 | VTE2-1 | 39 | Unknown |
| c15601_g1_i2 | 39 | 30 | NH | NH | 26 | NH | 44 | CIPT | 44 | VTE2-1 | 39 | Unknown |
| c16042_g1_i1 | 48 | 29 | NH | NH | 23 | NH | 95 | CIPT | 95 | VTE2-1 | 48 | CIPT1 |
| c17212_g1_i1 | 95 | 34 | NH | NH | 23 | NH | 44 | VTE2-1 | 95 | VTE2-1 | 95 | VTE2-1 |
| c17212_g1_i2 | 90 | 37 | NH | NH | 29 | NH | 43 | VTE2-1 | 90 | VTE2-1 | 90 | VTE2-1 |
| c17212_g1_i3 | 98 | 38 | NH | NH | 25 | NH | 48 | VTE2-1 | 98 | VTE2-1 | 98 | VTE2-1 |
| c18070_g1_i1 | NH | NH | NH | NH | 25 | 99 | NH | COX10 | 99 | COX10 | 99 | COX10 |
| c18070_g1_i2 | NH | NH | NH | NH | 25 | 99 | NH | COX10 | 99 | COX10 | 99 | COX10 |
| c18070_g1_i3 | NH | NH | NH | NH | NH | 94 | NH | COX10 | 94 | COX10 | 94 | COX10 |
| c18070_g1_i4 | NH | NH | NH | NH | 25 | 99 | NH | COX10 | 99 | COX10 | 99 | COX10 |
| c18070_g1_i5 | NH | NH | NH | NH | NH | 100 | NH | COX10 | 100 | COX10 | 100 | COX10 |
| c18955_g2_i1 | NH | 48 | NH | NH | 100 | NH | 36 | ATG4 | 100 | ATG4 | 100 | ATG4 |
| c18955_g3_i1 | 20 | NH | 30 | NH | 100 | NH | NH | ATG4 | 100 | ATG4 | 100 | ATG4 |
| c18955_g3_i2 | 24 | 22 | 30 | 25 | 100 | NH | NH | ATG4 | 100 | ATG4 | 100 | ATG4 |
| c19000_g3_i1 | 34 | 30 | NH | NH | NH | NH | 50 | CIPT | 50 | VTE2-1 | 34 | Unknown |
| c19000_g3_i2 | 29 | NH | NH | NH | NH | NH | 49 | CIPT | 49 | VTE2-1 | 29 | Unknown |
| c21508_g1_i1 | 49 | 35 | NH | NH | 30 | NH | 48 | VTE2-1 | 49 | VTE2-1 | 49 | Unknown |
| c22985_g1_i1 | 39 | 26 | NH | NH | NH | NH | 47 | CIPT | 47 | VTE2-1 | 39 | Unknown |
| c23037_g1_i1 | NH | NH | NH | 97 | NH | NH | NH | ABC4 | 97 | ABC4 | 97 | ABC4 |
| c23764_g1_i1 | NH | NH | NH | NH | NH | NH | 39 | CIPT | 39 | NH | NA | Unknown |
| c28572_g1_i1 | 44 | 35 | NH | NH | 32 | NH | 56 | CIPT | 56 | VTE2-1 | 44 | Unknown |
| c32272_g1_i1 | 37 | NH | NH | NH | NH | NH | 58 | CIPT | 58 | VTE2-1 | 37 | Unknown |
| c35003_g1_i1 | 35 | NH | NH | NH | NH | NH | 96 | CIPT | 96 | VTE2-1 | 35 | CIPT1 |

100% NA

Munakata *et al.*,

#### Supplementary Table 2 *In silico* screening of a grapefruit leaf transcriptome

| Query | Grapefruit Leaf RNA-seq | |
|---|---|---|
| | Top hit | Nucleotide identity (%) |
| c12976_g1_i1 | No hit | Not applicable |
| c12976_g1_i2 | No hit | Not applicable |
| c13428_g1_i1 | UHJR_scaffold_2029749 | 100 |
| c13428_g1_i2 | UHJR_scaffold_2029749 | 100 |
| c13877_g1_i1 | UHJR_scaffold_2055721 | 99 |
| c13877_g1_i2 | UHJR_scaffold_2055721 | 99 |
| c15601_g1_i1 | UHJR_scaffold_2000939 | 99 |
| c15601_g1_i2 | UHJR_scaffold_2000939 | 96 |
| c16042_g1_i1 | UHJR_scaffold_2003469 | 99 |
| c19000_g3_i1 | UHJR_scaffold_2020849 | 100 |
| c19000_g3_i2 | UHJR_scaffold_2020849 | 100 |
| c21508_g1_i1 | UHJR_scaffold_2011569 | 99 |
| c22985_g1_i1 | UHJR_scaffold_2006570 | 100 |
| c23764_g1_i1 | No hit | Not applicable |
| c28572_g1_i1 | No hit | Not applicable |
| c32272_g1_i1 | UHJR_scaffold_2011569 | 100 |
| c35003_g1_i1 | UHJR_scaffold_2048908 | 100 |

#### Supplementary Table 3 PT sequences used for *in silico* analyses

**(a)**

| PT in primary metabolism | Plant species | Accession No. |
|---|---|---|
| **ABC4s in Phylloquinone biosynthesis** | | |
| AtABC4 | *Arabidopsis thaliana* | NP_001117518.1 |
| CsABC4 | *Citrus sinensis* | XP_024948946.1 |
| DcABC4 | *Daucus carota* | XP_017230268.1 |
| GmABC4 | *Glycine max* | XP_003532605.1 |
| OsABC4 | *Oryza sativa* | NP_001049226.1 |
| ZmABC4 | *Zea mays* | NP_001152170.1 |
| **ATGs in Chrolophyll biosynthesis** | | |
| AtATG4 | *Arabidopsis thaliana* | NP_190750.1 |
| CsATG4 | *Citrus sinensis* | XP_006481064.1 |
| DcATG4 | *Daucus carota* | XP_017243789.1 |
| GmATG4 | *Glycine max* | NP_001239633.1 |
| OsATG4 | *Oryza sativa* | ABO31092.1 |
| ZmATG4 | *Zea mays* | NP_001142204.1 |
| **COX10s in Heam *a* biosynthesis** | | |
| AtCOX10 | *Arabidopsis thaliana* | NP_566019.1 |
| CsCOX10 | *Citrus sinensis* | XP_006464403.1 |
| DcCOX10 | *Daucus carota* | XP_017227264.1 |
| GmCOX10 | *Glycine max* | XP_003556552.1 |
| OsCOX10 | *Oryza sativa* | EEC70799.1 |
| ZmCOX10 | *Zea mays* | AFW89544.1 |
| **PPTs in Ubiquinone biosynthesis** | | |
| AtPPT1 | *Arabidopsis thaliana* | NP_567688 |
| CsPPT | *Citrus sinensis* | XP_015387502.1 |
| DcPPT | *Daucus carota* | XP_017222930.1 |
| GmPPT | *Glycine max* | XP_006602724.1 |
| OsPPT1 | *Oryza sativa* | BAE96574.1 |
| ZmPPT | *Zea mays* | NP_001148558.1 |
| **VTE2-1s in Tocopherol biosynthesis** | | |
| AtVTE2-1 | *Arabidopsis thaliana* | NP_849984.1 |
| CsVTE2-1 | *Citrus sinensis* | XP_006474206.1 |
| DcVTE2-1 | *Daucus carota* | XP_017253952.1 |
| GmVTE2-1 | *Glycine max* | NP_001241496.1 |
| TaVTE2-1 | *Triticum aestivum* | ABB70123.1 |
| ZmVTE2-1 | *Zea mays* | ACG45339.1 |
| **VTE2-2s in Plastoquinone biosynthesis** | | |
| AtVTE2-2 | *Arabidopsis thaliana* | NP_001154609.1 |
| CsVTE2-2 | *Citrus sinensis* | XP_006481679.1 |
| DcVTE2-2 | *Daucus carota* | XP_017246707.1 |
| GmVTE2-2 | *Glycine max* | NP_001237900 |
| OsVTE2-2 | *Oryza sativa* | XP_015646905.1 |
| ZmVTE2-2 | *Zea mays* | NP_001146703.1 |

#### Supplementary Table 3 PT sequences used for *in silico* analyses *-continued*

**(b)**

| VTE2-1-related PTs | Plant species | Accession No. |
|---|---|---|
| **Apiaceae** | | |
| PcPT | *Petroselinum crispum* | BAO31627.1 |
| PsPT1 | *Pastinaca sativa* | AJW31563.1 |
| PsPT2 | *Pastinaca sativa* | AJW31564.1 |
| **Asteraceae** | | |
| AcPT1 | *Artemisia capillaris* | BBG56301.1 |
| **Ericaceae** | | |
| RdPT1 | *Rhododendron dauricum* | LC381857 |
| **Fabaceae** | | |
| AhR3'DT-1 | *Arachis hypogaea* | AQM74173.1 |
| AhR3'DT-2 | *Arachis hypogaea* | AQM74174.1 |
| AhR3'DT-3 | *Arachis hypogaea* | AQM74175.1 |
| AhR3'DT-4 | *Arachis hypogaea* | AQM74176.1 |
| AhR4DT-1 | *Arachis hypogaea* | AQM74172.1 |
| GmC4DT | *Glycine max* | BAW32575.1 |
| GmG2DT | *Glycine max* | BAW32578.1 |
| GmG4DT | *Glycine max* | NP_001235990 |
| GmIDT1 | *Glycine max* | BAW32576.1 |
| GmIDT2 | *Glycine max* | BAW32577.1 |
| GmIDT3 | *Glycine max* | XP_014618511.1 |
| GmPT01 | *Glycine max* | KRH76147.1 |
| GuA6DT | *Glycyrrhiza uralensis* | AIT11912.1 |
| GuILDT | *Glycyrrhiza uralensis* | AMR58303.1 |
| LaPT1 | *Lupinus albus* | AER35706.1 |
| LjG6DT | *Lotus japonicus* | ARV85585.1 |
| PcM4DT | *Psoralea corylifolia* | AYV64464.1 |
| SfFPT | *Sophora flavescens* | AHA36633.1 |
| SfG6DT | *Sophora flavescens* | BAK52291.1 |
| SfiLDT | *Sophora flavescens* | BAK52290.1 |
| SfN8DT-1 | *Sophora flavescens* | BAG12671.1 |
| SfN8DT-2 | *Sophora flavescens* | BAG12673.1 |
| SfN8DT-3 | *Sophora flavescens* | BAK52289.1 |
| **Poaceae** | | |
| HvHGGT | *Hordeum vulgare* | AAP43911.1 |
| OsHGGT | *Oryza sativa* | AAP43913.1 |
| TaHGGT | *Triticum aestivum* | AAP43912.1 |
| ZmHGGT | *Zea mays* | XP_008659772.1 |
| **Rutaceae** | | |
| ClPT1 | *Citrus limon* | BAP27988.1 |

#### Supplementary Table 3 PT sequences used for *in silico* analyses *-continued*

**(c)**

| VTE2-2- and PPT-related PTs | Plant species | Accession No. |
|---|---|---|
| **Cannabaceae** | | |
| CsPT3 | *Cannabis sativa* | DAC76713.1 |
| CsPT4 | *Cannabis sativa* | DAC76710.1 |
| HlPT-1 | *Humulus lupulus* | BAJ61049.1 |
| HlPT-2 | *Humulus lupulus* | AJD80255.1 |
| **Hypericaceae** | | |
| HcPT | *Hypericum calycinum* | ALD84371 |
| HcPT8pat | *Hypericum calycinum* | AZK16227.1 |
| HcPT8px | *Hypericum calycinum* | AZK16226.1 |
| HsPT8pat | *Hypericum sampsonii* | AZK16225.1 |
| HsPT8px | *Hypericum sampsonii* | AZK16224.1 |
| **Moraceae** | | |
| CtIDT | *Cudrania tricuspidata* | AJD80983.1 |
| FcPT1a | *Ficus carica* | LC369744 |
| FcPT1b | *Ficus carica* | LC369745 |
| MaIDT | *Morus alba* | AJD80982.1 |
| MaOGT | *Morus alba* | AXN57307.1 |
| **Boraginaceae** | | |
| AePGT | *Arnebia euchroma* | ABD59796.2 |
| AePGT4 | *Arnebia euchroma* | ANC67957.1 |
| AePGT6 | *Arnebia euchroma* | ANC67959.1 |
| LePGT1 | *Lithospermum erythrorhizon* | BAB84122.1 |
| LePGT2 | *Lithospermum erythrorhizon* | BAB84123.1 |

### Supplementary Table 4 tBlastn search of an *Angelica archangelica* transcriptome using CpPT1 as a query

| Query | Hit | Score (bits) | E-vlaue | Amino acid identity (%) |
|---|---|---|---|---|
| CpPT1 | scaffold_2002205 | 249 | 2.44E-78 | 38 |
| | scaffold_2014187 | 248 | 1.52E-77 | 44 |
| | scaffold_2060869 | 234 | 2.35E-72 | 41 |
| | scaffold_2001569 | 171 | 7.80E-51 | 40 |
| | scaffold_2012634 | 149 | 1.47E-42 | 49 |
| | scaffold_2001568 | 129 | 1.78E-41 | 44 |
| | scaffold_2008065 | 127 | 1.34E-33 | 38 |
| | scaffold_2052354 | 114 | 1.79E-30 | 47 |
| | scaffold_2061747 | 121 | 4.70E-30 | 29 |
| | scaffold_2005436 | 107 | 1.64E-26 | 46 |
| | scaffold_2005435 | 98.2 | 2.74E-24 | 47 |
| | scaffold_2007401 | 94.7 | 4.58E-23 | 43 |
| | scaffold_2012456 | 77 | 2.19E-16 | 41 |
| | scaffold_2012457 | 77 | 6.09E-16 | 41 |
| | scaffold_2012413 | 54.7 | 9.50E-09 | 30 |
| | scaffold_2053505 | 51.2 | 1.10E-07 | 42 |
| | scaffold_2019790 | 37.7 | 4.33E-04 | 51 |
| | scaffold_2019789 | 37 | 6.88E-04 | 51 |
| | scaffold_2013732 | 38.5 | 0.007 | 23 |
| | scaffold_2013731 | 38.5 | 0.007 | 23 |
| | scaffold_2023841 | 28.5 | 0.9 | 37 |

| Query | Hit | Score (bits) | E-vlaue | Amino acid identity (%) |
|---|---|---|---|---|
| CsVTE2-1 | scaffold_2002205 | 540 | 0 | 68 |
| | scaffold_2060869 | 268 | 7.86E-86 | 44 |
| | scaffold_2014187 | 268 | 2.83E-85 | 43 |
| | scaffold_2012634 | 189 | 6.06E-58 | 60 |
| | scaffold_2001569 | 167 | 3.35E-49 | 42 |
| | scaffold_2052354 | 161 | 1.52E-48 | 53 |
| | scaffold_2061747 | 162 | 1.4E-44 | 34 |
| | scaffold_2001568 | 116 | 9.97E-40 | 44 |
| | scaffold_2005436 | 127 | 2.35E-34 | 47 |
| | scaffold_2008065 | 126 | 2.66E-33 | 43 |
| | scaffold_2005435 | 100 | 4.36E-25 | 48 |
| | scaffold_2007401 | 93.6 | 8.21E-23 | 46 |
| | scaffold_2012413 | 82.8 | 1.67E-18 | 48 |
| | scaffold_2012456 | 81.6 | 4.57E-18 | 45 |
| | scaffold_2012457 | 81.6 | 1.35E-17 | 45 |
| | scaffold_2053505 | 67.4 | 3.17E-13 | 42 |
| | scaffold_2023841 | 43.1 | 4.4E-06 | 51 |
| | scaffold_2013732 | 47.8 | 9.56E-06 | 23 |
| | scaffold_2013731 | 47.8 | 1.01E-05 | 23 |
| | scaffold_2009436 | 44.3 | 1.42E-05 | 42 |
| | scaffold_2019790 | 34.3 | 0.006 | 51 |
| | scaffold_2019789 | 33.5 | 0.011 | 48 |
| | scaffold_2004116 | 32.3 | 0.5 | 23 |
| | scaffold_2006080 | 32 | 0.78 | 26 |
| AkPT1 | scaffold_2001569 | 318 | 3.39E-108 | 79 |
| | scaffold_2012413 | 277 | 5.79E-93 | 87 |
| | scaffold_2001568 | 240 | 2.74E-91 | 92 |

Munakata *et al.,*

#### Supplementary Table 5 tBlastn search of a *Heracleum lanatum* transcriptome using CpPT1 as a query

| Query | Hit | Score (bits) | E-vlaue | Amino acid identity (%) | Query | Hit | Score (bits) | E-vlaue | Amino acid identity (%) |
|---|---|---|---|---|---|---|---|---|---|
| CpPT1 | scaffold_2070463 | 251 | 8.61E-79 | 43 | CsVTE2-1 | scaffold_2070463 | 521 | 0 | 68 |
| | scaffold_2015756 | 251 | 1.27E-78 | 40 | | scaffold_2069621 | 279 | 2.21E-90 | 45 |
| | scaffold_2069621 | 237 | 4.25E-74 | 38 | | scaffold_2006947 | 275 | 2.27E-87 | 42 |
| | scaffold_2006947 | 240 | 1.03E-73 | 38 | | scaffold_2015756 | 258 | 1.64E-81 | 42 |
| | scaffold_2002855 | 119 | 4.06E-31 | 32 | | scaffold_2071546 | 157 | 5.92E-43 | 35 |
| | scaffold_2071546 | 120 | 1.14E-29 | 30 | | scaffold_2002855 | 134 | 8.16E-37 | 39 |
| | scaffold_2061773 | 70.1 | 4.68E-14 | 36 | | scaffold_2061773 | 85.5 | 1.65E-19 | 46 |
| | scaffold_2061767 | 67.4 | 4.12E-13 | 35 | | scaffold_2061767 | 84.3 | 3.32E-19 | 47 |
| | scaffold_2008481 | 43.9 | 1.35E-04 | 35 | | scaffold_2002856 | 67.4 | 3.55E-13 | 35 |
| | scaffold_2008482 | 43.5 | 1.52E-04 | 36 | | scaffold_2027734 | 47 | 2.52E-07 | 55 |
| | scaffold_2002856 | 40 | 8.26E-04 | 33 | | scaffold_2071595 | 47 | 1.76E-05 | 23 |
| | scaffold_2027734 | 36.6 | 0.001 | 39 | | scaffold_2008481 | 46.2 | 2.74E-05 | 39 |
| | scaffold_2071595 | 37 | 0.029 | 24 | | scaffold_2001060 | 44.3 | 9.52E-05 | 57 |
| | scaffold_2054803 | 34.3 | 0.035 | 28 | | scaffold_2008482 | 41.6 | 4.96E-04 | 40 |
| | | | | | | scaffold_2047558 | 37.4 | 0.002 | 100 |
| | | | | | | scaffold_2065322 | 27.7 | 0.57 | 40 |
| | | | | | AkPT1 | scaffold_2002855 | 338 | 6.57E-116 | 73 |
| | | | | | | scaffold_2002856 | 161 | 3.75E-48 | 67 |
| | | | | | | scaffold_2061767 | 152 | 2.22E-44 | 84 |

Munakata *et al.,*

#### Supplementary Table 6 tBlastn search of a grapefruit flavedo transcriptome using AkPT1 as a query

| Query | Hit | Score (bits) | E-vlaue | Amino acid identity (%) |
|---|---|---|---|---|
| AkPT1 | c17212_g1_i1 | 272 | 3.00E-73 | 45 |
| | c15601_g1_i2 | 254 | 7.00E-68 | 43 |
| | c16042_g1_i1 | 228 | 7.00E-60 | 47 |
| | c15601_g1_i1 | 175 | 5.00E-44 | 44 |
| | c13877_g1_i2 | 164 | 2.00E-40 | 45 |
| | c13877_g1_i1 | 163 | 3.00E-40 | 45 |
| | c13428_g1_i2 | 129 | 4.00E-30 | 33 |
| | c28572_g1_i1 | 126 | 3.00E-29 | 43 |
| | c22985_g1_i1 | 121 | 1.00E-27 | 32 |
| | c17212_g1_i2 | 117 | 2.00E-26 | 45 |
| | c14187_g1_i1 | 108 | 1.00E-23 | 27 |
| | c13428_g1_i1 | 88 | 1.00E-17 | 28 |
| | c17212_g1_i3 | 86 | 7.00E-17 | 48 |
| | c32272_g1_i1 | 85 | 1.00E-16 | 42 |
| | c21508_g1_i1 | 80 | 4.00E-15 | 37 |
| | c19000_g3_i1 | 71 | 2.00E-12 | 37 |
| | c19000_g3_i2 | 52 | 6.00E-07 | 33 |
| | c35003_g1_i1 | 47 | 2.00E-05 | 26 |
| DcVTE2-1 | c17212_g1_i1 | 514 | E-146 | 64 |
| | c15601_g1_i2 | 265 | 6E-71 | 41 |
| | c16042_g1_i1 | 244 | 1E-64 | 46 |
| | c17212_g1_i2 | 215 | 4E-56 | 81 |
| | c15601_g1_i1 | 186 | 3E-47 | 42 |
| | c14187_g1_i1 | 164 | 9E-41 | 34 |
| | c22985_g1_i1 | 156 | 2E-38 | 39 |
| | c13877_g1_i2 | 152 | 6E-37 | 41 |
| | c13877_g1_i1 | 151 | 1E-36 | 41 |
| | c17212_g1_i3 | 147 | 1E-35 | 75 |
| | c13428_g1_i2 | 126 | 3E-29 | 44 |
| | c28572_g1_i1 | 120 | 1E-27 | 41 |
| | c32272_g1_i1 | 94 | 2E-19 | 34 |
| | c13428_g1_i1 | 92 | 6E-19 | 38 |
| | c21508_g1_i1 | 90 | 3E-18 | 40 |
| | c19000_g3_i1 | 85 | 9E-17 | 50 |
| | c19000_g3_i2 | 64 | 2E-10 | 44 |
| | c35003_g1_i1 | 49 | 0.000007 | 36 |
| | c12976_g1_i1 | 40 | 0.004 | 26 |
| | c18955_g3_i2 | 32 | 0.7 | 25 |
| | c18955_g3_i1 | 32 | 0.7 | 25 |
| CpPT1 | c13877_g1_i2 | 329 | 3.00E-90 | 97 |
| | c13877_g1_i1 | 328 | 6.00E-90 | 97 |

**Supplementary Table 7 Amino acid identities between bacterial and plant UbiA *O*-PTs for aromatics**

| | Bacteria | | Plant | | | |
|---|---|---|---|---|---|---|
| | CnqPT1 | AgqD | AkPT1 | CmiPT1a | CmiPT1b | CpPT1 |
| CnqPT1 | 100 | 16 | 13 | 13 | 13 | 13 |
| AgqD | 16 | 100 | 13 | 16 | 16 | 16 |
| AkPT1 | 13 | 13 | 100 | 42 | 42 | 42 |
| CmiPT1a | 13 | 16 | 42 | 100 | 100 | 99 |
| CmiPT1b | 13 | 16 | 42 | 100 | 100 | 99 |
| CpPT1 | 13 | 16 | 42 | 99 | 99 | 100 |

0% 100%

### Supplementary Table 8 List of PCR primers used in this study

| Primer name | Sequence (5’–3') |
|---|---|
| CpPT1_Fw | GCACAGTGAAGAAGAAGGAA |
| CpPT1_Rv | AACAAGTTCTCCTCATTTCG |
| CpPT1_TOPO_Fw1 | CACCATGCTGCTTCAAATGAATTT |
| CpPT1_TOPO_Rv | TCAGCGTACAAAGTGCATAA |
| CpPT2_Fw | GGATCCCATATGCTTCAAATGCAGTGCTTAAACCTCGCTC |
| CpPT2_Rv | GTCGACTCAGCGTAGAAAATGGATAAGGATGTATTC |
| CpPT3_Fw | GGATCCCATATGATGCTTCAAATGCATTCGAGTTCAAGC |
| CpPT3_Rv | CTCGAGTCATCGTAAGAAGTGAATAAGGAGATATTC |
| CpPT3_BamHI_Fw | GTGGATCCATGATGCTTCAAATGCATTCG |
| CpPT3_XhoI_Rv | TCTCGAGTCATCGTAAGAAGTGAATAAGGAGATA |
| CmiPT1_Fw | TCAGTCGACTGGATCATGCTGCTTCAAATGAATTTG |
| CmiPT1_Rv | GTCTAGATATCTCGATCAGCGTACAAAGTGCATAA |
| AkPT_DGP1_Fw | GCNHTNTTYAARGAYATHCCNGA |
| AkPT_DGP1_Rv | GCRTANARNARBTTCCADATRAA |
| AkPT_DGP2_Rv | ARBTTCCADATRAANMTRTARAA |
| AkPT1_3'RACE_Fw | CCCAGACTTATGTGCTCGGCAGACCAT |
| AkPT1_5'RACE_Rv | GCACCGTGACCCAACACTGTTATTAAC |
| AkPT1_Fw | AGAAGCCATCACCGATAAGC |
| AkPT1_Rv | CTCAACGTACATATCCCTCATCT |
| AkPT1_TOPO_Fw | CACCATGATACTGATACTGACTTCG |
| AkPT1_TOPO_Rv | TCAACGTAGAAAATGAACAAGTAG |
| CpPT1_qPCR_Fw | GGGTTATGCTTCGATCTCCA |
| CpPT1_qPCR_Rv | CATCAATGGACTTGCCTTCC |
| CpEF1α_Fw | TGGTGATGCTGGGTTTGTTA |
| CpEF1α_Rv | CACGCTCTTGATGACTCCAA |
| CpPT1_TOPO_Fw2 | CACCATGCTGCTTCAAATGAATTTG |
| CpPT1_TP210_Rv | CATCATATCCTCATTGTTTCC |
| CpPT1_woStop_Rv | GCGTACAAAGTGCATAAGGAAG |
| CpPT3_TOPO_Fw | CACCATGATGCTTCAAATGCATTCG |
| CpPT3_TP165_Rv | ATCACTTTGTGTGCATTTGATAGCG |
| AkPT1_TP180_Rv | GTAACGGTAGCTGTTTCCAAAA |
| citron_Fw | TCGACGAGCTTTCTGGATTT |
| citron_Rv | CAATTGGTTTATGGCAACCAC |
